# Development of compact transcriptional effectors using high-throughput measurements in diverse contexts

**DOI:** 10.1101/2023.05.12.540558

**Authors:** Josh Tycko, Mike V. Van, Aradhana, Nicole DelRosso, David Yao, Xiaoshu Xu, Connor Ludwig, Kaitlyn Spees, Katherine Liu, Gaelen T Hess, Mingxin Gu, Adi Xiyal Mukund, Peter H. Suzuki, Roarke A. Kamber, Lei S. Qi, Lacramioara Bintu, Michael C. Bassik

## Abstract

Human nuclear proteins contain >1000 transcriptional effector domains that can activate or repress transcription of target genes. We lack a systematic understanding of which effector domains regulate transcription robustly across genomic, cell-type, and DNA-binding domain (DBD) contexts. Here, we developed dCas9-mediated high-throughput recruitment (HT-recruit), a pooled screening method for quantifying effector function at endogenous targets, and tested effector function for a library containing 5092 nuclear protein Pfam domains across varied contexts. We find many effectors depend on target and DBD contexts, such as HLH domains that can act as either activators or repressors. We then confirm these findings and further map context dependencies of effectors drawn from unannotated protein regions using a larger library containing 114,288 sequences tiling chromatin regulators and transcription factors. To enable efficient perturbations, we select effectors that are potent in diverse contexts, and engineer (1) improved ZNF705 KRAB CRISPRi tools to silence promoters and enhancers, and (2) a compact human activator combination NFZ for better CRISPRa and inducible circuit delivery. Together, this effector-by-context functional map reveals context-dependence across human effectors and guides effector selection for robustly manipulating transcription.

## Introduction

Over 10% of human genes encode proteins that localize to the nucleus and function to regulate gene expression at the level of mRNA transcription (Göös et al., 2022; Lambert et al., 2018; Medvedeva et al., 2015). These transcription factors and chromatin regulator proteins represent a reservoir of potentially useful effector domains for constructing transcriptional and epigenomic perturbation tools for synthetic biology and dissecting biological processes (Gao et al., 2016; Gilbert et al., 2013; Mali et al., 2013; Perez-Pinera et al., 2013; Sanson et al., 2018). Transcriptional effector domains, also known as activators and repressors, are the regions of these proteins that can increase or silence transcription of a gene upon recruitment to its promoter (Lambert et al., 2018; Soto et al., 2021). When fused to DNA binding domains (DBD), they can be used as tools to manipulate the expression of endogenous genes and lncRNAs (Beerli et al., 1998; Gilbert et al., 2013; Joung et al., 2017), and control synthetic gene circuits (Rivera et al., 1996) or endogenous regulatory elements (Fulco et al., 2016; Hilton et al., 2015), which has therapeutic potential (Bailus et al., 2016; Segal et al., 2004; Thakore et al., 2018). However, transcriptional effectors can function differently depending on the biological context of their recruitment to a gene. For example, targeting of effector domains to the same genomic context but in different cell lines (Amabile et al., 2016; Hathaway et al., 2012; O’Geen et al., 2017), or in different mouse developmental stages *in vivo* (Ying et al., 2015), can result in variable transcriptional and epigenetic effects. The target locus of effector recruitment can also play a role (Cano-Rodriguez et al., 2016; Hong & Cohen, 2022; O’Geen et al., 2022; Sahu et al., 2022), wherein recruitment of effector domains to different promoters and enhancers in the same cell type results in distinct effects on transcription (Amabile et al., 2016; Hilton et al., 2015; Kearns et al., 2015; Nuñez et al., 2021; O’Geen et al., 2017, 2019). Finally, the DBD can affect effector function (Amabile et al., 2016; Cano-Rodriguez & Rots, 2016).

We currently lack a systems-level understanding of how frequently effector function depends on target, cell-type, or DBD context, which is critical for identifying effectors that are robust across these conditions. Recently, pooled screening methods that measure effector strength for large libraries of domains have greatly expanded the list of sequences known to function as effectors (Alerasool et al., 2022; Arnold et al., 2018; DelRosso et al., 2023; Sanborn et al., 2021; Tycko et al., 2020), but these scalable methods use synthetic reporter genes instead of targeting endogenous loci and have not yet been used to compare activities across many contexts. Meanwhile, other salient features like deliverability, size, expression level, and off-target toxicities remain uncharacterized for the majority of effectors.

To address these questions, we performed high-throughput recruitment (HT-recruit) to measure effector function of 5092 Pfam-annotated domains from human nuclear proteins (the Pfam library) across 17 biological contexts and integrated this data with 2 pre-existing datasets (Tycko et al., 2020). By adapting HT-recruit to use the programmable DBD dCas9, we were able to target endogenous promoters and enhancers. Using these large-scale datasets, we identify forms of context-specificity, which varied by effector family and were striking in the HLH family of domains. We then confirmed and expanded these results with screens of 114,288 tiling sequences from all human chromatin regulators and transcription factors (the CRTF tiling library) across 2 endogenous gene contexts, which we integrated with 3 pre-existing reporter gene datasets (DelRosso et al., 2023). Finally, with additional measurements of effector expression level, deliverability, and cellular toxicity, we developed improved effectors for CRISPRi/a and synthetic TF circuits, including a stronger ZNF705 KRAB repressor and a short potent activator combination we called NFZ.

## Results

### HT-recruit quantifies transcriptional effector function across DNA-binding domain, cell type, and target gene contexts

We set out to measure transcriptional effectors quantitatively across biological contexts by performing high-throughput recruitment (HT-recruit) screens using two DBDs (rTetR and dCas9) to recruit effectors to various gene targets in two cell-types (**Figure 1A and Supplementary Figure 1A**). First, to study different promoter contexts while keeping other parameters constant, we developed a set of reporters with varied promoter origins and strengths (**Figure 1B**). To study how effector-promoter interactions differ across cell types, we installed these reporters at the AAVS1 safe harbor in both K562 and HEK293T cells (**Figure 1A**). As expected, all minimal promoters (minCMV, NTX, and NT21) started OFF, and could be activated with the VP64 activator domain (**Figure 1B**). Two of the constitutive promoters, pEF1α and UbC, started ON and could be repressed by the ZNF10 KRAB domain (**Figure 1B**). The RSV promoter was rapidly silenced upon installation in both cell types (**Supplementary Figure 1B**), so we did not use it for screens. In certain cases we observed promoter silencing to be cell-type specific: the PGK promoter was constitutively ON in K562 cells, but was background silenced in HEK293T cells (**Supplementary Figure 1A**). Based on this observation, we decided to use PGK as a repressible promoter in K562 and as an activatable promoter in HEK293T cells (**Figure 1B**). Overall, the expression levels for each promoter were similar across the two cell types (R^2^=0.86) except for the PGK promoter (**Supplementary Figure 1C**).

**Figure 1.**
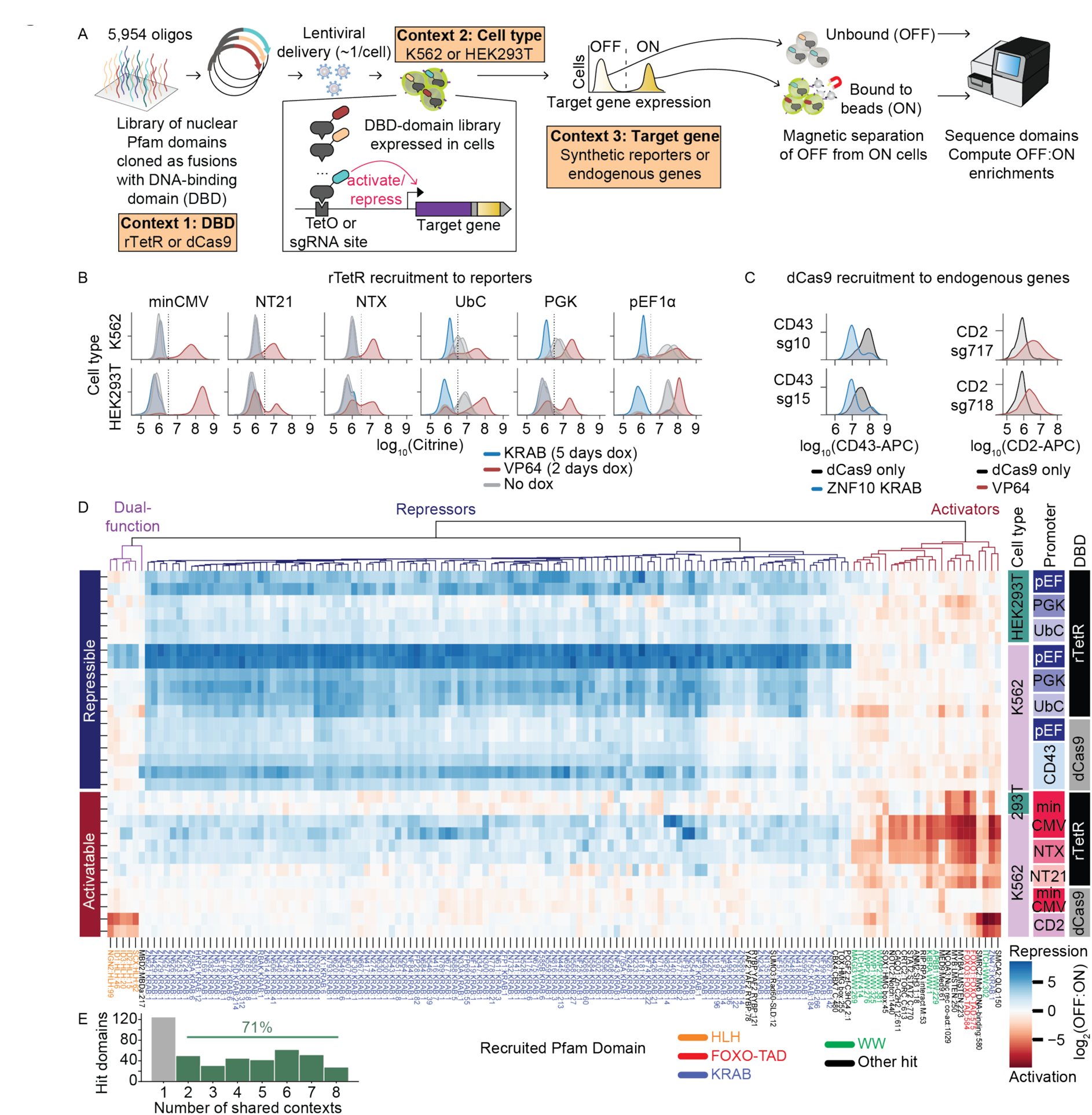
HT-recruit quantifies transcriptional effector functions across DNA-binding domain, cell-type, and target gene contexts. **A**) Schematic of high-throughput recruitment (HT-recruit) to quantify transcriptional effector function while varying the context of DNA-binding domains (DBDs), cell type, and target synthetic reporters or endogenous genes. A pooled library of Pfam domains from human nuclear proteins of ≤80 amino acids is synthesized as 300-mer DNA oligonucleotides, cloned downstream of the doxycycline (dox)-inducible rTetR DNA-binding domain (DBD) or dCas9 (Context 1), and delivered to either K562 or HEK293T cells (Context 2) at a low multiplicity of infection (MOI) such that the majority of cells express a single DBD-domain fusion. The target gene (inset) can be silenced or activated by recruitment of repressor or activator domains to the promoter. The synthetic reporters are integrated in the AAVS1 safe harbor and can be driven by different promoters (Context 3) and encode a synthetic surface marker (Igκ-hIgG1-Fc-PDGFRβ, purple) and fluorescent marker (Citrine, yellow). The endogenous target genes encode for surface markers (Context 3). After recruitment of Pfam domains, ON and OFF cells were magnetically separated using beads that bind these synthetic or endogenous surface markers, and the domains were sequenced in the Bound and Unbound populations to compute enrichments. **B)** Expression of synthetic reporters in K562 and HEK293T cells. The minimal reporter promoters, expected to be activatable, are minCMV, NTX, and NT21 and the stronger promoters, expected to be repressible, are pEF1α, PGK, UbC, and RSV. Positive control effectors, ZNF10 KRAB repressor or VP64 activator, were stably delivered by lentivirus. Cells were treated with 1000 ng/mL doxycycline for 5 days for repression and 2 days for activation (or untreated as a negative control) and Citrine expression was measured by flow cytometry after gating for rTetR delivery (mCherry^+^) (n=2 infection replicates shown as curves). **C)** Expression of endogenous surface marker genes CD2 and CD43 in K562 cells measured by immunostaining and flow cytometry. dCas9 fusions and sgRNAs were delivered by lentivirus and selected for by blasticidin and puromycin, respectively. Data are gated for sgRNA delivery (mCherry^+^ in CD43 and GFP^+^ in CD2 samples) and for dCas9 (BFP^+^) (n=1 infection replicate). **D)** Clustered heatmaps of transcriptional effector hits’ activation and silencing activity across different target gene, DBD, and cell-type recruitment contexts. To visualize a set of strong hits, a subset of effectors (columns) are shown that are a hit at a high threshold of 3 standard deviations stronger than the median of the poorly-expressed domains in ≥2 samples (rows) (n=143). Unbiased column clustering shows 3 major clusters of effectors that can be repressors, activators, or either depending on context (top). dCas9 targets pEF1α and minCMV with sgTetO-1, CD2 with sg717, and CD43 with sg10 (upper 2 rows) and sg15. Column labels on bottom show the protein, Pfam domain, and domain start position within the protein; select Pfam domain families are colored. Rows are manually ordered, with the targets that are predominantly repressible (strong reporters and CD43) above, and the predominantly activatable targets (minimal reporters and CD2) below. **E)** Distribution of the number of screen contexts in which a Pfam domain was a hit effector in two replicates. The percentage of domains that are shared hits in multiple contexts is colored.

To extend HT-recruit to endogenous loci, we used dCas9 and targeted the promoters of endogenous genes encoding cell surface proteins. Targeting surface proteins allows us to use fluorescent antibodies to immunostain cells, thus providing a way to monitor single-cell gene expression variability during individual recruitment assays by flow cytometry and to magnetically separate a large number of ON and OFF cells during HT-recruit (**Figure 1A**). For a repressible context, we targeted the highly expressed surface marker CD43 in K562 cells. First, we individually recruited either dCas9 alone or dCas9-KRAB with sgRNAs targeting the CD43 transcriptional start site (TSS) and found two sgRNAs, sg10 and sg15, for which repression depended on the KRAB repressor. Similarly, we identified sgRNAs with which dCas9-VP64 or VPR could activate the lowly expressed CD2, CD20, and CD28 genes (**Figure 1C and Supplementary Figure 1D,E**).

Then, we generated lentiviral libraries with 5092 Pfam domains from nuclear-localized human proteins and 499 random and 362 negative control sequences tiling the non-nuclear protein DMD (hereafter Pfam library) fused to rTetR or dCas9 and delivered them to cell lines with a low multiplicity of infection (MOI) such that most cells express one DBD-domain fusion. Using these components, we performed 8 rTetR-targeted screens across 6 synthetic reporter contexts in two cell types and 11 dCas9-targeted screens with sgRNAs targeting endogenous genes plus 2 dCas9 screens at reporters using an sgRNA targeting the TetO motif. We also integrated data from 2 previous screens using rTetR and the Pfam library (**Figure 1D and Supplementary Figure 1F,G**). To measure the activities of sequences beyond annotated Pfam domains, we also used our recently designed 20x larger library that tiles all human chromatin regulators and transcription factors in 80 amino acid tiles at 10 AA steps (n=128,565 elements, hereafter CRTF tiling library) (DelRosso et al., 2023), and recruited it with dCas9 to CD2 and CD43 in K562 cells (**Supplementary Figure 2A,B**). We additionally integrated data from 3 previously performed rTetR screens with the CRTF library at pEF, PGK, and minCMV (DelRosso et al., 2023).

To minimize confounding effects due to protein stability, we identified domains that were well-expressed in both cell types and when fused to either DBD. Expression was measured by permeabilizing the cells, staining with an anti-FLAG antibody, sorting into high and low protein expression level bins, and then sequencing the domains in those two cell populations (R^2^=0.44-0.78 across comparisons of replicates, **Supplementary Figure 3A**). Overall, domain expression was similarly correlated between cell types (R^2^=0.6, **Supplementary Figure 3B**) and whether they were fused to rTetR or dCas9 (R^2^=0.56, **Supplementary Figure 3C,D**). We labeled a domain as well-expressed if its score was 1σ above the median of the random controls. Meanwhile, the poorly-expressed domains, which included most random sequence controls, served as a large set of negative controls for activation and repression screens in order to set our hit threshold.

After filtering out poorly-expressed domains, we conservatively called activation and repression hit domains with log2(OFF:ON) scores 2 standard deviations beyond the median of poorly-expressed elements (higher scores for repressors, lower for activators). This pipeline called 435 Pfam domains as hits in both replicates of a context, with 71% of these hits being re-discovered in multiple contexts (**Figure 1E**). We also called 1,261 tiled regions as hit domains from the larger CRTF tiling library (**Methods**). The hits clustered into 3 categories: activators, repressors, and dual-function effectors that are context-dependent activators or repressors (**Figure 1D**). We then set out to investigate the diverse categories of context-dependent and context-robust effectors.

### Activator function across targets and cell types

We first considered activator promoter-specificity when using rTetR to target the Pfam library at minimal promoter reporters in K562 cells. We found that these core promoters responded very similarly to activator recruitment, wherein 85% of well-expressed NTX activators also activate minCMV, and do so with well-correlated strengths (R^2^=0.8, **Figure 2A,B and Supplementary Figure 4A,B**), which we individually validated with flow cytometry. In contrast, only 9% of the well-expressed minCMV activators, such as FOXO3 TAD, were able to reactivate the background silenced PGK promoter in HEK293T cells (**Supplementary Figure 4C,D**).

**Figure 2.**
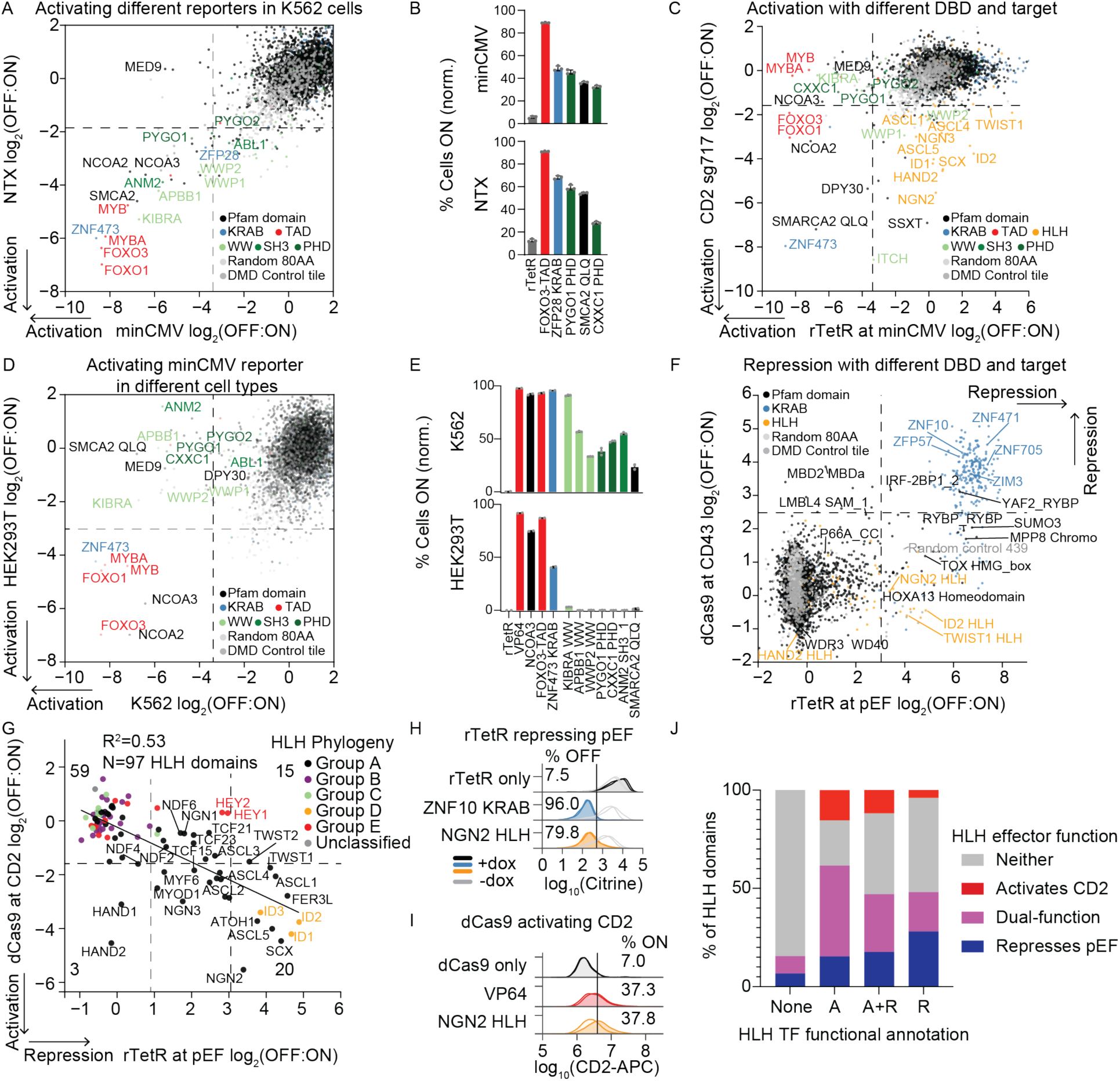
Context-dependent transcriptional effectors. **A**) HT-recruit with rTetR targeting two minimal promoters, minCMV and NTX in K562 cells (n=2 biological replicates). Dashed lines show hit thresholds at 2 standard deviations above the median of the poorly-expressed domains. Selected effectors are labeled with their gene name. **B)** Validation of activator domains across minimal promoter reporters in K562 cells. Individual rTetR-activator fusions or the rTetR-only negative control were delivered by lentivirus and, after selection, cells were treated with 1000 ng/ml doxycycline for 2 days to induce reporter activation. The percentage of cells activated was measured by flow cytometry for the Citrine reporter, after gating for delivery with mCherry. Bars show average percentage of cells ON normalized to no doxycycline control, error bars are standard deviation (n=3 infection replicates shown as dots). **C)** HT-recruit with dCas9 targeting endogenous gene CD2 with sg717 compared with rTetR targeting the minCMV reporter in K562 cells (n=2 biological replicates). Dashed lines show hit thresholds at 2 standard deviations above the median of the poorly-expressed domains. **D)** HT-recruit with rTetR targeting the minCMV reporter in K562 and HEK293T cells (n=2 biological replicates per cell type). Dashed lines show hit thresholds at 2 standard deviations above the median of the poorly-expressed domains. **E)** Individual rTetR fusions were delivered to minCMV reporter K562 or HEK293T cells by lentivirus, selected with blasticidin, and then recruitment was induced with 1000 ng/ml doxycycline for 2 days. Activation was measured by flow cytometry with a gate for rTetR delivery (mCherry^+^). Bars show average percentage of cells ON normalized to no doxycycline control, error bars are standard deviation (n=2-3 infection replicates shown as dots). **F)** HTrecruit with dCas9 targeting endogenous gene CD43 with sg15 compared with rTetR targeting the pEF1α reporter in K562 cells (n=2 biological replicates). Dashed lines show hit thresholds at 2 standard deviations above the median of the poorlyexpressed domains. **G)** HT-recruit with dCas9 to activate the endogenous gene CD2 using sg717 compared with repression of pEF1α promoter with rTetR in K562 cells, showing only the HLH domains within the Pfam library (n=2 replicates per screen). Black line shows best linear fit. The conservative hit threshold (black dashed line) was chosen to identify robust effectors; some sub-threshold domains can have weaker repressor activity. The gray dashed vertical line is equivalent to the strength of the weakest repressor that was individually validated (**Methods**). The HLH phylogenetic groups are shown as colors (Atchley & Fitch, 1997; Ledent et al., 2002; Torres-Machorro, 2021). **H)** rTetR recruitment to the pEF1α reporter in K562 cells. Shaded distributions show cells after 6 days of treatment with 1000 ng/ml of doxycycline while light gray curves show the untreated cells (n=2 infection replicates). **I)** dCas9 recruitment to CD2 in K562 cells (n=2 sgRNAs, sg717 in darker shade and sg718 in lighter shade). **J)** Full-length HLH TFs were defined as Activators (A, n=13), Repressors (R, n=25), both (A+R, n=17) or not yet defined (None, n=45) in previous studies, which were reviewed in (Torres-Machorro, 2021). Colors show the fraction of HLH domains from the TFs within these categories that activate CD2 with dCas9, repress pEF1α with rTetR, or are dual-functioning.

To extend these experiments to endogenous gene targets, we used dCas9 recruitment. First, we targeted the minCMV reporter using a TetO-targeting sgRNA. dCas9 recruitment resulted in fewer hits than rTetR at the same reporter, perhaps due to weaker and/or lower copy number recruitment; however the strongest hit, the Med9 mediator component, validated individually (**Supplementary Figure 4E–G**). In contrast, when we used dCas9 to target the Pfam library to endogenous genes, we found 68 hits targeting CD2 with sg717, which systematically led to stronger activation than any other sgRNA targeting CD2, CD20, or CD28 (**Figure 2C and Supplementary Figure 4H–L**). Only 19% of rTetR activator hits at minCMV recurred as dCas9 hits at CD2 (e.g. ZNF473 KRAB, SMARCA2 QLQ); conversely, 89% of domains capable of activating CD2 were not hits in the rTetR-and reporter-based screens, including 24 HLH domains (e.g. NGN2, HAND2). The second strongest HLH activator domain was from HAND2, consistent with a recent report that full-length HAND2 can be an activator (Alerasool et al., 2022). Meanwhile, the two MYB LMSTEN domains were only activators when recruited with rTetR at reporters. Using the larger CRTF tiling library, dCas9 recruitment to CD2 identified 19 activator domains that were not hits with rTetR at minCMV. 42% of the CD2-only activators again overlapped with HLH domains (e.g. in MYOD1, PTF1A, MAX, ASCL5) while others were novel activators that did not overlap any domain annotation (e.g. PBRM1) (**Supplementary Figure 5A–D**). A notably strong shared activator hit with both rTetR at minCMV and dCas9 at CD2 was the DUX4 C-terminus, which interacts with histone acetyltransferase P300 (Choi et al., 2016). Therefore, rTetR recruitment to different minimal promoters identified similar activators, while dCas9 recruitment to an endogenous gene identified a largely distinct set of activators including HLH domains.

We then investigated the cell-type dependence of different activators by comparing their effects in HEK293T versus K562 cells when recruited by rTetR at the minCMV reporter. In general, we found much greater differences between activators targeted to the same promoter in different cell types than when targeting distinct minimal promoters within the same cell type. Overall, only 19% of activators in K562 cells were shared across the cell types, including strong activators like FOXO-TADs, NCOAs, and the ZNF473 KRAB domain (**Figure 2D,E and Supplementary Figure 6A**). Meanwhile, several WW, SH3, PHD and the poorly characterized SMARCA2/4 QLQ domains (Alfert et al., 2019; Treich et al., 1995) were much stronger activators in K562 cells, despite being similarly expressed in the two cell types as measured by FLAG staining (**Figure 2E, Supplementary Figure 3B, and Supplementary Figure 6B–D**). We hypothesized that one source of cell type dependence of an effector could arise from competition for coactivators with its endogenous copy, and found that knocking down the endogenous SMARCA2/4 resulted in increased activation by the recruited SMARCA2 QLQ (from 1.3% to 23.7% of HEK293T cells ON) (**Figure 2D,E and Supplementary Figure 6B–H**).

Overall, while we cannot rule out all technical reasons (e.g. related to the reporter chromatin state or rTetR) that these activators are stronger in K562 cells, these results show that, controlling for DBD and target sequence, some activators function very differently across cell types (e.g. KIBRA WW activates 91% of K562 vs 3% of HEK293T cells) while others are more consistent (e.g. FOXO3-TAD activates 93% of K562 vs 87% of HEK293T cells).

### Repressor function across targets and cell types

We next tested repressor promoter-specificity using rTetR to target pEF1α, PGK, and UbC promoters in K562 cells. We found repression scores at the moderate strength PGK and UbC were highly correlated with each other, largely due to signal from KRAB repressors (R^2^=0.74 for n=2718 well-expressed Pfam domains, **Supplementary Figure 7A**), whereas the stronger pEF1α was more silenced by weaker repressors such as HOX homeodomains (**Supplementary Figure 7B,C**). We previously observed that HOX homeodomain repression strength correlated with the presence of an RKKR motif and positive charge in the homeodomain N-terminal region (Tycko et al., 2020), so we recruited motif-and charge-modifying HOX mutants to the pEF1α reporter and confirmed they contribute to repression (**Supplementary Figure 7D,E**). In sum, the similarly strong PGK and UbC reporters responded similarly to repressors, whereas the stronger pEF1α reporter uniquely identified weak non-KRAB repressors and thus could be used to characterize their sequence dependencies.

To identify repressors of endogenous genes, we used dCas9 recruitment. When targeting the pEF1α reporter with dCas9, we found fewer hits than with rTetR (possibly due to lower recruitment copy number) and all the hits were KRAB repressors (**Supplementary Figure 7F–H**). In contrast, targeting dCas9 to the endogenous CD43, we found that in addition to KRAB domains, a distinct set of domains showed repressor activity, including the NuRD-interacting Methyl-binding protein domains such as the P66_CC domain from P66A and the MBDa domain from MBD2 (which bind one another (Gnanapragasam et al., 2011)), that were not repressors in the rTetR screens (**Figure 2F and Supplementary Figure 7I–K**). Relatedly, an MBD2B repressor was previously shown to silence a reporter in HEK293T cells with dCas9 recruitment (Yeo et al., 2018). Other repressor hits unique to targeting CD43 with dCas9 were a zinc finger from the IRF-2BP1 co-repressor protein (Childs & Goodbourn, 2003) and the SAM1 domain from the putative polycomb protein L3MBTL4 (whose analog is involved in silencing in Drosophila cells (A. K. Robinson et al., 2012)). MBDa, P66_CC, IRF-2BP1 and SAM1 silencing of CD43 depended on which sgRNA was used while KRAB repressors were efficient with either of two sgRNAs (**Supplementary Figure 7I,J**). Meanwhile, Chromo and Chromoshadow domains were only hits with rTetR.

dCas9 recruitment of the CRTF tiling library to the CD43 promoter confirmed these results, with the strongest shared repressors being KRAB domains, and additionally revealing 274 repressor domains that were not hits at pEF1α including from additional Methyl-binding domain proteins (**Supplementary Figure 8A–C**). This larger library also uncovered repressors in the unannotated regions of proteins, including CDCA7L, which is known to be a repressive protein (K. Chen et al., 2005). 78% of CD43 repressor domains were also hits in the rTetR screens and they overlapped similar annotations for SUMOylation, short linear interactions motifs, and DNA-binding domains (DelRosso et al., 2023) (**Supplementary Figure 8D**).

We then investigated the cell type-specificity of repressors using rTetR to target the Pfam library at pEF1α reporters. The HEK293T screen identified strong repressors which were in agreement across cell types, such that 96% of repressor hits in HEK293T cells were also hits in K562 cells (including >200 KRAB domains, **Supplementary Figure 9A–D**). When using the moderately strong UbC promoter, we found fewer hits than with pEF1α, and again the majority were shared across cell types (e.g. >130 KRAB domains) (**Supplementary Figure 9E**). Altogether, these results show that some repressors (e.g. MDBa) strongly depend on recruitment context while KRAB repressors are particularly robust across cell-type, target, and DBD contexts.

### Context-specific HLH effector domains activate some promoters and repress others

Effector domains are classified as activators or repressors; however, like the transcription factors that consist of these domains (Soto et al., 2021), these effectors may have a dual activator-repressor function (DelRosso et al., 2023) that could be context-dependent. When comparing most activatable and repressible contexts, we saw no overlap in hits; however, when considering the most sensitive repressible context (rTetR at pEF1α in K562 cells) we found some activators could repress (**Supplementary Figure 10A–D**). Certain minCMV strong activators either silence a percentage of the pEF1α cells (i.e. bifurcate the population) or super-activate it in a cell type-dependent manner (**Supplementary Figure 10E,F**).

In addition, the large HLH family, which includes transcription factors shown to function either as an activator or repressor depending on context in yeast (McIsaac et al., 2012), stood out as a major source of dual-functioning effectors in our data. Specifically, many of the same HLH effectors that activate CD2 when recruited with dCas9 also repressed pEF1α when recruited with rTetR with correlated strength (R^2^=0.53, n=97 HLH domains) (**Figure 2G-I**). Furthermore, we found that effector function is enriched within certain HLH phylogenetic groups (Atchley & Fitch, 1997; Ledent et al., 2002; Torres-Machorro, 2021); for example, HAND HLHs from Group A are only activators, HEY HLHs from Group E are only repressive, ID HLHs from Group D are dual-functioning, and other subfamilies (e.g. HES from Group E and all of Group B) did neither in the contexts tested here (**Figure 2G and Supplementary Figure 11A**). These HLH effector functions partially corresponded with annotations from previous studies of the HLH transcription factors as activators, repressors, or both (**Figure 2J**) (Torres-Machorro, 2021). These results were confirmed with the tiling library, where 74% of proteins with a dual-function tile that activates CD2 (but not minCMV) and represses pEF1α were again HLH proteins. The hit tiles overlapped the HLH heterodimerization portion of their bHLH domains (**Supplementary Figure 11B–D**) (DelRosso et al., 2023).

Overall, while some effectors were highly context dependent (e.g. HLH and WW domains), we noticed many of the strongest effectors were consistent across contexts (e.g. KRAB repressors), which suggests they would improve synthetic tools for manipulating transcription.

### Efficient CRISPR interference with the ZNF705/ZNF471 KRAB repressors

We set out to identify repressors that are robust across contexts using the Pfam library because it was screened against the largest number of contexts. In addition, we performed an additional screen with this library at the *GATA1* locus to compare repressors at an enhancer (**Supplementary Figure 12A**). Since GATA1 is an essential gene, we used the growth phenotype associated with targeting dCas9-repressors to its enhancer (eGATA1) as a proxy for repression strength (Fulco et al., 2016; Tycko et al., 2019). The KRAB domains found in ZNF705B/D/F (here called ZNF705) and ZNF471 were the strongest hit Pfam domains with the eGATA1 sgRNA and they did not have growth effects with a control safe-targeting sgRNA (**Supplementary Figure 12B**). Meanwhile, some domains did show growth effects with the safe-targeting sgRNA, suggesting their expression is toxic. The most toxic were the cyclin-dependent kinase inhibitor (CDI) domains from CDKN1A/B, which inhibit cell cycle progression by binding to CDK2 (Russo et al., 1996). Interestingly, the random sequences were significantly more toxic than the Pfam domains or DMD tile controls (**Supplementary Figure 12C,D**). We confirmed these toxic domains were not confounding repressor hits when targeting the non-essential gene CD43 (**Supplementary Figure 12E**).

To identify strong repressors that function well across many different contexts, we then focused on the KRAB family, which were commonly top hits, and ranked 323 KRAB domains by their scores across all of the repressor screens. ZNF705 KRAB ranked as #18 and ZNF471 KRAB as #1 (**Figure 3A**). In agreement with recent findings, our approach also identified ZIM3 KRAB (#13) as a stronger repressor than ZNF10 KRAB (#166) (Alerasool et al., 2020; Replogle et al., 2022). Further supporting their context-robustness and reproducibility, we found ZNF705, ZNF471 and ZIM3 KRAB were hits in both replicates of 8, 7, and 6 of the original 8 repressive contexts, respectively (**Figure 1E**). We could not identify more efficient non-KRAB repressors (**Supplementary Text and Supplementary Figure 13A–H**), consistent with KRAB repressors being most potent across contexts in the screens.

**Figure 3.**
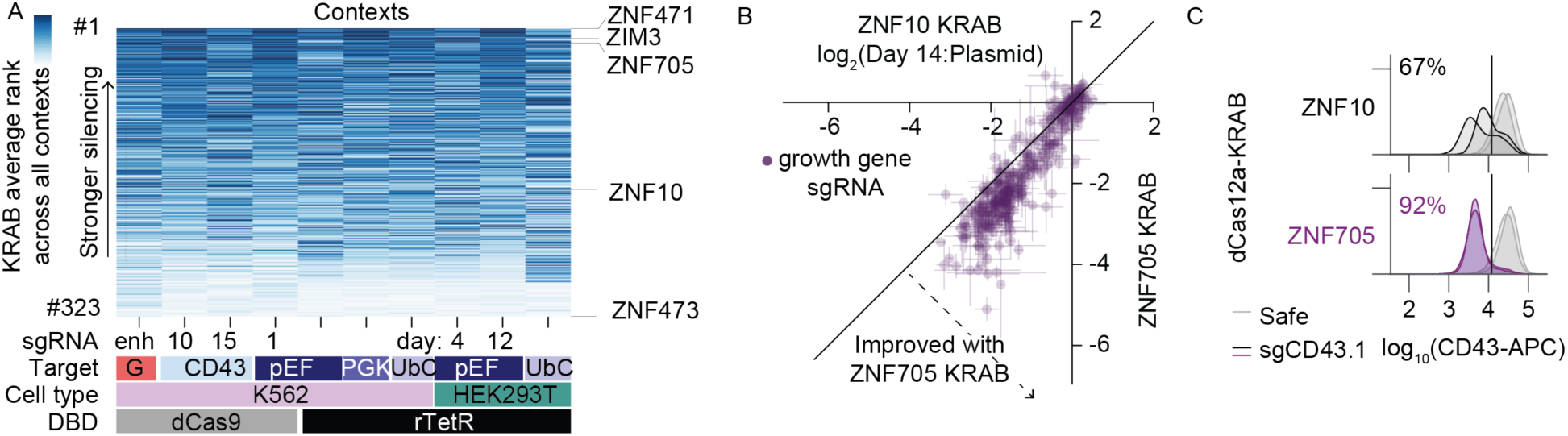
ZNF705 KRAB improves CRISPRi across target and DBD contexts. **A**) KRAB domains ranked by repression in HT-recruit screens across target, cell type, and DBD contexts (a higher average log_2_(OFF:ON) from 2 biological replicates is better ranked). Row order and color shows average rank across all screens. Select KRAB proteins are labeled on the right. The red target labeled G shows the growth-based screen at the GATA1 enhancer. The HEK293T pEF screen has two timepoints: a silencing measurement at 4 days after doxycycline addition, and a memory measurement at day 12 (8 days after doxycycline removal). The 323/336 KRAB domains that had no missing data across these screens are included. **B)** A CRISPRi screen was performed to compare KRAB repressors. An sgRNA library targeting 37 essential gene promoters was delivered into K562 cell lines that stably express either dCas9-ZNF10 KRAB or dCas9-ZNF705 KRAB, cells were passaged for 14 days, and then the guides were sequenced to measure fitness effects (shown as a log_2_ fold-change from the original plasmid pool to the final day 14 measurement from genomic DNA). Greater depletion is a measure of stronger silencing of the essential genes. Each dot shows the average effect for an sgRNA and the error bars show the S.D. from 2 screen replicates (n=405 sgRNAs). The diagonal line represents identity between KRAB domains. **C)** dCas12a recruitment of ZNF10 or ZNF705 KRAB with guide RNAs to target CD43. dCas12a fusions and then guide RNAs were delivered by lentivirus to K562 cells. 9 days after guide RNA infection, cells were stained for surface CD43 expression and analyzed by flow cytometry, with gates applied for guide RNA expression (mCherry) and dCas12a expression (HA-tag stain). Infection replicates are shown as separate histograms and their average percentage of silenced cells is shown (n=2 biological replicates shown as lines).

We set out to validate the top-ranked KRAB domains across different target contexts compared to a strong benchmark. Previously, using a deep mutational scan of the ZNF10 KRAB, we identified a mutant, WSR7EEE, in the N-terminal KRAB domain region that provides increased expression in cells and silencing strength when fused to rTetR (Tycko et al., 2020). We found the same is true when this mutant is fused to dCas9, so we used this enhanced ZNF10 KRAB[WSR7EEE] as a benchmark when determining how well the KRAB repressor paralogs work (**Supplementary Figure 14A–E**). Encouragingly, the KRAB domains from ZNF705 and ZNF471 both outperform the ZNF10 KRAB[WSR7EEE] at silencing CD43 (**Supplementary Figure 14F**). To test additional targets, we performed CRISPRi screens with a library including 405 sgRNAs targeting promoters of 37 essential genes (**Supplementary Figure 14G**) and found that the ZNF705 KRAB consistently provides ∼1.5x greater effect sizes across a range of ZNF10 baseline effects (**Figure 3B and Supplementary Figure 14H,I**). These KRABs completely silence a reporter in HEK293T cells (**Supplementary Figure 14J,K**) and, in separately described results, we found ZNF705 KRAB consistently exhibits stronger silencing at enhancers (Yao *et al*. 2023, *in review*).

To assess the KRAB repressors in a different DBD context, we used an engineered form of dAsCas12a (dEnAsCas12a, 1308 aa) (Kleinstiver et al., 2019; Tak et al., 2017). dCas12a is weaker than dCas9 for recruiting repressors, so there is more need to improve its efficiency (O’Geen et al., 2017). Fusing ZNF705 KRAB to dCas12a resulted in a mean 1.5x more complete silencing of a population of cells than ZNF10 KRAB, across multiple targets (**Figure 3C and Supplementary Figure 14L**). Encouragingly, a 99% identical ZNF705 family KRAB (from ZNF705E) was ranked #12 out of >1,000 repressor hit tiles recruited in 3 contexts with the larger tiling library, further establishing ZNF705 KRAB as a particularly strong repressor (**Supplementary Figure 14M,K**). Other highly-ranked KRABs likely perform similarly well.

### Compact activator combinations for lentiviral CRISPRa

We next ranked activators from HT-recruit measurements performed across target, DBD, and cell-type contexts, and selected context-robust domains that were a hit in at least 5 activation screens (**Figure 4A**). ZNF473 KRAB (Z) is an acidic domain that was a strong hit with both dCas9 recruitment to CD2 and most rTetR contexts. Meanwhile, NCOA3 (N) and FOXO3 TAD (F), which bind p300 (H. Chen et al., 1997; Karlsson et al., 2022; F. Wang et al., 2012), were rTetR hits and stronger than ZNF473 in HEK293T cells. These domains were well-expressed across DBDs and cell types, and there was no toxicity associated with their expression (**Supplementary Figure 3D and Supplementary Figure 12E**). We had previously found that the trimmed Pfam-annotated domains (Z=44 AA, N=48 AA, and F=41 AA) for these three activators performed equally to the 80 AA sequence centered on the domain with rTetR (Tycko et al., 2020), so we used the trimmed sequences to build activator tools (**Figure 4B**). This trimmed FOXO3 sequence was within the #11 ranked activator tile in the larger library ranked by the average of the minCMV and CD2 screens.

**Figure 4.**
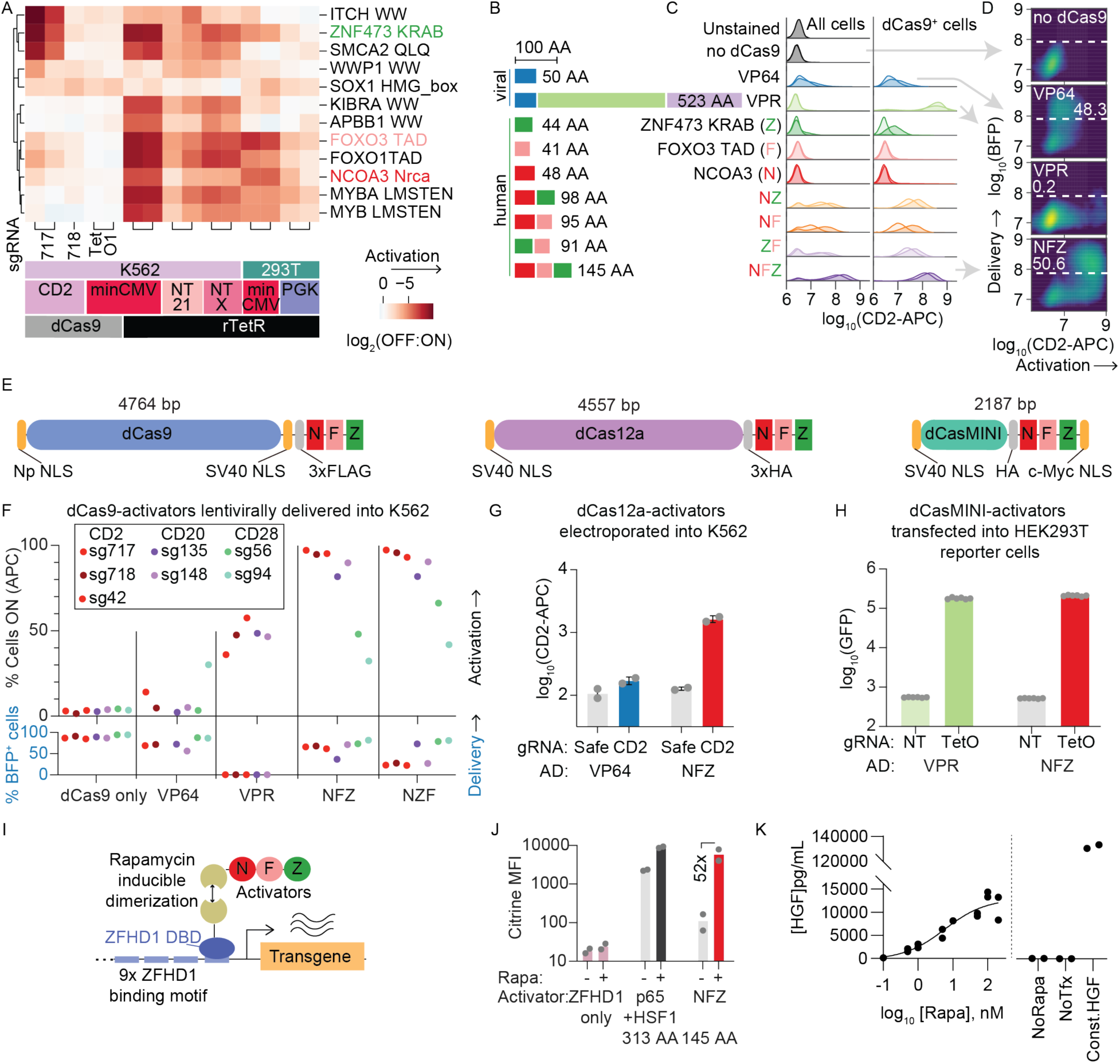
Compact NFZ activator improves CRISPRa and inducible systems. **A**) HT-recruit scores for activators that were hits in ≥5 samples across target, cell-type, and DBD contexts (n=2 replicates per rTetR screen and n=1-2 replicates per sgRNA for dCas9 screens shown as columns). The rows are clustered in an unbiased manner. Labels for NCOA3 [1045-1092] (N), FOXO3 FOXO-TAD [604-644] (F), and ZNF473 KRAB [5-48] (Z) are colored. **B)** Size of N, Z, and F human activator domains and combinations compared to VP64 and VPR viral activators. **C)** The effectors were recruited with dCas9 to activate the endogenous CD2 surface marker in K562 cells. sgRNAs were delivered by lentivirus and cells were selected with puromycin, then dCas9 fusions were delivered by 10x concentrated lentivirus. 4 days after dCas9 delivery, cells were stained for CD2 and expression was measured by flow cytometry. On the left, CD2 expression is shown after gating cells for sgRNA delivery (GFP) and, on the right, after gating for both sgRNA and dCas9 delivery (BFP). The darker shaded histogram is CD2 sg717 and the lighter shade is sg718. **D)** Comparison of CD2 activation and dCas9-activator delivery (BFP), after gating for CD2 sg717 delivery (GFP^+^). Color shows smoothed density of single cell flow cytometry data. The percentage of BFP^+^ cells is labeled (n=2 infection replicates). **E)** Schematics of CRISPR activator systems using the NFZ tripartite activator fused to dCas9, dEnAsCas12a, or dCasMINI_V4. Np NLS=nucleoplasmin nuclear localization signal. **F)** dCas9-activator fusions targeting CD2, CD20, CD28 surface marker genes in K562 cells. sgRNAs were first installed by lentiviral delivery and puromycin selection. dCas9-activators were delivered by 5x concentrated lentivirus, 5 days later blasticidin selection was started, and then 4 days later cells were stained and analyzed by flow cytometry. Top: the percentage of cells ON (APC) is shown after gating for the sgRNA (GFP^+^) and dCas9 (BFP^+^). Note all of the VPR samples had <250 BFP^+^ cells counted (associated with poor infection and survival of blasticidin selection), while all other samples had >6k BFP^+^ cells (mean=53k). Bottom: the percentage of BFP^+^ cells is shown. The CD28 sgRNAs infected with VPR poorly survived blasticidin selection and <300 cells total could be counted, so they are not shown. Each dot represents a different sgRNA targeting the gene (n=5-7 sgRNAs per effector). **G)** NFZ or VP64 were fused to dEnCas12a on a marker-less plasmid and co-electroporated into K562 cells with a CD2-targeting or safe-control gRNA expression plasmid with an mCherry marker. After 3 days, cells were stained for CD2 with APC-conjugated antibody and APC MFI was measured by flow cytometry after gating for high mCherry (n=2 replicates). **H)** NFZ or VPR were fused to dCasMINI on a plasmid with an mCherry marker and co-transfected into HEK293T GFP reporter cells with a TetO-targeting or non-targeting control gRNA expression plasmid with a BFP marker. 2 days later, GFP reporter activation was measured by flow cytometry with gates for BFP^+^/mCherry^+^ cells (n=6 transfection replicates shown as dots, bar shows mean). **I)** Schematic of the minimized rapamycin-inducible transcription cassette using the ZFHD1 DBD fused to NFZ. **J)** Rapamycin-inducible expression of citrine with ZFHD1 recruitment of NFZ or a control activator combination, p65+HSF1. 3 ug of plasmids were transfected into HEK293T cells, one day later 10 nM rapamycin was added, and then two days later citrine mean fluorescence intensity (MFI) was measured by flow cytometry (n=2 transfection replicates shown as dots, bar shows mean). **K)** Rapamycin-inducible expression of HGF with ZFHD1 recruitment of NFZ, after transfection in HEK293T cells. Secreted HGF protein concentrations were measured in the cell culture supernatant by ELISA after 2 days of rapamycin treatment with varied doses. Transfection of no plasmid and a constitutive pEF1α-HGF plasmid served as a negative and positive control, respectively (n=2 transfection replicates, curve shows nonlinear regression fit to dose response data).

We fused these trimmed activators, or two commonly used synthetic activators (VP64 and VPR) (Chavez et al., 2015; Maeder et al., 2013), individually to dCas9 in a lentiviral vector with BFP as a delivery marker. Given previous observations of synergistic activation from concatenated activators, including VP64 compared to VP16 and VPR compared to its components (Chavez et al., 2015; Dominguez et al., 2022; Konermann et al., 2015; Perez-Pinera et al., 2013; Tanenbaum et al., 2014; Zhou et al., 2018), we reasoned that fusing N, F, and Z could improve activation. Indeed, when we combined any of these domains together as bipartite activators, we observed synergistic activation of the CD2 gene (**Figure 4C**).

Further fusion of all three domains together to create the tripartite activator, NFZ, led to even stronger activation (**Figure 4C**). At 145 AA long, NFZ is more compact than the tripartite 523 AA VPR, and does not use viral components. We found that NFZ was more efficiently delivered by lentivirus, more efficiently generated stable cell lines after blasticidin selection, and provided a similar or higher level of activation across 3 target genes (**Figure 4C-F and Supplementary Figure 15A–C**). Further, we could use lentivirus to deliver NFZ, but not VPR, to J774 mouse macrophages, which are more difficult to infect than K562 cells, and successfully select a stable line in which we could activate endogenous genes (**Supplementary Figure 15D–F**). Additionally, by changing the N, F, Z domain orientation, we identified NZF as a moderately stronger activator with lentiviral delivery efficiencies between VPR and NFZ (**Figure 4F and Supplementary Figure 15A,B,G**).

NFZ may be more deliverable by lentivirus than VPR due to differences in transduction efficiency and/or cell toxicity related to the expression of VPR. While VPR is the largest effector, the lentiviral insert size (between cPPT and WPRE) is 8.5 kb, which does not exceed packaging limits (Counsell et al., 2017). Meanwhile, transcriptional squelching is associated with VP16-derived activators (Ptashne & Gann, 1990; Sadowski et al., 1988), and VPR can be toxic in cell lines (Jones et al., 2020), which could affect the packaging cells and/or transduced cells. When using transient plasmid delivery, we observed similar or better activation with VPR than NFZ (**Supplementary Figure 15H,I**), despite observing lower expression of VPR transcripts (**Supplementary Figure 15J,K**) (consistent with others’ observations of low dCas9-VPR expression (Mahata et al., 2022; K. Wang et al., 2022)), suggesting NFZ is specifically advantageous for viral delivery.

Notably, the tripartite fusions were strong activators even with dCas9 targeting CD20, CD28, CD2 (with its sgRNAs beyond sg717), or the minCMV reporter, which were all contexts with poor activation from nearly any single domain in the Pfam library (**Figure 4C, F, Supplementary Figure 15G–I, L–N**). We found no further improvement by fusing a fourth domain, QLQ, or using homotypic combinations (**Supplementary Figure 15L–O and Supplementary Text**).

### Fusing compact tripartite activator NFZ to compact CRISPR DBDs

dCas9 is a relatively large DBD (1368 aa) and it is difficult to multiplex with multiple sgRNAs. To make a further miniaturized CRISPRa system, we ported NFZ to dEnAsCas12a (1308 aa) (Guo et al., 2022; Kleinstiver et al., 2019), and found NFZ activated CD2 ∼10-fold more strongly than VP64 (**Figure 4E,G and Supplementary Figure 16A,B**). As with dCas9, we could not establish a stable dCas12a-VPR cell line using lentivirus (7.6 kb) and blasticidin selection, but we were successful with dCas12a-NFZ (**Supplementary Figure 16C**).

Recently, a very compact dCas12f DBD called dCasMINI (529 aa) was engineered as a CRISPRa system in human cells (Xu et al., 2021). We found dCasMINI-NFZ (2.2 kb) strongly activated HEK293T reporter cells (**Figure 4E,H and Supplementary Figure 17A–E**).

### Inducible expression with the compact tripartite activator NFZ

Another useful application of compact human activators is engineering circuits that tunably control transgene expression using a synthetic transcription factor that conditionally activates a promoter in the presence of a small molecule. Previously, the zinc finger-homeodomain DBD (ZFHD1) (Pomerantz et al., 1995) was fused to FKBP-dimerization domains that recruit HSF1 and P65 activators in the presence of rapamycin (Rivera et al., 2005). We designed a >1 kb smaller version of this design using NFZ (**Figure 4I and Supplementary Figure 18A**). Upon transfection, we observed rapamycin-dependent reporter transgene inducibility and low levels of leakiness with NFZ relative to P65+HSF1 (**Figure 4J and Supplementary Figure 18B–D**). Next, we used ZFHD1-NFZ to inducibly express the human hepatocyte growth factor (HGF) gene (2.2 kb) (Lee et al., 2019; Matsuda et al., 2021; Morishita et al., 2020; Schievenbusch et al., 2010; Suzumura et al., 2008), and observed a rapamycin dose-dependent increase in secreted HGF protein, with minimal leakiness in untreated cells (**Figure 4K**). This compact circuit could enable longer proteins (such as HGF) to be inducibly expressed from limited-size viral vectors like AAV.

## Discussion

A better understanding of transcriptional effector context-specificity is needed to improve strategies for manipulating transcription. Our systematic effector measurements across cell-type, genomic target, and DBD contexts revealed both context-dependent effectors (e.g. WW and HLH domains) and context-independent effectors that can improve transcriptional perturbation tools (e.g. the tripartite NFZ activator and the ZNF705 KRAB repressor). Further, our larger tiling library both confirmed context effects we originally observed with fewer elements in the Pfam library (e.g. in the HLH, KRAB, and MBD families) and provides a valuable resource of transcriptional effectors, including from unannotated protein regions, that function on dCas9 at endogenous genes. These transcriptional repressors and activators, in addition to others (Alerasool et al., 2020; DelRosso et al., 2023; Ludwig et al., 2022; Ma et al., 2018; Mahata et al., 2022; Mukund et al., 2022; Tycko et al., 2020; Vora et al., 2018; Yeo et al., 2018), can now be combined with varied DBDs (Li et al., 2022) to enable gene regulation tools that are more compact, humanized, and deliverable.

Looking beyond the context-robust effectors, this and other recent work (Jacobs et al., 2022; Policarpi et al., 2022) describe the extent of context-dependence across effectors. Overall, for both activators and repressors, differences across cell types were more likely to be changes in magnitude, whereas differences across target and DBD were more likely to be complete changes from effective to ineffective (or even from activating to repressing). More specific trends can be found within domain families such as the target-dependent HLH domains or the cell-type dependent WW and SH3 domains that interact with proline-rich motifs (Bedford et al., 1997; Macias et al., 2002) that are common in activators (Gerber et al., 1994). Integrating this effector function data with protein interaction maps could identify co-factors that help explain context-dependence.

Scalable synthetic systems can control for variability across contexts and isolate a single biological variable to measure, but they have technical limitations. For example, across effectors that function as activators, repressors, or both, we generally find stronger effects in K562 versus HEK293T cells and it seems possible that a global effect, for example on rTetR recruitment strength, contributes to this result. While these cell lines are useful for screening, further assessing effectors in a broader set of cell types including primary and stem cells will be important to uncover dependencies that are particularly relevant in disease and development. We also found dCas9 HT-recruit screens varied in their dynamic range depending on the sgRNA and target, and more measurements in endogenous target contexts are needed to test the relationship between chromatin state and effector function at scale. While 80 amino acid domain libraries are powerful due to their scalability, programmability, focus on short functional regions, and the deliverability of the compact domains, using longer sequences could uncover additional forms of context-dependence as longer sequences are more likely to encode multiple functions and layers of regulation.

Despite these limitations, our work highlights the potential of a high-throughput process for protein engineering in synthetic biology that is enabled by the ability to synthesize and test large pooled libraries of domains for function directly in human cells. In the future, HT-recruit could be adapted to systematically discover and characterize effectors for the other forms of gene and epigenetic editing, including base and prime editing (Velimirovic et al., 2022).

## Supplementary Figures

**Supplementary Figure 1.**
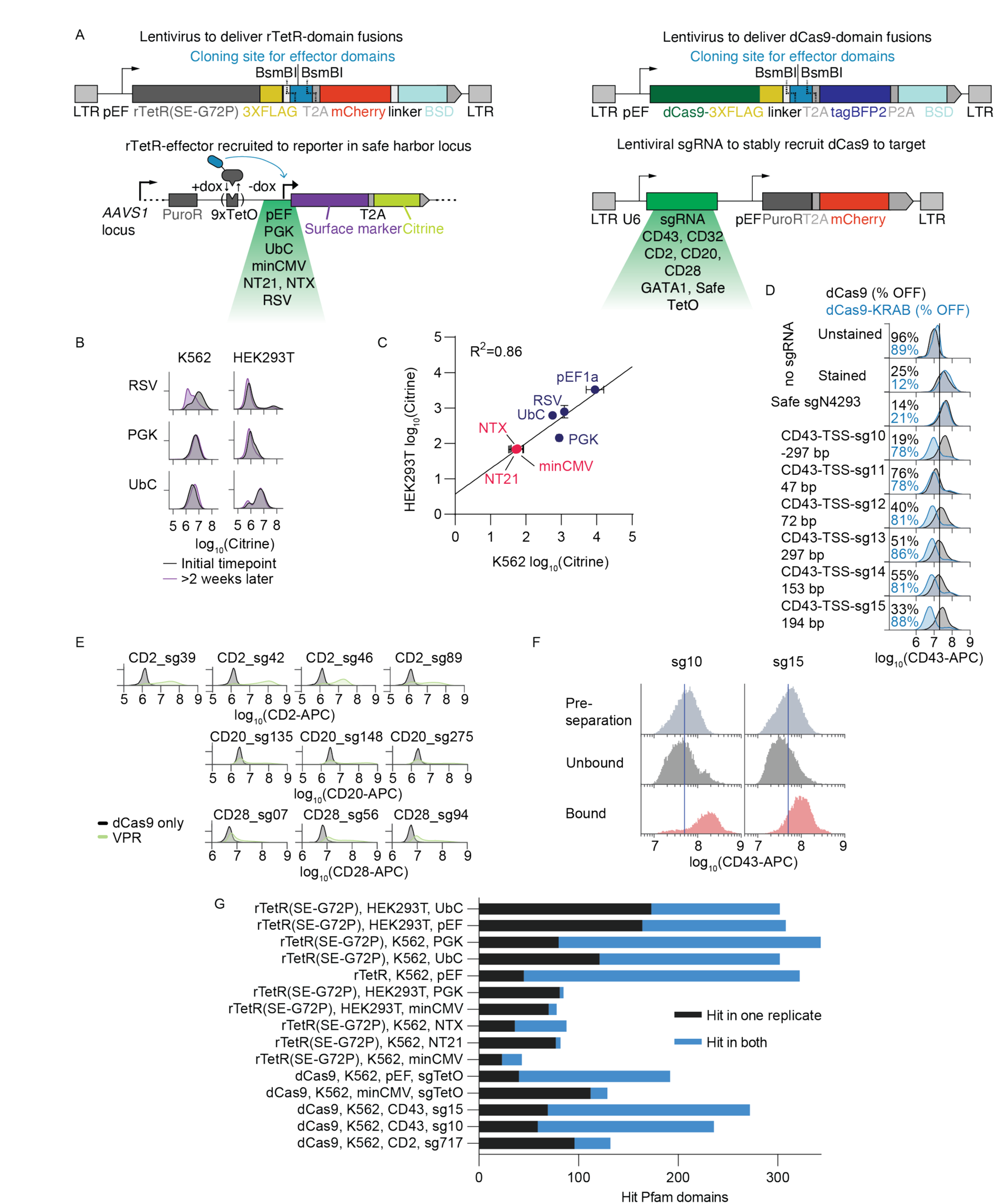
HT-recruit to varied reporters and endogenous gene targets in K562 and HEK293T cells. **A**. Schematics of recruitment constructs with rTetR (left) or dCas9 (right) as the DBD. These can be used to recruit effectors to either reporter constructs that are integrated into the AAVS1 safe harbor, or to endogenous genes. Safe sgRNAs target the genome at a safe location (Morgens et al., 2016) and the TetO sgRNA targets the synthetic reporter at an overlapping location as rTetR (the TetO motif upstream the promoter). **B**. Observation of background silencing at the RSV, PGK, and UbC promoters in K562 and HEK293T cells. Reporters were stably integrated at the AAVS1 safe harbor locus in both cell types by TALEN-mediated homology directed repair. **C**. Mean fluorescent intensity (MFI) of the Citrine reporter under different promoters: minCMV, NT21, NTX, PGK, UbC, pEF1α, PGK, and RSV. Each dot represents a mean taken from three replicates. Promoters with red text represent reporters that are OFF in both cell types. **D**. Testing CD43 TSS-targeting sgRNAs in K562 cells to identify guides that allow for analysis of KRAB-mediated silencing of the CD43 gene. Silencing was measured by CD43 surface marker immunostaining and flow cytometry 7 days after lentiviral sgRNA infection in stable dCas9-ZNF10 KRAB (blue) or dCas9-only (black) cell lines. Data is gated for sgRNA (mCherry^+^, only in samples with an sgRNA) and dCas9 (BFP^+^) delivery (n=1 infection). **E**. dCas9-activators targeting CD2, CD20, CD28 surface marker genes in K562 cells. sgRNAs were stably installed by lentiviral delivery and puromycin selection. Then 500 ng of dCas9 plasmids were electroporated into 1e6 cells. Two days later, cells were stained for surface CD2 (APC), CD20 (APC), or CD28 (PE) expression and analyzed by flow cytometry after gating for dCas9 (BFP) and the stably expressed sgRNA (GFP) (n=1). **F**. Magnetic separation using anti-CD43 antibody and Protein G Dynabeads performed 9 days after lentiviral delivery of dCas9-Pfam library in K562 cells expressing sgRNAs that target CD43. Separation is measured by flow cytometry with gates for dCas9 (BFP^+^) and sgRNA (mCherry^+^). **G**. Overlap in hit Pfam domains in both biological replicates for HT-recruit screens, defined as elements that had ≥5 sequencing reads in Bound and Unbound and log_2_(OFF:ON) scores 2 standard deviations beyond (i.e. higher for repressors, lower for activators) the median of the poorly-expressed controls.

**Supplementary Figure 2.**
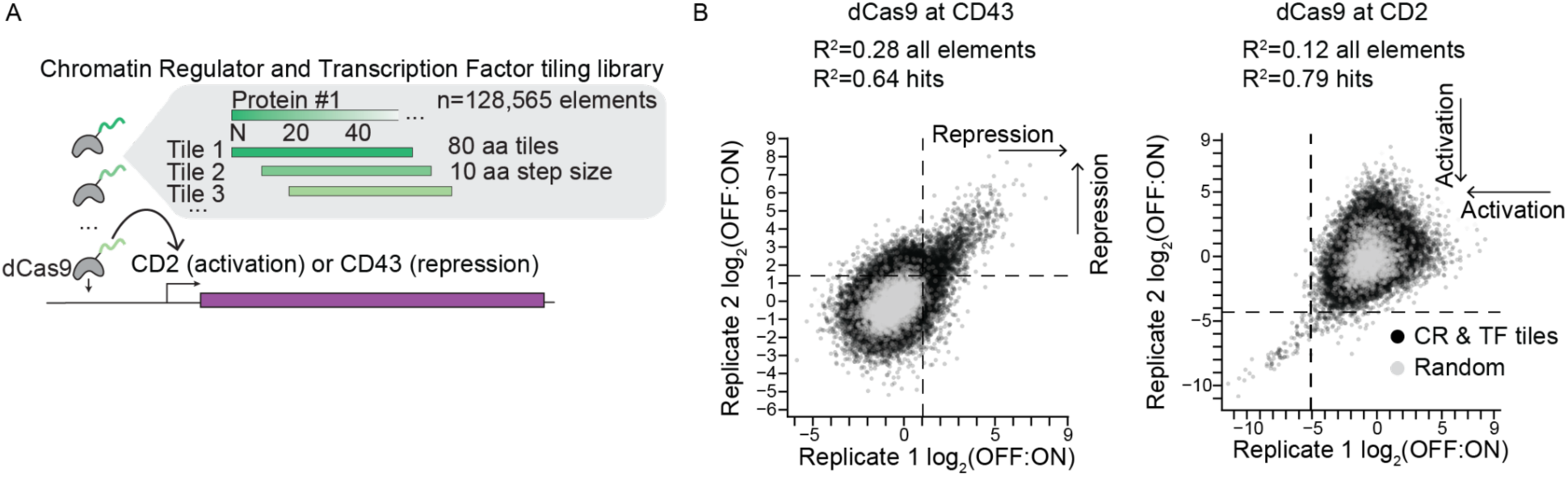
dCas9 fusions to tiles of all human chromatin regulator and transcription factors uncovers unannotated effectors. **A**. Schematic of a library tiling all human chromatin regulator and transcription factor (CR & TF) proteins in 80 amino acid tiles with a 10 amino acid step size (n=128,565 elements) (DelRosso et al., 2023). This library was fused to dCas9 and used to target CD43 with sg15 and CD2 with sg717. **B**. Replicates of CR & TF library fused to dCas9 and recruited to CD43 or CD2 in K562 cells. Hit threshold shown at 2 standard deviations above (for CD43 screen) or below (CD2) the median of the random controls. Elements with >20 sequencing counts in both the Bound (ON) and Unbound (OFF) samples are included. The linear regression goodness of fits (R^2^) are shown for all elements and for the subset that are hits in both replicates.

**Supplementary Figure 3.**
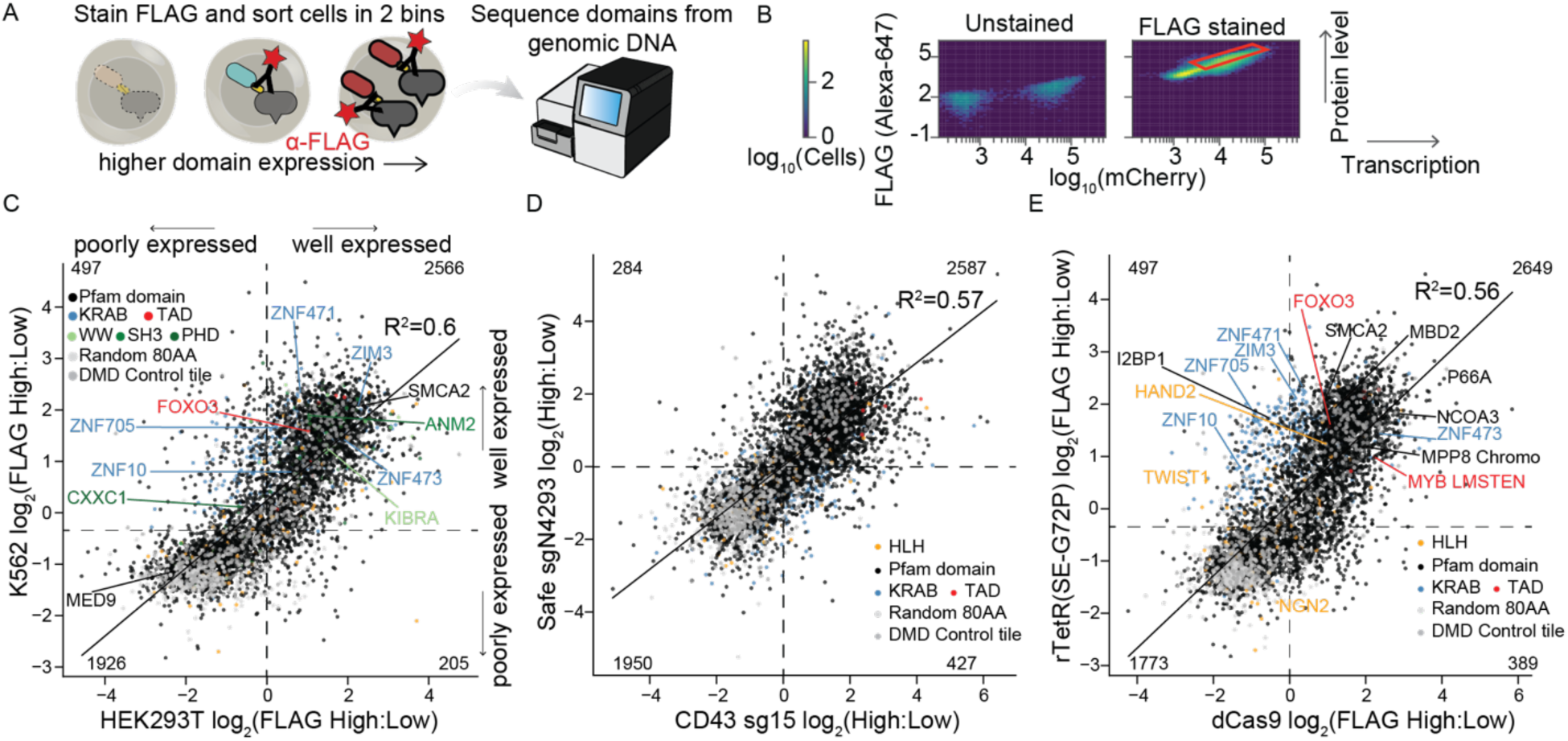
Pooled measurement of domain expression level across cell types and DBDs. **A**. Schematic of pooled approach to measure domain expression. Cells are permeabilized and then stained with a fluorescent anti-FLAG antibody. Then, cells are sorted into high and low FLAG level bins using gates that account for transcription variability by using the fluorescent delivery marker (mCherry for rTetR constructs and BFP for dCas9 constructs), which is found on the same transcript after the T2A cleavage signal. Genomic DNA is extracted and then the domains are sequenced in those two cell populations. **B**. Gating strategy for sorting based on FLAG stain intensity in HEK293T cells. Example gate shown in red accounts for variation across cells in transcription level of the rTetR-3XFLAG-effector-T2A-mCherry transcript by using mCherry fluorescence. **C**. Comparison of Pfam domain library expression levels between HEK293T and K562 cells (Spearman ρ=0.84). In both cell types, the DBD was rTetR(SE-G72P) and the cell line was the minCMV reporter (n=2 replicates). Data was filtered for elements with >5 reads in both FLAG High and Low samples. Well-expressed domains were identified based on a hit threshold (dashed lines) set 1 S.D. above the median of the random controls. The number of library elements in each quadrant is labeled in the corners. **D**. Comparison of Pfam domain library expression levels when fused to dCas9 and delivered to cell lines expressing either a Safe control sgRNA or an sgRNA targeting CD43 (n=1 replicate). **E**. Comparison of Pfam domain library expression levels when fused to dCas9 or rTetR(SE-G72P) and delivered to K562 cells (n=2 replicates). For dCas9, one replicate each from the Safe sgN4293 and CD43 sg15 cells are averaged.

**Supplementary Figure 4.**
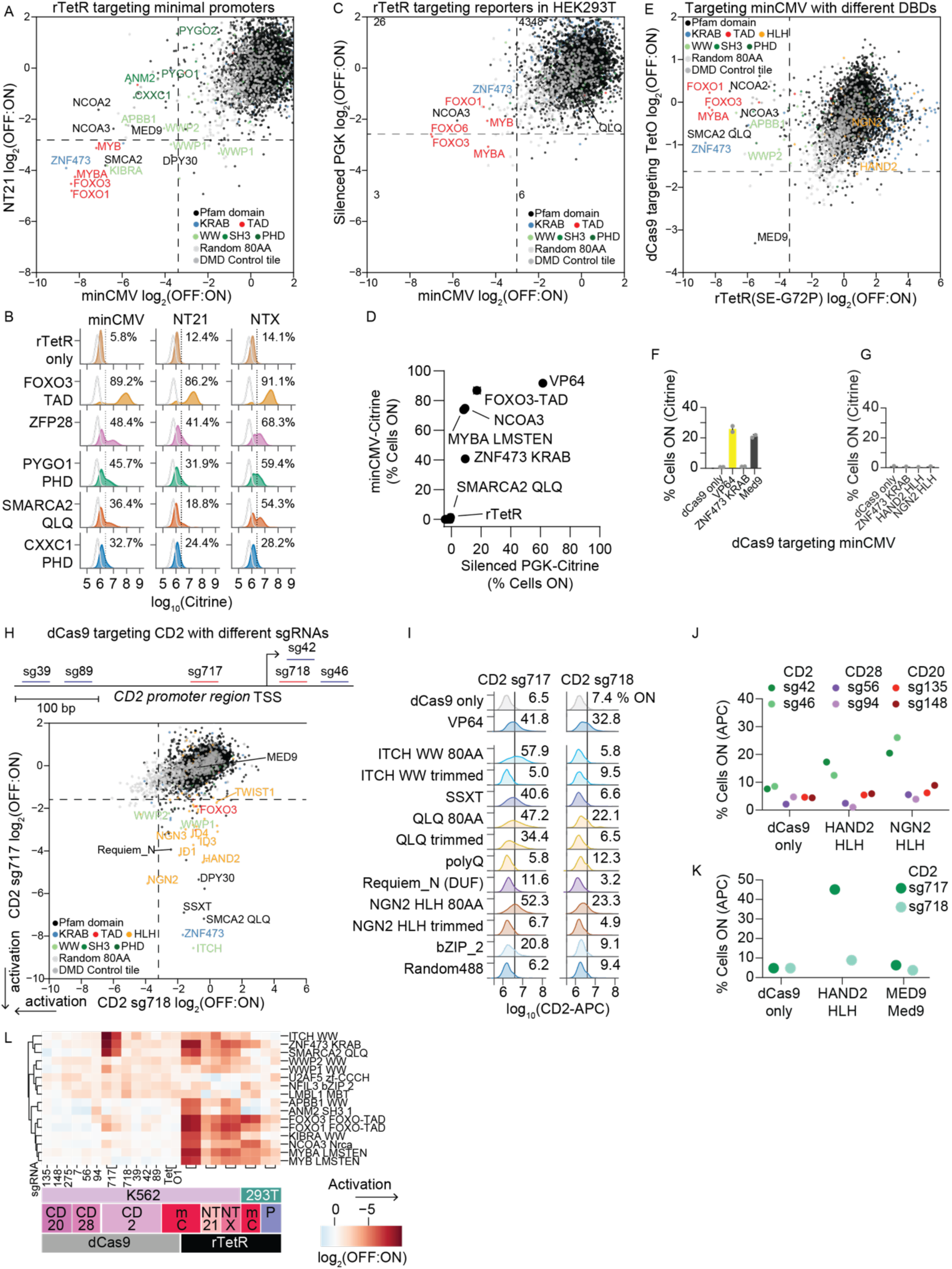
Activators function more similarly at minimal promoters than with a different DBD or at a silenced promoter. **A**. HT-recruit with rTetR targeting two minimal promoters, minCMV and NT21 in K562 cells (n=2 biological replicates). Dashed lines show hit thresholds at 2 standard deviations above the median of the poorly-expressed domains. **B**. Validation of SMARCA2 QLQ and CXXC1 and PYGO1 PHD activator domains across core promoter reporters (minCMV, NT21, NTX) in K562 cells. rTetR-activator fusions or the rTetR-only negative control were delivered by lentivirus to reporter cells. After selection, cells were treated with 1000 ng/ml doxycycline for 2 days to induce reporter activation. The percent of cells activated was measured by flow cytometry for the Citrine reporter, after gating for delivery with mCherry (n=3 replicates). **C**. HT-recruit with rTetR targeting the minCMV and background-silenced PGK reporters in HEK293T cells (n=2 biological replicates per promoter). rTetR-domain fusions were recruited to the reporter with 1000 ng/ml doxycycline for 2 days. The number of Pfam domains in each quadrant is labeled. **D**. Individual validations of activators across PGK and minCMV promoter types in HEK293T as measured by average percentage of cells ON normalized to no-doxycycline control. Cells were treated with 1000 ng/ml doxycycline for 2 days to induce reporter activation (n=2 independently transduced replicates for each promoter type). **E**. HT-recruit with dCas9 recruitment of activators with an sgRNA that binds the TetO site upstream the minCMV reporter in K562 cells (n=2 biological replicates). **F**. dCas9-activators recruited with an sgRNA that binds the TetO site upstream the minCMV reporter in K562 cells (n=2 infection replicates shown as dots). dCas9 fusions were delivered by lentivirus, selected with blasticidin starting on day 5, and cells were analyzed on day 9. Flow cytometry measurements were gated for dCas9 and TetO_sg1 using BFP and mCherry, respectively. Bars show mean and error bars show standard deviation. **G**. Other dCas9-activators recruited in K562 cells with sgRNA that binds the TetO site upstream the minCMV reporter (n=1 replicate). dCas9 fusions were delivered by lentivirus, selected with blasticidin starting on day 3, and cells were analyzed on day 9. Flow cytometry measurements were gated for dCas9 and TetO_sg1 using BFP and mCherry, respectively. **H**. Comparison of HT-recruit with the dCas9-Pfam domain library targeted to the CD2 gene TSS using two different guides (n=2 replicates for sg717 and n=1 for sg718) in K562 cells. Screen measurement was taken 10 days after library delivery. Schematic shows locations of the CD2-targeting guides (sg39 sg89, sg717, sg718, sg42) at the CD2 promoter region. **I**. Recruitment of dCas9-activator hits at the CD2 gene using two different guides (sg717 and sg718) in K562 cells. sgRNA were stably delivered by lentivirus and selected for with puromycin, then dCas9 fusion plasmids were delivered by electroporation, then cells were analyzed 3 days later by flow cytometry for surface stained CD2 after gating for dCas9 (BFP) and sgRNA (GFP). The percentage of cells ON is shown (n=1 replicate). The 80 AA sequences match the library elements while the trimmed sequences match the annotated Pfam domain. The polyQ is a homopolymer of 15 repeated glutamines, which is also found at the C-terminus of the 80 AA QLQ and is not present in the trimmed QLQ. bZIP_2 domain from CEBPE was filtered due to low counts in the screen. **J**. dCas9-HLHs were delivered to K562 cells by lentivirus and selected for with blasticidin, then sgRNAs were delivered by lentivirus and selected with puromycin, then 8 days after sgRNA delivery the cells were stained for the targeted surface markers and measured by flow cytometry. Data was gated for dCas9 with BFP and sgRNA with GFP. **K**. 9 days after lentiviral delivery of dCas9-fusions, K562 cells were immunostained with CD2 antibody to measure gene activation by flow cytometry (n=1). **L**. HT-recruit scores for activators that were hits in ≥5 samples across target, cell-type, and DBD contexts (n=2 replicates per rTetR screen and n=1-2 replicates per sgRNA for dCas9 screens shown as columns). The rows are clustered in an unbiased manner. mC=minCMV and P=PGK.

**Supplementary Figure 5.**
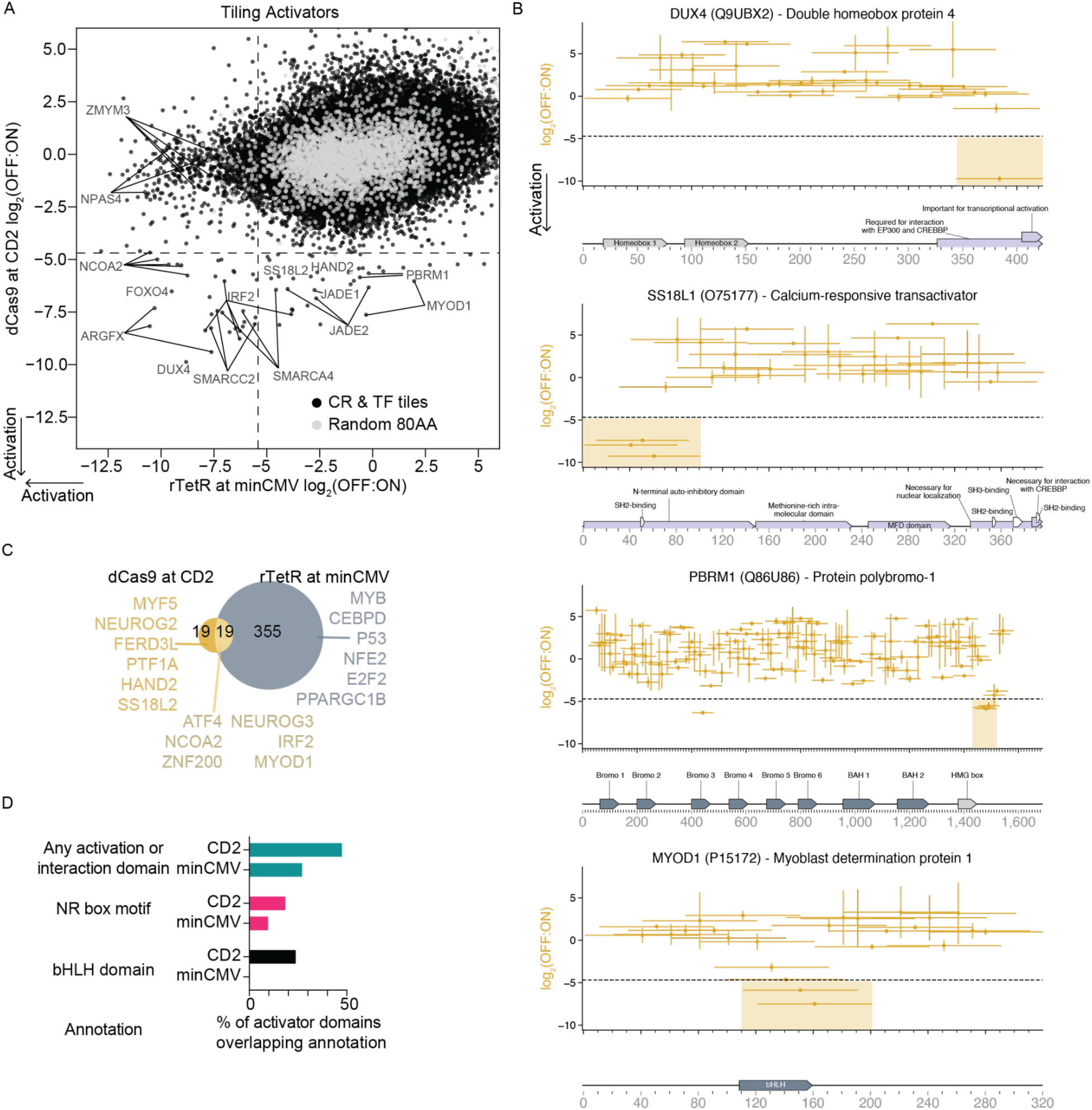
dCas9 fusions to tiles of all human chromatin regulator and transcription factors uncovers unannotated and HLH activators. **A**. dCas9 recruitment of CR & TF tiles to CD2 compared with rTetR recruitment to minCMV. Dashed lines show average of hit thresholds (n=2 replicates per screen). **B**. Proteins with activator hit tiles. Each horizontal line is a tile, and vertical bars show the range (n=2 screen replicates). Dashed horizontal line is the hit calling threshold based on random controls. Hit domains, defined as the sequence from the start of the first hit tile to the end of the last hit tile for a stretch of 1 or more consecutive hit tiles (that are below the hit threshold in both replicates), are shaded. UniProt annotations and Pfam domains are shown below. **C**. Overlap of hit activator domains from different contexts. Shared hits are defined as hit domains with any overlapping sequence. Proteins containing the top 6 strongest hit domains are listed, and for the shared hit category the proteins with the top 6 strongest CD2 activators are listed. **D**. Percentage of hit domains overlapping annotations. NR box motif is associated with recruitment of nuclear receptor coactivators (Leers et al., 1998). Some domains are hits in both contexts or overlap multiple annotations.

**Supplementary Figure 6.**
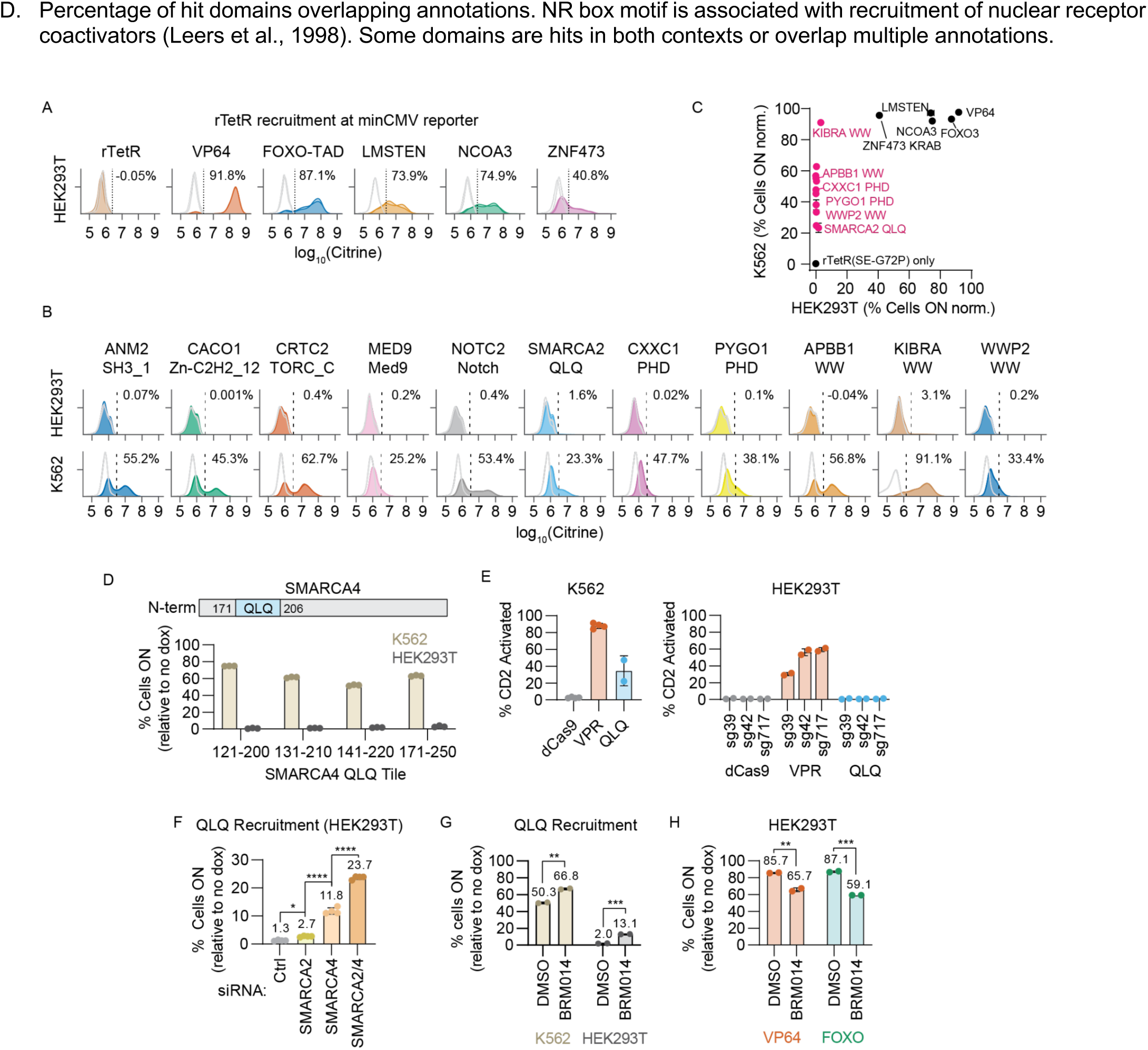
Activators are often stronger in K562 than HEK293T cells. **A**. Individual validations of strong activator domains (FOXO3-TAD, LMSTEN, NCOA3, and ZNF473 KRAB which were previously validated in K562 cells (Tycko et al., 2020)) at the minCMV reporter in HEK293T cells. rTetR(SE-G72P)-domain fusions were delivered to minCMV reporter cells by lentivirus and selected with blasticidin. Cells were treated with 1,000 ng/mL doxycycline for 2 days. Untreated cell distributions are shown in light gray and doxycycline-treated cells are shown in colors, with 2 biological replicates in each condition. The dotted vertical line shows the Citrine gate used to determine the fraction of cells ON. The average percentage of cells ON normalized to no doxycycline control is shown. VP64 and rTetR only were included as positive and negative controls, respectively. **B**. Individual validations of cell-type specific activator domains in HEK293T (top) and K562 (bottom) cells as also shown in Figure 2E. rTetR(SE-G72P)-domain fusions were delivered to minCMV reporter cells by lentivirus and selected with blasticidin. Cells were treated with 1,000 ng/mL doxycycline for 2 days, and then citrine reporter levels were measured by flow cytometry. Untreated cell distributions are shown in light gray and doxycycline-treated cells are shown in colors, with 3 replicates in each condition. The dotted vertical line shows the citrine gate used to determine the fraction of cells ON, and the average fraction ON for the doxycycline-treated cells is shown. **C**. Full set of individual validations of activator hits in K562 and HEK293T cells as measured by average percentage of cells ON normalized to no doxycycline control. Cells were treated with 1000 ng/ml doxycycline for 2 days to induce reporter activation (n=2-3 independently transduced replicates). K562-specific activators are shown as pink dots based on a 2σ hit threshold from the minCMV activator screen. **D**. minCMV reporter activation in K562 and HEK293T cells after 2 days of recruitment of SMARCA4 tiles overlapping the QLQ domain. The QLQ domain from SMARCA4 is not included in our Pfam domain library and shares ∼81% homology with SMARCA2 QLQ. Bars show mean and dots show 3 replicates. **E**. Left: Percentage of CD2 endogenous gene activated after transient expression and targeting of dCas9-effector fusions to the CD2 endogenous gene for 3 days in K562 cells using sg717 and sg718 guides. Cells were immunostained for CD2 (APC) expression and analyzed by flow cytometry after gating for dCas9 (TagBFP) and sgRNA (GFP). Each dot represents an independently transfected sgRNA (n=2-4). Right: Percentage of CD2 endogenous gene activated after transient expression and targeting of dCas9-domain fusions to the gene using multiple sgRNAs (sg39, sg42, sg717) for 3 days in HEK293Ts. Cells were immunostained for CD2 (APC) expression and analyzed by flow cytometry after gating for dCas9 (TagBFP) and sgRNA (GFP). Each dot represents an independently transfected biological replicate (n=2). dCas9 only and VPR are negative and positive controls, respectively. **F**. Activation of HEK293T minCMV reporter cells after 2 days of QLQ recruitment with doxycycline and SMARCA2/4 knockdown with co-transfected siRNA. Bars show mean and dots show 4 transfection replicates; asterisks show result of two-tailed Tukey’s test (Ctrl vs. SMARCA2: \**p*=0.040; SMARCA2 vs. SMARCA4: \*\*\*\**p*= 1.3e-9; SMARCA4 vs. SMARCA2/4: \*\*\*\**p*=3.2e-11). **G**. Activation of K562 and HEK293T minCMV reporter cells after 2 days of treatment with both doxycycline (to recruit SMARCA2 QLQ) and BRM014, a SMARCA2/4 ATPase inhibitor which reduces the levels of SMARCA2/4 in a chromatin fraction immunoprecipitation (Papillon et al., 2018; Schick et al., 2021). Bars show mean and dots show 2 replicates; asterisks show result of two-tailed unpaired *t*-test (K562: \*\**p*=0.0025; HEK293T: \*\*\**p*=0.0001). **H**. Activation of HEK293T cells after 2 days of treatment with both doxycycline and BRM014 when recruiting VP64 or FOXO3 FOXO-TAD. Bars show mean and dots show 2 replicates; asterisks show result of two-tailed unpaired *t*-test (VP64: \*\**p*=0.0069; FOXO: \*\*\**p*=0.0004).

**Supplementary Figure 7.**
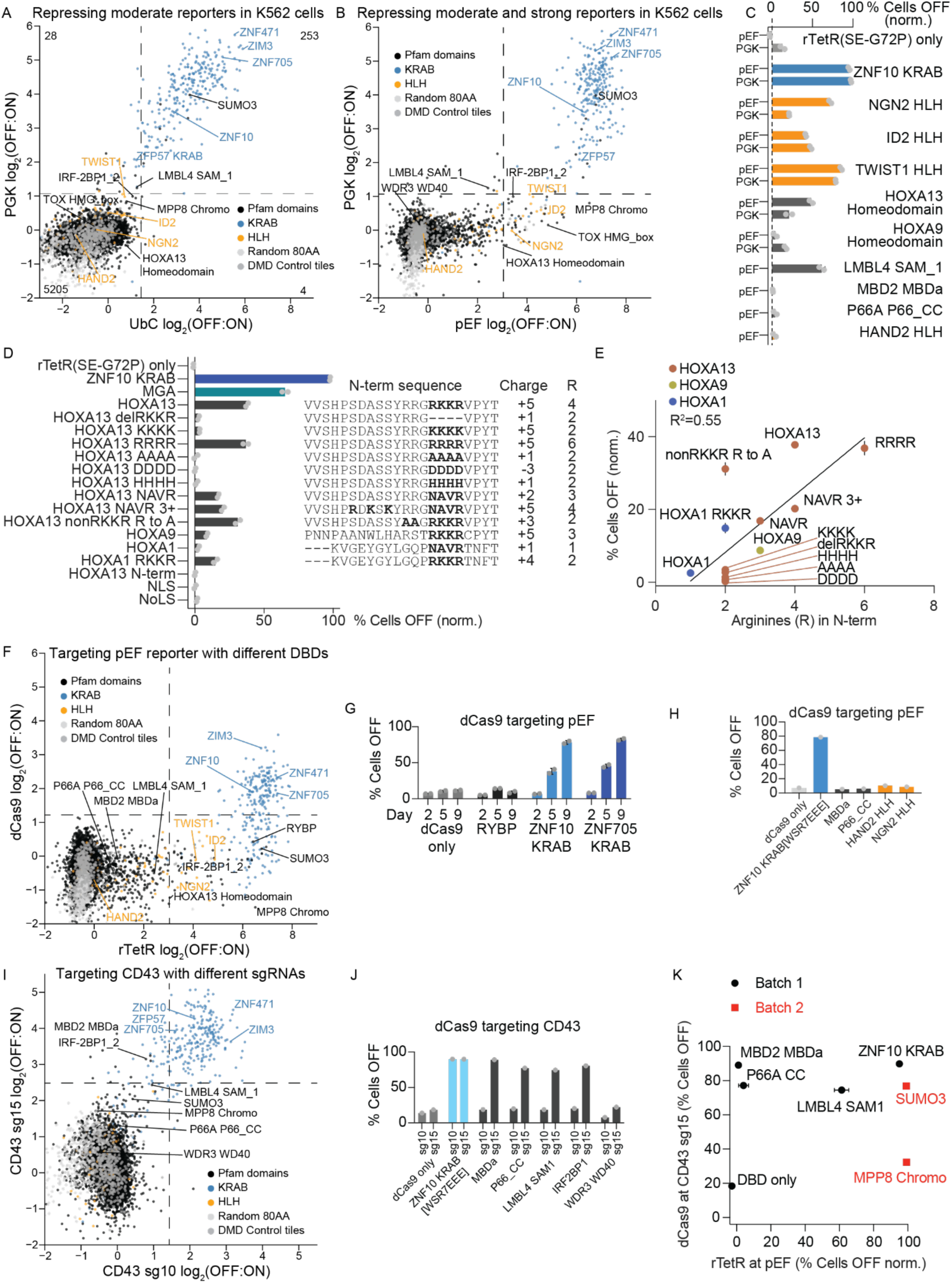
Repressors vary across targets and DBDs. **A**. HT-recruit with rTetR-mediated recruitment at the PGK and UbC reporters in K562 cells (n=2 biological replicates per reporter). **B**. HT-recruit with rTetR-mediated recruitment at the PGK and pEF1α reporters in K562 cells (n=2 biological replicates per reporter). **C**. Individual rTetR-mediated recruitment at the PGK and pEF1α reporters in K562 cells. rTetR fusions were delivered by lentivirus, selected with blasticidin, and then cells were treated with doxycycline for 6 days. The percentage of cells silenced was measured by flow cytometry for the citrine reporter after gating for delivery with mCherry and normalizing to the matched no-doxycycline control (n=2 infection replicates shown as dots). Domains at bottom were only tested in pEF1α reporter cells. **D**. 80 amino acid sequences encoding HOX homeodomains and mutants that vary the RKKR motif and charge of their N-terminal region were fused to rTetR and recruited to the pEF1α reporter in K562 cells. The percentage of cells silenced after 5 days of doxycycline treatment was measured by flow cytometry for the citrine reporter after gating for delivery with mCherry and normalizing to the matched no-doxycycline control (n=2 infection replicates shown as dots). KRAB and MGA are positive control repressors. The HOXA13 N-terminal region alone and a nuclear (NLS) or nucleolar (NoLS) localization signal were also fused to rTetR as additional controls. The RKKR motif and/or the mutated residues are in bold. The net charge and number of arginines in the homeodomain N-terminal regions are shown on the right; these characteristics correlate with repression strength of the wild-type HOX homeodomains (Tycko et al., 2020). **E**. Number of arginines in the N-terminal region of the HOX homeodomains or mutants compared with normalized repression. Dots colored by homeodomain protein. **F**. HT-recruit with dCas9 targeting the pEF1α reporter with an sgRNA targeting the TetO motif compared with rTetR targeting the pEF1α reporter in K562 (n=2 biological replicates per DBD). **G**. dCas9 recruitment of repressors with an sgRNA that binds the TetO site upstream the pEF1α reporter in K562 cells (n=2 infection replicates shown as dots). dCas9 fusions were delivered by lentivirus and cells were analyzed 2-9 days later with blasticidin selection starting at day 5. Flow cytometry measurements were gated for dCas9 and TetO_sg1 using BFP and mCherry, respectively. Bars show mean and error bars show standard deviation. **H**. dCas9 recruitment of Pfam domains with an sgRNA that binds the TetO site upstream the pEF1α reporter in K562 cells (n=1 replicate). dCas9 fusions were delivered by lentivirus, selected with blasticidin starting on day 3, and cells were analyzed on day 9. Flow cytometry measurements were gated for dCas9 and TetO_sg1 using BFP and mCherry, respectively. **I**. HT-recruit with dCas9 targeting CD43 with sg10 or sg15 (n=2 biological replicates per sgRNA). **J**. dCas9 recruitment of Pfam domains to the endogenous CD43 gene in K562 cells. Cell lines stably expressing an sgRNA were infected with dCas9 lentiviruses, then fixed and stained for analysis by flow cytometry 10 days after infection. Data is gated for the sgRNA (mCherry) and dCas9 delivery (BFP) (n=1 replicate). **K**. Comparison of individual recruitment of repressors across targets and DBDs in K562 cells. For rTetR Batch 1, the percentage of cells silenced was measured by flow cytometry for the citrine reporter 6 days after doxycycline addition with gating for rTetR delivery with mCherry (n=2 infection replicates, error bar shows standard deviation) and normalized to the no-doxycycline control. For the rTetR measurements on the Batch 2 domains, the experiment was previously performed similarly (Tycko et al., 2020) and cells were analyzed 5 days after doxycycline (n=2). For dCas9 in both batches, silencing was measured by flow cytometry for CD43 staining after gating for dCas9 and sgRNA delivery with BFP and mCherry (n=1).

**Supplementary Figure 8.**
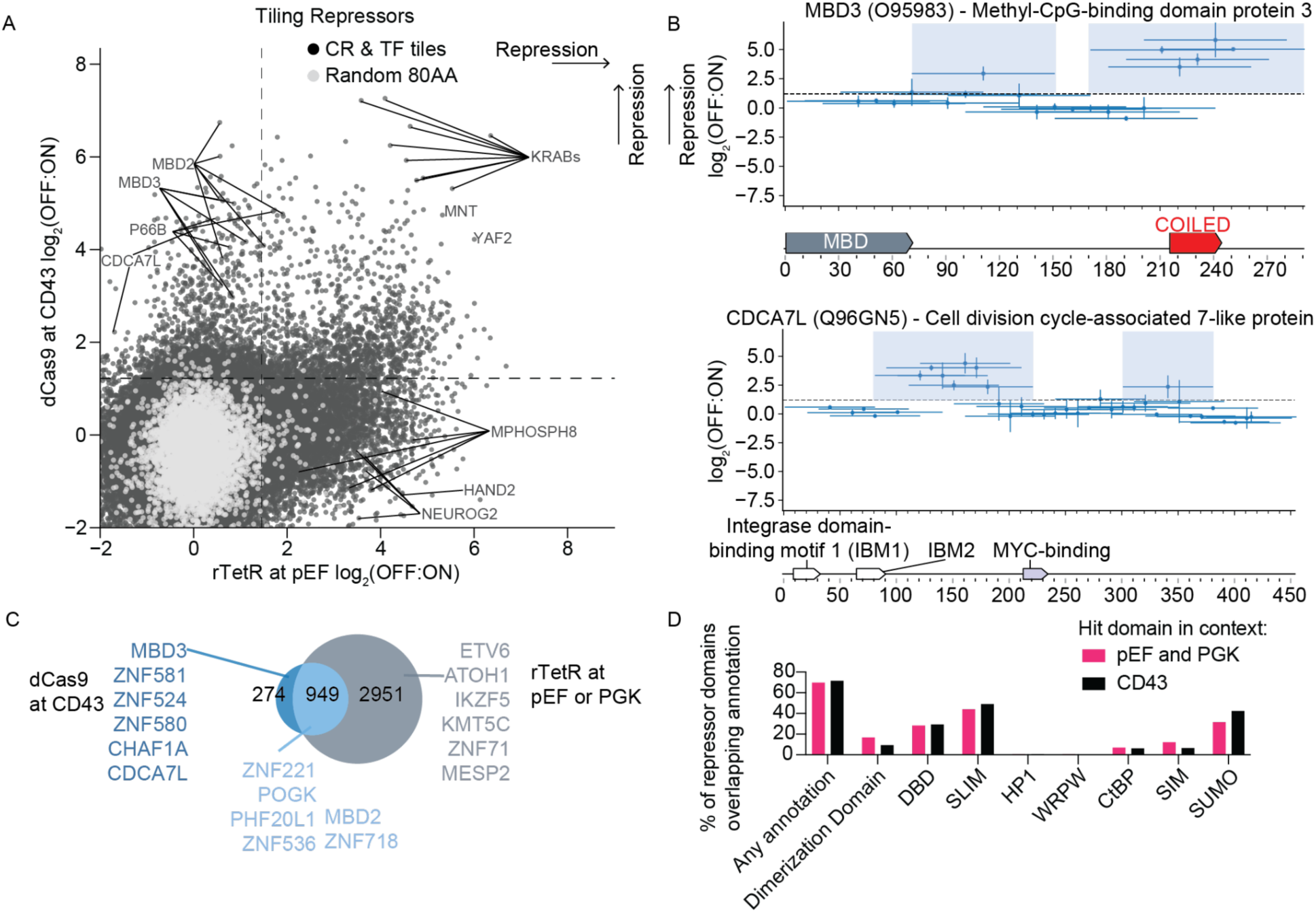
dCas9 recruitment of tiling library to CD43 uncovers unannotated repressors. **A**. Comparison of dCas9 recruitment of the CRTF tiling ilbrary to CD43 with rTetR recruitment to pEF1α. Dashed lines show hit threshold (n=2 replicates per screen) based on random 80AA controls. **B**. Tiling Methyl-binding domain protein MBD3 and CDCA7L. Each horizontal line is a tile, and vertical bars show the range (n=2 screen replicates). Dashed horizontal line is the hit calling threshold based on random controls. **C**. Overlap of hit repressor domains from different contexts. Proteins containing the top 6 strongest hit domains are listed, and for the shared hit category the proteins with the top 6 strongest CD43 repressors are listed. **D**. Percentage of hit domains overlapping a curated set of annotations of interest. SLIM are short linear interaction motifs. HP1, WRPW, and CtBP motifs are associated with co-repressor recruitment. SIM are SUMO-interacting motifs, and SUMO are SUMOylation sites. ‘Any annotation’ refers to any of these. To conservatively select hit domains from the rTetR screens, here we used domains that are hits with both pEF and PGK. Some domains are hits in both rTetR and dCas9 contexts or overlap multiple annotations.

**Supplementary Figure 9.**
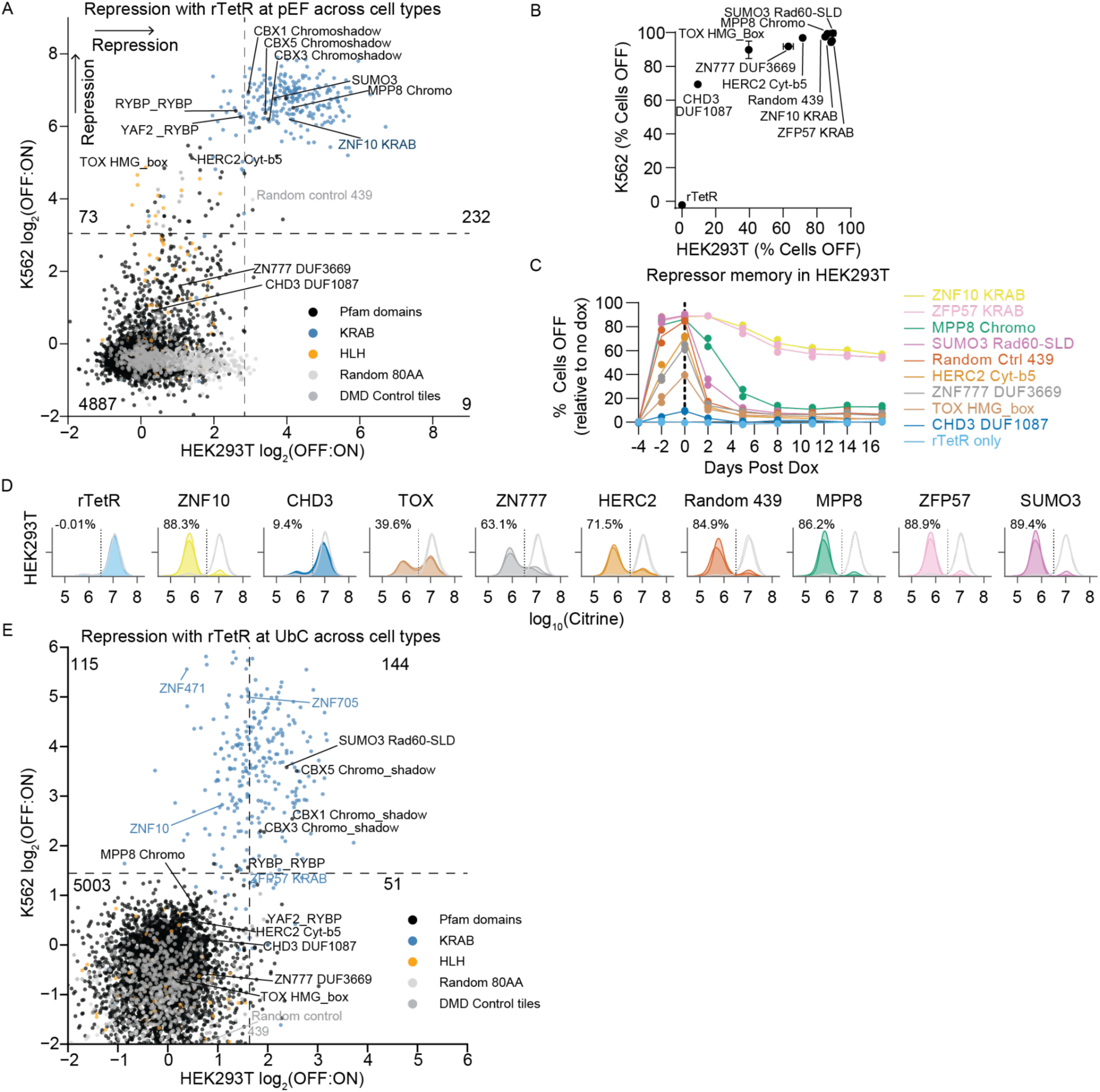
Repressors compared in K562 and HEK293T cells. **A**. HT-recruit with rTetR targeting the strong pEF1α promoter in K562 and HEK293T cells. Repression strength is measured by log_2_(OFF:ON) such that domains with high log_2_(OFF:ON) values are strong pEF1α repressors (n=2 replicates per cell type). rTetR-domain fusions were recruited to the pEF1α reporter with 1000 ng/ml doxycycline for 5 days in K562 and for 4 days in HEK293T cells. Dashed lines show hit thresholds at 2 standard deviations above the median of the poorly-expressed domains. The number of library elements in each quadrant is labeled. **B**. rTetR fusions were delivered via lentivirus to the pEF1α reporter K562 or HEK293T cells. After selection, cells were treated with 1000 ng/ml doxycycline for 5 days in K562 and 4 days in HEK293T cells. The percent of cells silenced was measured by flow cytometry for the citrine reporter after gating for delivery with mCherry. Dot shows mean, error bars are standard deviation (n=2 replicates). K562 data was previously published in (Tycko et al., 2020) and is re-plotted here for comparison purposes. **C**. Silencing and memory dynamics of repressors at the pEF1α reporter in HEK293T cells. Cells were treated with 1000 ng/ml doxycycline for 4 days, then doxycycline was removed to release recruitment and measure epigenetic memory. Silencing was measured by flow cytometry for the Citrine reporter after gating for rTetR(SE-G72P)-domain delivery with mCherry. Mean percentage of cells silent (line) were calculated from 2 biological replicates (dots). **D**. Day 4 of recruitment of repressor domains in HEK293T cells. The 80 amino acid sequences from the Pfam library were used. Untreated cell distributions are shown in light gray and doxycycline-treated cells are shown in colors (n=2 biological replicates shown as lines). The dotted vertical line shows the citrine gate used to determine the fraction of cells OFF. The average percentage of cells OFF normalized to no doxycycline control is shown. ZNF10 KRAB and rTetR only were included as positive and negative controls, respectively. **E**. HT-recruit with rTetR targeting the moderately strong UbC promoter in K562 and HEK293T cells (n=2 replicates per cell type). rTetR-domain fusions were recruited with 1000 ng/ml doxycycline for 5 days in K562 and for 4 days in HEK293T cells.

**Supplementary Figure 10.**
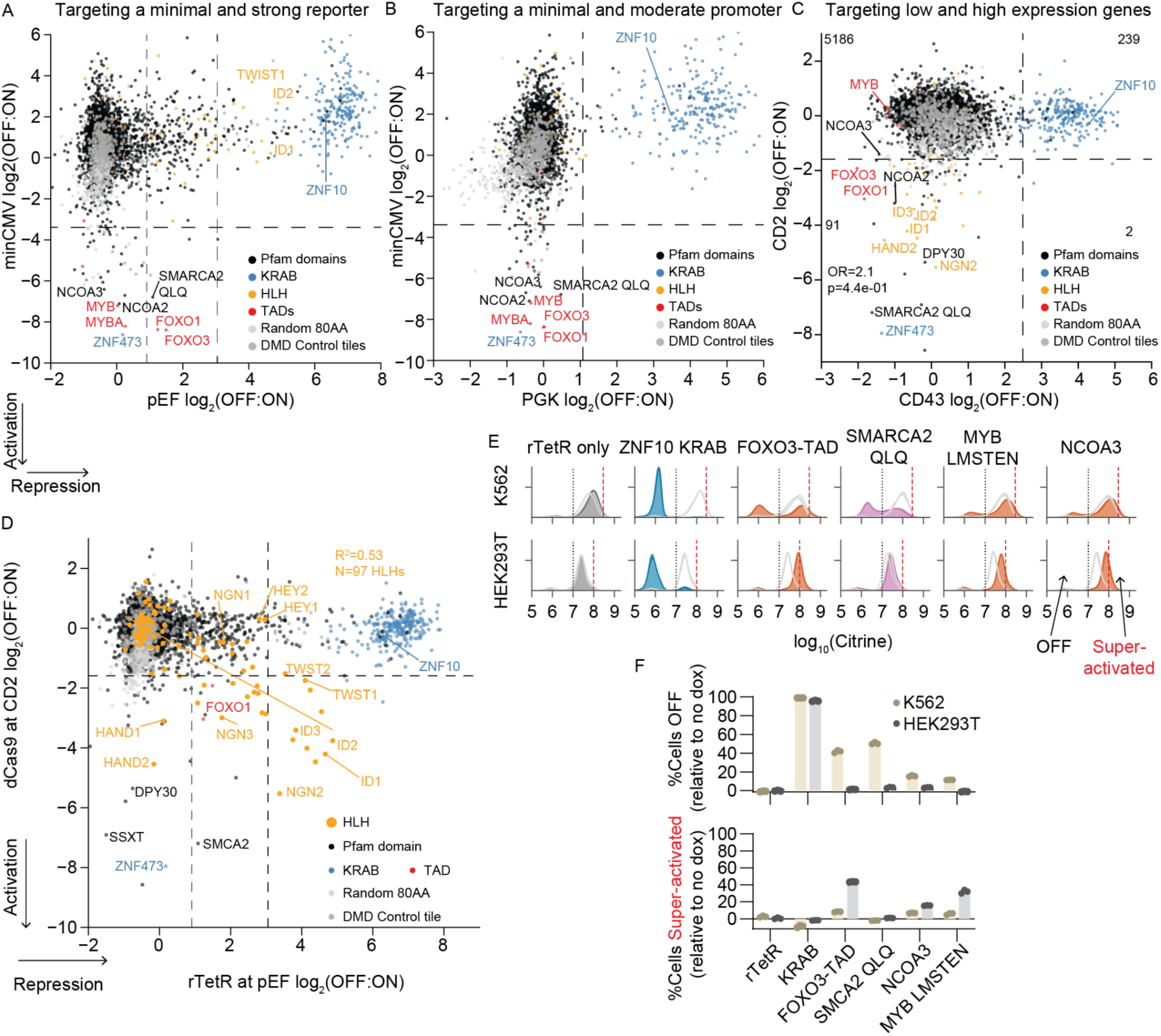
Some activators can super-activate or bifurcate expression of a strong reporter. **A**. HT-recruit with rTetR targeting the pEF1α promoter for 5 days and to the minCMV promoter for 2 days in K562 cells (n=2 biological replicates per reporter). Note the conservative hit threshold (black dashed line, 2 SD beyond median of poorly expressed domains), chosen to identify robust effectors, calls the strongest effects as hits; some sub-threshold domains might have weaker activity. The gray dashed vertical line shows a lower threshold, equivalent to the strength of the weakest repressor that was individually validated. **B**. HT-recruit with rTetR targeting the PGK promoter for 5 days and to the minCMV promoter for 2 days in K562 cells (n=2 biological replicates per reporter). **C**. HT-recruit with dCas9 targeting CD43 with sg15 and CD2 with sg717 in K562 cells (n=2 biological replicates per gene). The number of domains in each quadrant is labeled and the odds ratio (OR) and p-value from Fisher’s exact test is shown. **D**. HT-recruit with dCas9 to activate the endogenous gene CD2 using sg717 compared with repression of pEF1α promoter with rTetR in K562 cells (n=2 replicates per screen). Orange line shows best fit to data from HLH domains (shown in larger orange dots). Black dashed line shows conservative hit threshold; gray dashed vertical line shows lower threshold as in (**A**). **E**. Activating effector domains (FOXO3-TAD, SMARCA2 QLQ, MYB LMSTEN, NCOA3) fused to rTetR were stably integrated into K562 and HEK293T cells with lentivirus. Domains were recruited to the constitutive pEF1α reporter in both ce types for 5 days with doxycycline and then analyzed by flow cytometry for repression. Untreated cell distributions are shown in light gray and doxycycline-treated cells are shown in colors, with 3 biological replicates. The two dotted vertical line shows the Citrine gate used to determine the fraction of cells OFF and super-activated. rTetR only and ZNF10 KRAB were included as negative and positive repressor controls, respectively. **F**. Top: Quantification of data in (**D**). The normalized percentage of K562 or HEK293T cells OFF after recruitment at pEF1α with activating effector domains (n=3 biological replicates). Bottom: The normalized percentage of K562 or HEK293T cells super-activated to a higher ON state (n=3 biological replicates).

**Supplementary Figure 11.**
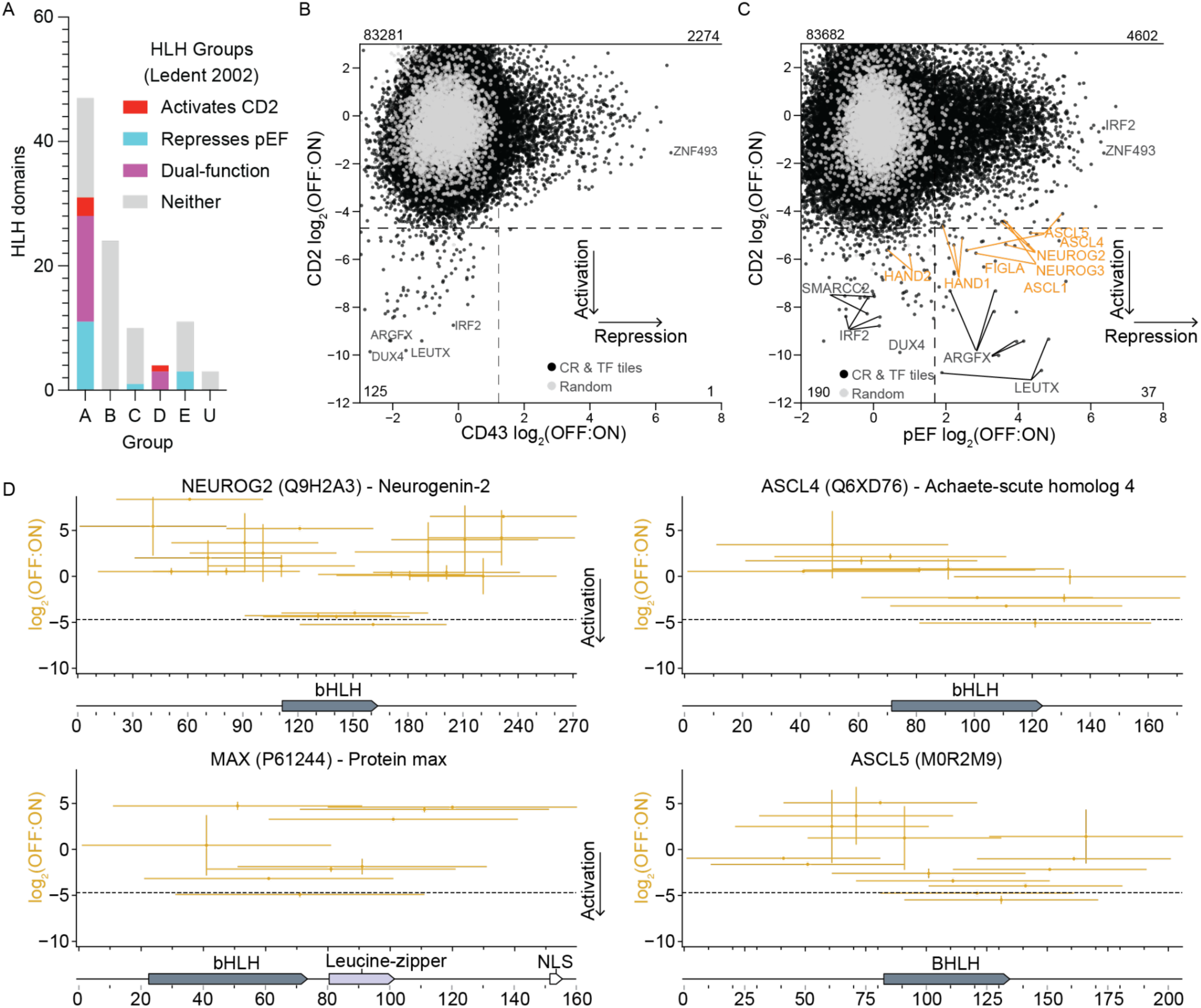
Dual-functioning HLH domains. **A**. Classification of HLH effector hits from the Pfam library screens at pEF with rTetR and CD2 with dCas9 in K562 cells. Hits are shown as colors and compared with phylogenetic grouping defined by (Atchley & Fitch, 1997; Ledent et al., 2002; Torres-Machorro, 2021) and reported as used here in (Torres-Machorro, 2021). For this analysis, an HLH domain is counted as repressing pEF if the average log_2_(OFF:ON) across biological replicates is above the lower threshold (>0.9, equivalent to the weakest individually validated repressor, **Methods**). U=Unclassified. **B**. Comparison of dCas9 recruitment of the CRTF tiling ilbrary to CD2 and CD43. Dashed lines show hit thresholds at 2 standard deviations below or above the median of the random controls (n=2 replicates per screen). Some example hits are labeled with their protein. **C**. Comparison of dCas9 recruitment of the CRTF tiling ilbrary to CD2 with rTetR recruitment to pEF. Dashed lines show hit thresholds at 2 standard deviations below or above the median of the random controls (n=2 replicates per screen). Select hits are labeled with their protein, and the labels are orange for HLH proteins. **D**. Tiling HLH proteins. Each horizontal line is a tile, and vertical bars show the range (n=2 screen replicates). Orange lines show activation of CD2. Dashed horizontal line is the hit calling threshold. UniProt annotations and Pfam domains are shown below. The bHLH domains contain a short N-terminal basic DNA-binding region followed by an HLH heterodimerization region.

**Supplementary Figure 12.**
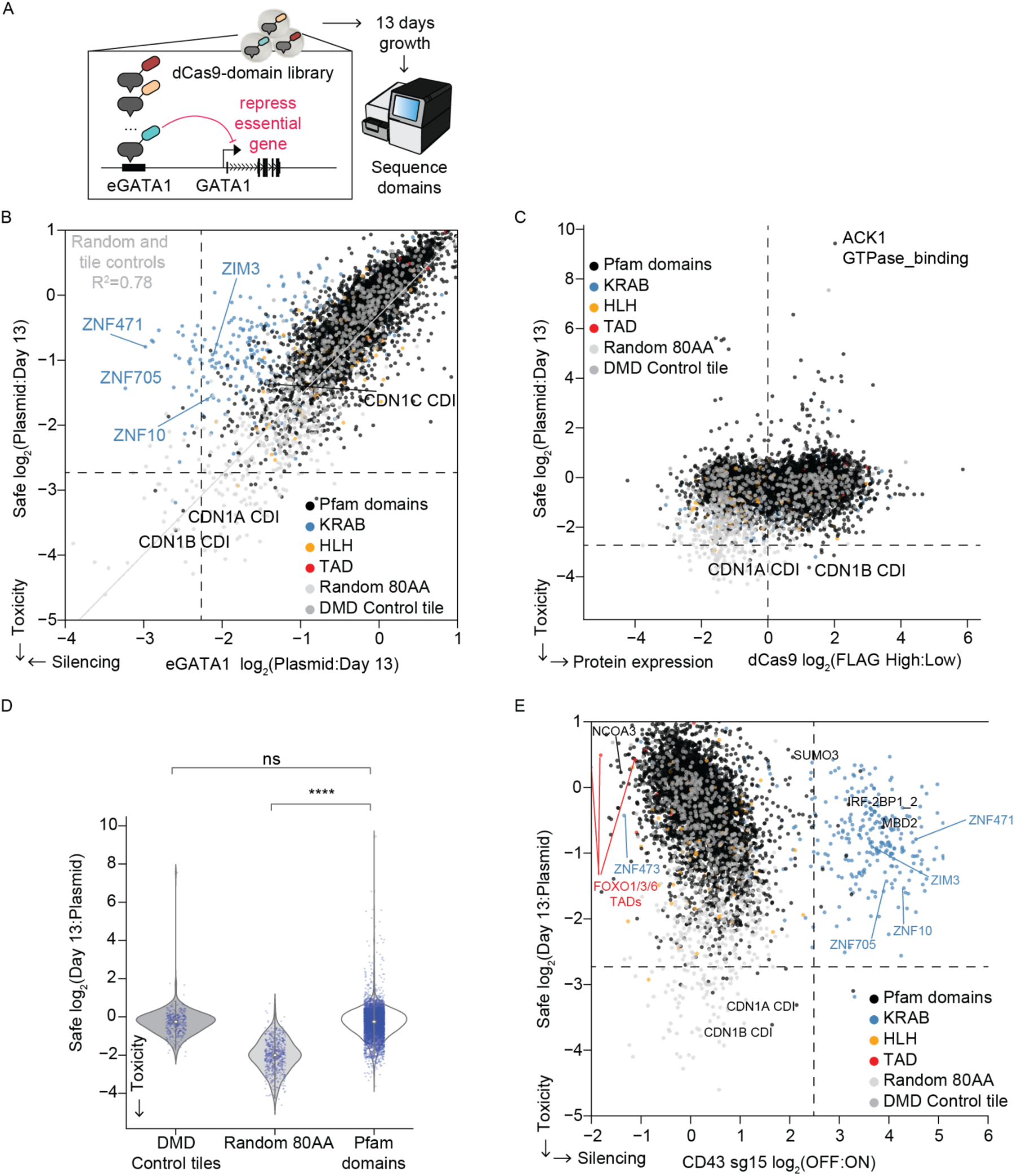
Pooled measurement of growth effects with dCas9-Pfam domain library. **A**. CRISPR HT-recruit to identify repressor domains by targeting the eGATA1 enhancer of the essential gene GATA1 and using growth as a selection strategy. K562 cells stably expressing an sgRNA targeting the enhancer or a safe control were infected with the dCas9-Pfam domain library. After 13 days of growth, genomic DNA was extracted and domains were sequenced. Growth phenotypes were quantified by comparing the domain counts at day 13 relative to the initial plasmid pool. **B**. Comparison of CRISPR HT-recruit growth phenotypes using a safe-targeting sgRNA (N4293) and an eGATA1 enhancer-targeting sgRNA (n=2 replicates per screen). Gray line is fit to random and DMD tiling controls combined. Dashed lines show hit thresholds drawn 2 standard deviations below the median of the poorly expressed controls. **C**. Comparison of growth effects measured with the safe sgRNA and expression levels measured with the FLAG tag (n=2 replicates per measurement). The threshold for expression is 1 standard deviation above the median of the random controls. **D**. Growth effects for sub-libraries, with white dot at median, bars to quartiles and a vertical line to 1.5x the interquartile range. Overlaid dots show individual domains. **** is p<1e-4 for two-sided Mann-Whitney-Wilcoxon test and ns is not significant. **E**. Silencing of CD43, which is not a growth gene, compared with growth effects measured with the safe sgRNA (n=2 replicates per screen).

**Supplementary Figure 13.**
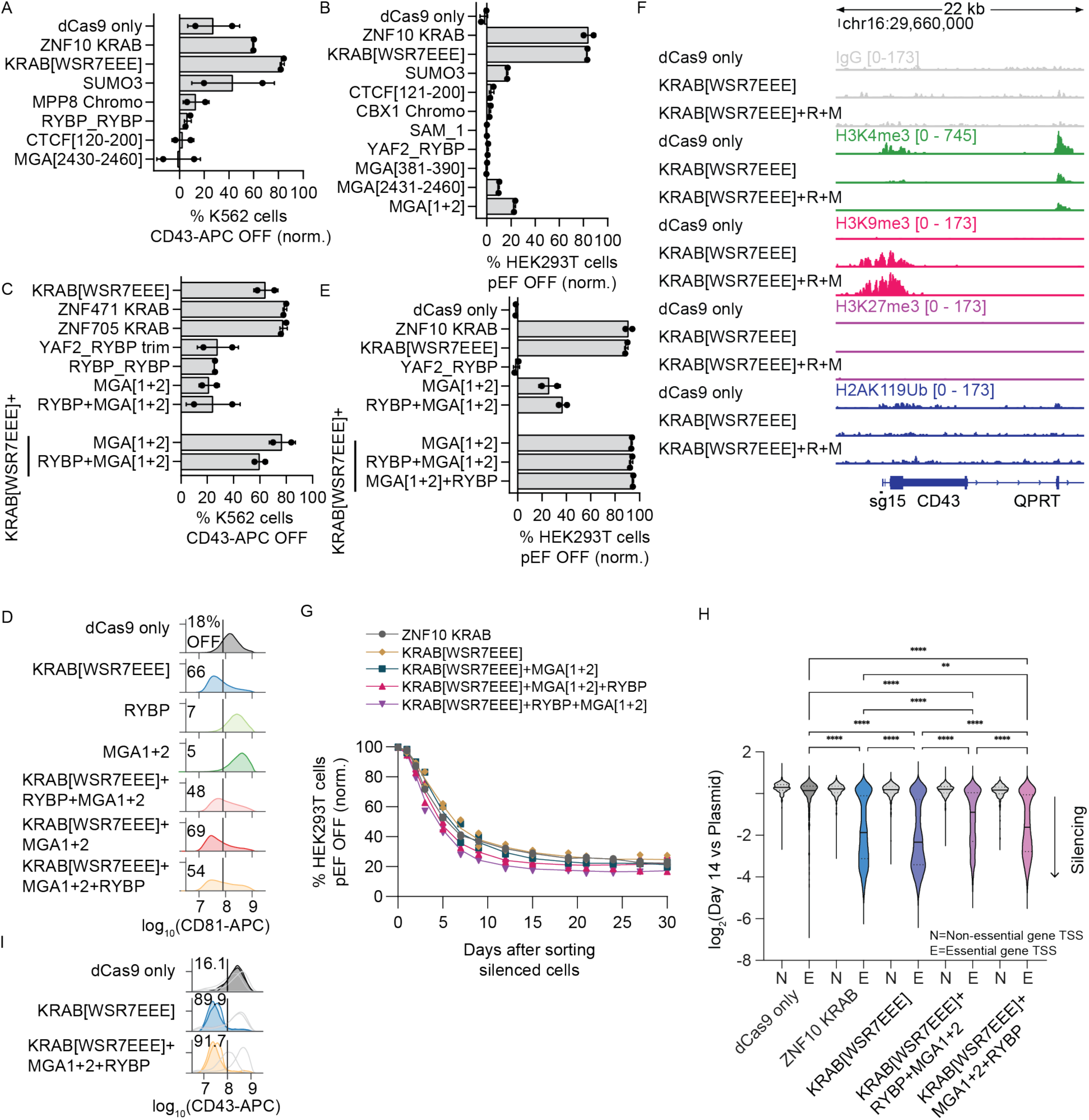
KRAB is most efficient of selected domains and combinations. **A**. Recruitment of dCas9-effectors to the endogenous surface marker gene, CD43, in K562 cells for 7 days. Means and standard deviations are from two independent infections using either CD43 sg10 or sg15, after gating for dCas9 (BFP) and guide (mCherry) expression. Percentage of cells OFF were normalized to safe-targeting sgRNA N4293. For reference, all of these effectors function as repressors when recruited with rTetR to the pEF1α reporter in K562 cells (Tycko et al., 2020). **B**. Transient expression and recruitment of dCas9-Pfam domain fusions to the TetO sites upstream of the reporter gene in HEK293T cells for 5 days. Means and standard deviations are from two biological replicates after gating for dCas9 (BFP) and guide (mIFP) expression. Percentage of cells OFF were normalized to safe-targeting sgRNA. **C**. Recruitment of dCas9-repressors and combinations to the endogenous CD43 gene in K562 cells for 9 days. Means and standard deviations are from two independent infections using either CD43 sg10 or sg15, after gating for dCas9 (BFP) and guide (mCherry) expression. The RYBP domain from the RYBP protein (RYBP_RYBP) was used in the repressor combinations. **D**. Recruitment of repressors combinations to CD81 in K562 cells using sg3, after gating for dCas9 (BFP) and guide (mCherry) expression. First the dCas9 fusions were stably delivered by lentivirus and selected for with blasticidin, then the sgRNA was delivered by lentivirus and 9 days later the cells were stained and fixed for flow cytometry analysis. The percentage of cells OFF is shown. **E**. Transient expression and recruitment of various dCas9-repressor combo fusions from (A) to the TetO sites upstream of the reporter gene in HEK293T cells for 5 days. Means are from two biological replicates after gating for dCas9 (BFP) and guide (mIFP) expression. Percentage of cells OFF were normalized to safe-targeting sgRNA. **F**. Chromatin modifications mapped by CUT&RUN after dCas9 recruitment of repressors to the CD43 endogenous gene using sg15. Stable lines expressing both the dCas9 fusion and the sgRNA were selected with both antibiotics and FACS before chromatin was analyzed. **G**. After targeting the dCas9-repressor combo fusions for 5 days by transient transfection at the pEF1α-TagRFP reporter in HEK293T cells, silenced cells were sorted, and memory dynamics was measured by flow cytometry throughout 30 days. Each dot is a biological replicate (n=2). The percentage of cells OFF at each day was normalized to safe-targeting sgRNA. **H**. A CRISPRi benchmarking growth screen for comparing repressors was performed with a library of sgRNAs targeting essential genes. The library was delivered by lentivirus with low MOI into K562 cell lines that stably express a dCas9-repressor fusion, cells were passaged for 14 days, then genomic DNA was extracted and the sgRNAs were sequenced. Depletion of an sgRNA over 14 days of growth relative to the original plasmid pool representation is associated with effector-mediated silencing of the essential genes. Violin shows distribution of average sgRNA-level depletion from 2 screen replicates, solid line shows median, dotted lines are quartiles (N=2468 sgRNAs targeting essential genes and 451 targeting non-essential genes, **** denotes P<0.0001 and ** is P=0.0012 by Kruskal-Wallis test). **I**. Individual dCas9-mediated recruitment of repressor combinations at the endogenous CD43 gene in K562 cells. Cell lines stably expressing the sgRNA were infected with dCas9-repressors lentiviruses, then fixed and stained for analysis by flow cytometry 10 days after infection. Shaded histograms show cells gated for the sgRNA (mCherry) and dCas9 delivery (BFP); the darker shade is sg10 and the lighter colored shade is sg15, and the light gray histogram shows cells that express neither and serve as an internal control. Control lines are shared with **Supplementary Figure 7J**, which was performed in parallel. The average percentage of cells OFF is shown (n=2 replicates per sgRNA for the combination and 1 for others).

**Supplementary Figure 14.**
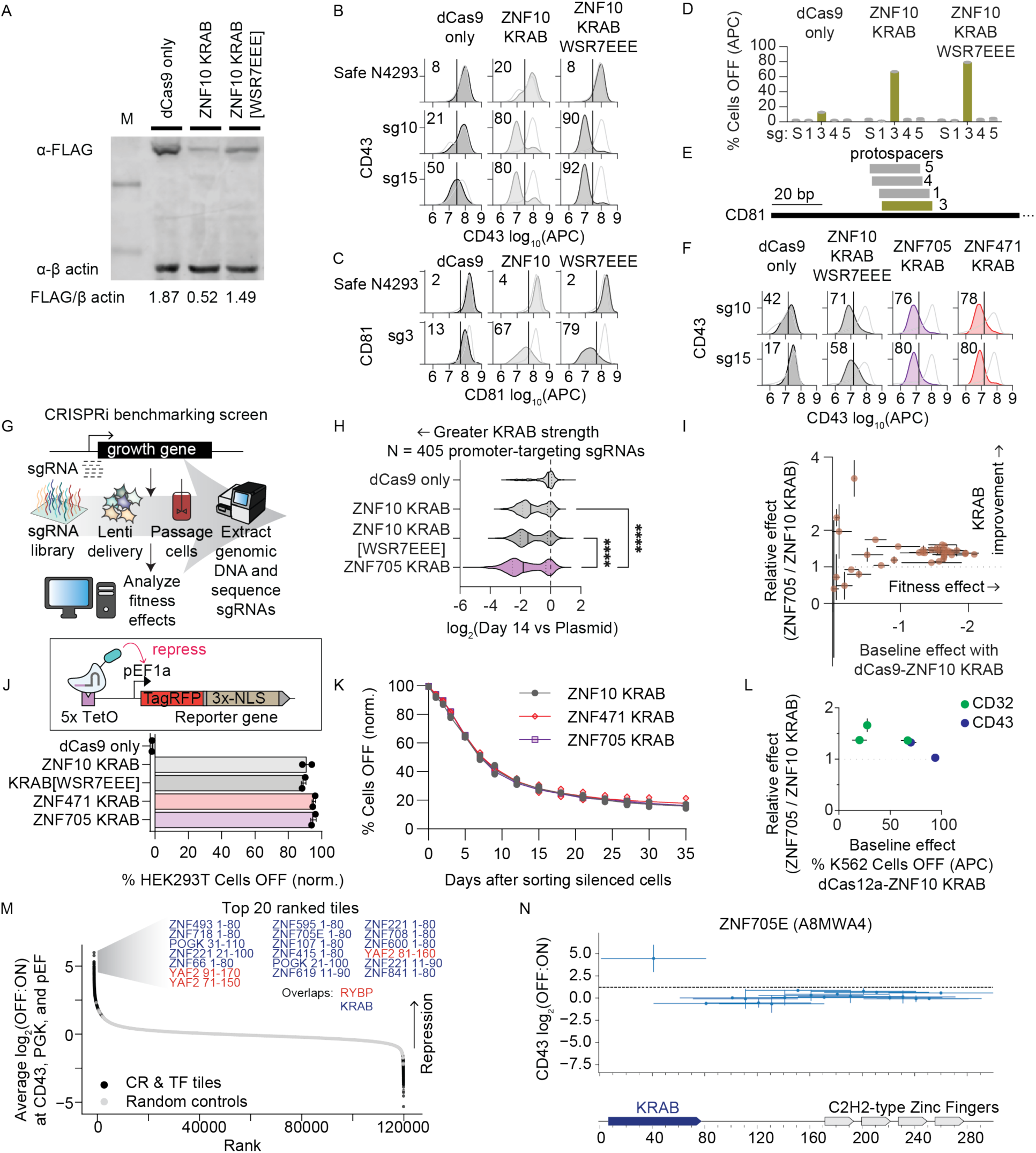
KRAB mutant and paralogs with improved CRISPRi efficiency. **A**. Western blot analysis of dCas9 ZNF10 KRAB and enhanced KRAB[WSR7EEE] mutant fusions. The dCas9-repressors were tagged with 3xFLAG to allow for probing with anti-FLAG antibodies. The band intensity ratio of FLAG to β-actin staining is shown below and used to quantify protein stability. dCas9 only was included as a control. **B**. dCas9 recruitment to the endogenous surface marker gene CD43 in K562 cells. sgRNAs were delivered by lentivirus and selected for with puromycin, then dCas9 constructs were delivered by lentivirus and selected for with blasticidin, and then cells were stained for target gene expression 7 days after dCas9 infection. The shaded histogram shows the cells after gating for both the dCas9 (BFP) and the sgRNA (mCherry) and their percentage of cells OFF is shown. The unshaded histogram shows the cells from the same sample that express neither and serve as an internal control (n=1). Data shared with **Supplementary Figure 13A**. **C**. dCas9 recruitment to the endogenous surface marker gene CD81 in K562 cells. First, dCas9 constructs were delivered by lentivirus, then sgRNAs were delivered by lentivirus, and 3 days later the cells were selected with puromycin and blasticidin. Then, 9 days after sgRNA infection, the cells were stained for CD81 expression, fixed, and analyzed by flow cytometry. The shaded histogram shows the cells expressing both the dCas9 vector (BFP) and the sgRNA (mCherry) their percentage of cells OFF is shown, while the unshaded histogram shows the cells from the same sample that express neither (n=1). **D**. Additional sgRNAs targeting CD81 in K562 cells, after gating for both dCas9 (BFP) and sgRNA (mCherry) (n=1). **E**. Schematic of the CD81 TSS region and the protospacer binding sites for the sgRNAs, colored to match **D**. **F**. dCas9 recruitment of KRAB paralogs to the endogenous gene CD43 in K562 cells. dCas9-repressors were delivered by lentivirus to cells that stably express sg10 or sg15, three days later blasticidin selection was initiated, and then cells were stained for CD43 expression 9 days after dCas9 infection. The shaded histogram shows the cells expressing both the dCas9 vector (BFP) and the sgRNA (mCherry) and their percentage of cells OFF is shown, while the gray unshaded histogram shows the cells from the same sample that express neither and serve as an internal control. Data shared with **Supplementary Figure 13C**. **G**. Schematic of a CRISPRi benchmarking screen for comparing KRAB repressors. K562 cell lines that stably express a dCas9-repressor were infected with a lentiviral library of sgRNAs targeting essential genes with a multiplicity of infection <0.3 such that most cells express one sgRNA. Then cells were passaged for 14 days, genomic DNA was extracted, and the sgRNAs were sequenced to measure depletion associated with fitness effects due to silencing. **H**. Results from CRISPRi benchmarking screen targeting the promoters of essential genes. Greater depletion over 14 days of growth relative to the original plasmid pool representation is associated with stronger effector-mediated silencing of the essential genes. Violin shows distribution of average sgRNA-level depletion from two screen replicates, solid line shows median, dotted lines are quartiles (N=405 sgRNAs, **** denotes P<0.0001 by Kruskal-Wallis test). The dashed line at 0 represents the median of the safe-targeting negative controls. **I**. Comparison of baseline silencing with ZNF10 KRAB and relative improvement with ZNF705 KRAB from CRISPRi benchmarking screen targeting the promoters of 37 essential genes. Effect sizes are the log_2_(fold-change) of sgRNA representation after 14 days of growth relative to the original plasmid pool representation. The median effect across 8-10 sgRNAs per gene was computed and each dot shows its average for two infection replicate screens. Horizontal and vertical error bars show the ranges. Dashed line shows parity between KRAB domains. **J**. Above: Schematic of dCas9-repressor recruitment at the pEF1α-TagRFP-T reporter in HEK293T using TetO targeting sgRNA. Below: Transient transfection of dCas9-KRAB paralogs with an sgRNA targeting the TetO sites upstream of the reporter gene (bar shows mean from n=2 biological replicates shown as dots). Percentage of cells OFF were normalized to safe-targeting sgRNA. **K**. After targeting the dCas9-KRAB paralog fusions for 5 days by transient transfection at the TagRFP reporter in HEK293T cells, silenced cells were sorted, and memory dynamics was measured by flow cytometry throughout 35 days. Each dot is a biological replicate (n = 2). The percentage of cells OFF were normalized to safe-targeting sgRNA. **L**. Comparison of baseline silencing with ZNF10 KRAB and relative improvement with ZNF705 KRAB when dCas12a fusions were recruited to silence CD43 or CD32, with the same method as in Figure 3D. Each dot is colored by the target gene and shows the average for two infection replicates of a guide RNA. Horizontal and vertical error bars show the ranges. Dashed line shows parity between KRAB domains. **M**. Chromatin regulator and transcription factor tiles and random controls are ranked by their mean repression scores from the pEF, PGK, and CD43 screens with the larger library (n=2 replicates per screen). The ZNF705E tile is 99% identical to the ZNF705B/D/F KRAB Pfam domain, which was not itself included in the tiling library because most of the >300 KRAB-containing ZNF proteins were excluded from the library design due to space constraints. The top 20 ranked tiles are listed, with their protein name, tile start, and end position, colored by whether they overlap a KRAB or RYBP domain. **N**. Tiling ZNF705E. Each horizontal line is a tile, and vertical bars show the range (n=2 screen replicates). Dashed horizontal line is the hit calling threshold based on random controls. UniProt annotations are shown below, associated with the UniProt accession ID written above.

**Supplementary Figure 15.**
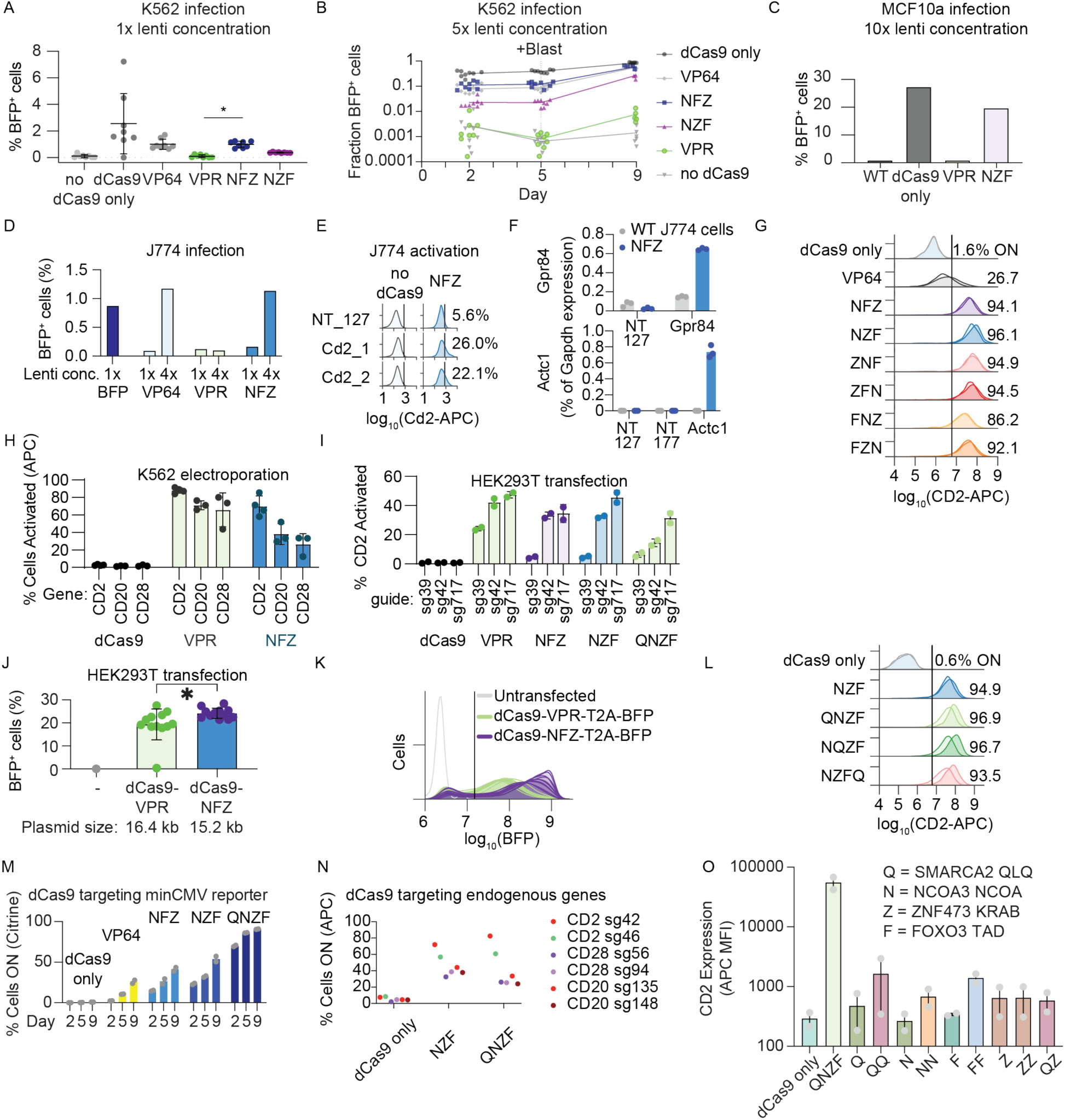
Characterization of transcriptional control tools using compact human activators. **A**. dCas9-activator delivery to K562 cells by lentiviral infection. Flow cytometry measuring BFP delivery marker was performed 3 days after infection (n=8 infection replicates). * denotes p<0.0001 by unpaired t test for comparison of NFZ and VPR. **B**. 5x LentiX-concentrated lentivirus was used and cells were analyzed by flow cytometry at 3 timepoints. Blasticidin selection was initiated on day 5. Each dot is an independent infection (n=8 replicates per dCas9 construct). **C**. Lentiviral delivery to MCF10a breast cancer cells with 10x LentiX concentration, measured by flow cytometry after 4 days (n=1 replicate). **D**. Lentiviral delivery to J774 mouse macrophages with or without 4x LentiX concentration. 1e5 cells were infected with 1 mL of virus for 24 hours. Flow cytometry was performed 6 days after infection. BFP is a positive control lentivirus (pEF-PuroR-P2A-BFP). **E**. Activation of endogenous Cd2 in J774 after lentiviral sgRNA delivery (n=1 infection). Data is gated for sgRNA delivery (GFP^+^) and for fair comparison, not gated for dCas9 delivery. Non-targeting (NT) sgRNAs are negative controls. **F**. Activation of endogenous Gpr84 or Actc1 measured by qPCR after lentiviral sgRNA delivery (n=1 infection). **G**. Optimization of the tripartite activator by changing the N, F, Z domains’ orientation. The various configurations were fused onto dCas9 and targeted to the CD2 gene in K562 cells. sgRNAs and then dCas9 fusions were delivered by unconcentrated lentivirus and selected with puromycin and blasticidin, respectively. Activation was measured 8 days after dCas9 infections by immunostaining CD2 with an APC-conjugated antibody followed by flow cytometry. The average percentage of cells ON is shown. The darker shaded histogram is CD2 sg717 and the lighter shade is sg718 (n=1 infection per sgRNA). dCas9-only and VP64 control data are shared with Figure 1C. **H**. dCas9-activators targeting CD2, CD20, CD28 surface marker genes in K562 cells. sgRNAs were stably installed by lentiviral delivery and puromycin selection. Then 500 ng of dCas9 plasmids were electroporated into 1e6 cells. Two days later, cells were stained for surface CD2 (APC), CD20 (APC), or CD28 (PE) expression and analyzed by flow cytometry after gating for dCas9 (BFP) and the stably expressed sgRNA (GFP). Each dot represents a different sgRNA targeting the gene (n=3-4 per gene). Control data are shared with **Supplementary Figure 1E**. **I**. Percentage of CD2 endogenous gene activated 3 days after transient transfection of dCas9-activators and an sgRNA in HEK293T cells. Cells were immunostained for CD2 (APC) expression and analyzed by flow cytometry after gating for transfection (GFP on the sgRNA plasmid). Each dot represents an independently transfected biological replicate (n=2). **J**. dCas9-activator delivery 2 days after plasmid transfection in HEK293T cells measured by flow cytometry for the BFP marker on the dCas9-effector-T2A-BFP-P2A-BlastR transcript, with no gating on co-transfected delivery markers. dCas9-NFZ is significantly better delivered than dCas9-VPR with a greater fraction of BFP+ cells using a linear gate at 1.5e7 (P<0.05, two-tailed t-test). Each dot is an individual co-transfection of 500 ng of dCas9 plasmid and 300 ng of an sgRNA plasmid in a 24-well plate (n=11 transfection replicates for NFZ and VPR) or an untransfected negative control. Bar shows mean and error bars show standard deviation. **K**. BFP expression level in the same samples after accounting for overall delivery efficiencies by gating for transfectable cells based on the presence of GFP (from the co-transfected sgRNA plasmid). Each line is an individual co-transfection or the untransfected control and the black line shows the linear gate for BFP^+^ cells. **L**. Fusion of SMARCA2 QLQ (Q) to dCas9-NZF. 9 days after lentiviral delivery of dCas9-activators, K562 cells were immunostained with CD2 antibody to measure gene activation by flow cytometry, and the average percentage of cells ON is shown. The darker shaded histogram is CD2 sg717 and the lighter shade is sg718. **M**. dCas9 recruitment of activators with an sgRNA that binds the TetO site upstream the minCMV reporter in K562 cells (n=2 infection replicates). dCas9 fusions were delivered by lentivirus and cells were analyzed 2, 5, and 9 days later with blasticidin selection starting at day 5. Flow cytometry measurements were gated for dCas9 and TetO_sg1 using BFP and mCherry, respectively (VPR measurements not shown due to having <100 BFP^+^ cells). Control data are shared with **Supplementary Figure 4G**. Bars show mean and error bars show standard deviation. **N**. dCas9-activators were delivered to K562 cells by lentivirus and selected for with blasticidin, then sgRNAs were delivered by lentivirus and selected for with puromycin, then 8 days after sgRNA delivery the cells were stained for the targeted surface marker genes and measured by flow cytometry. Surface marker expression is shown after gating for dCas9 with BFP and sgRNA with GFP, and dCas9-only data is shared with **Supplementary Figure 4J**, which was performed in parallel. Top: the darker shaded histogram is CD2 sg42 and the lighter shade is sg46; middle: the darker shade is CD20 sg135 and the lighter shade is sg148; bottom: the darker shade is CD28 sg56 and the lighter shade is sg94. **O**. Homotypic combination of Q, N, Z, F activators fused onto dCas9 and delivered stably by lentivirus to target CD2 in K562 cell lines stably expressing the sgRNA. The mean fluorescence intensity (MFI) of CD2 staining (Alexa 647) of the cell population after gating for delivery of the sgRNA (GFP) and dCas9 fusion (BFP) is shown. Staining was performed 9 days after dCas9 fusion infection. Each point is an sgRNA (sg717 or sg718) and bars show the mean of two different sgRNAs.

**Supplementary Figure 16.**
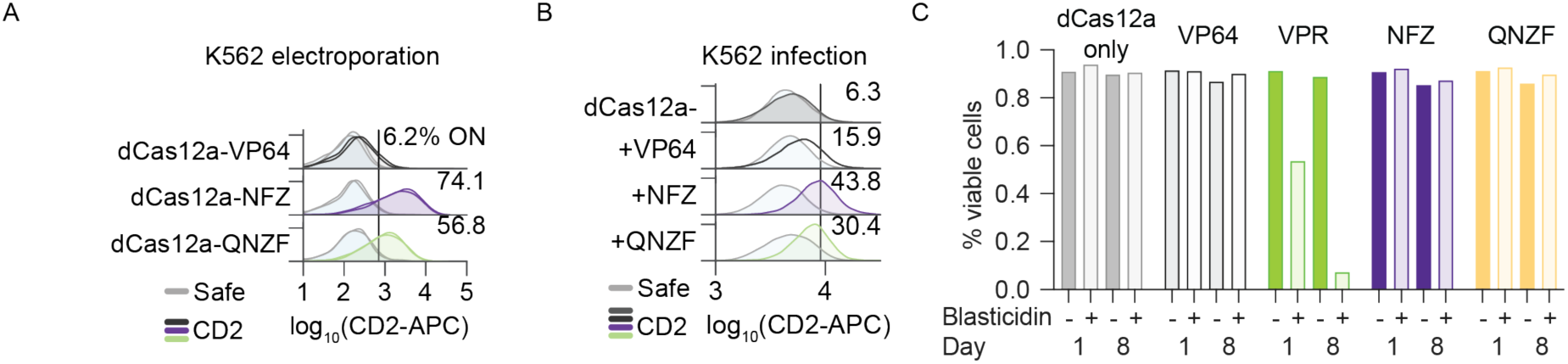
Compact activators are effective when fused to dCas12a. **A**. NFZ and QNZF, or VP64 as a control, were fused to dCas12a. 1 ug of dCas12a plasmid and 1 ug of a CD2-targeting or safe-control gRNA expression plasmid with an mCherry marker was delivered to K562 cells by co-electroporation. After 3 days, cells were stained for CD2 with APC-conjugated antibody and activation was measured by flow cytometry. mCherry-APC bleedthrough compensation was applied and data was gated for high gRNA delivery (mCherry). Two replicates are shown as shaded histograms, and the mean percentage of CD2-targeted and activated cells is shown. **B**. dCas12a-only or dCas12a-activator fusions and then gRNAs were delivered to K562 cells by lentiviral infections. 8 days after the gRNA infection, cells were stained for HA and CD2 and activation was measured by flow cytometry. Data was gated for dCas12a (HA-tag) and gRNA delivery (mCherry). One replicate is shown and the percentage of CD2-targeted and activated cells is shown. **C**. dCas12a fusions were delivered to K562 cells by lentivirus. 5 days later, cells were split into blasticidin treatment or control wells. The percentage of viable cells was determined by flow cytometry 1 or 8 days later (n=1 replicate).

**Supplementary Figure 17.**
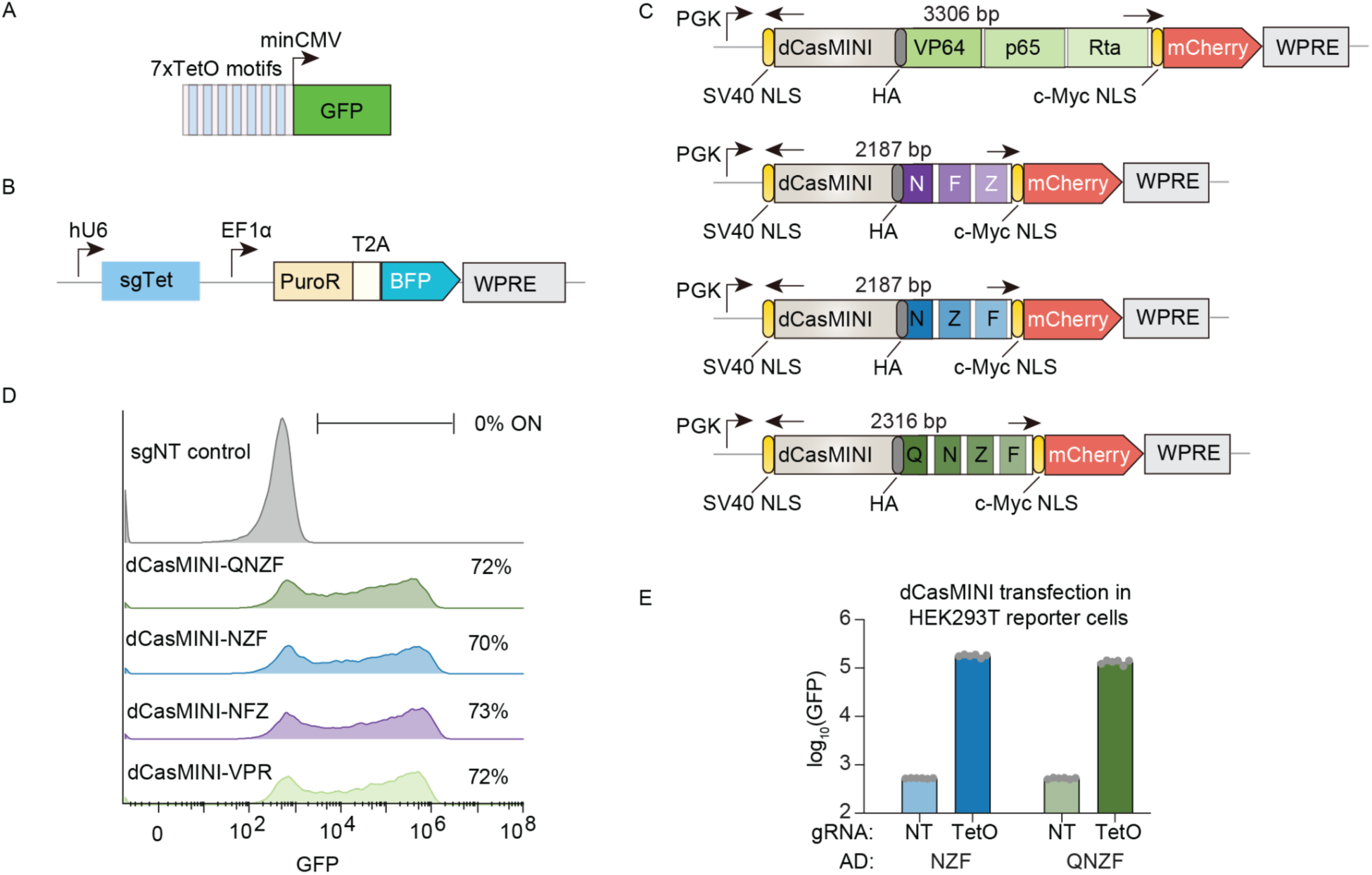
Compact activators are effective when fused to dCasMINI. **A**. Schematic of pTRE3G GFP reporter with 7xTetO motifs and the minCMV promoter upstream, which was integrated in HEK293T cells using lentivirus. **B**. Schematics of construct for expressing the sgRNA that targets the TetO motif. **C**. Schematics of constructs for expressing the dCasMINI fusions. **D**. Representative GFP distributions after dCasMINI and an sgRNA expression plasmid were co-transfected into HEK293T reporter cells. 2 days later, GFP reporter activation was measured by flow cytometry with gates for BFP^+^/mCherry^+^ cells. **E**. Mean GFP fluorescence is shown after gating for sgRNA (BFP) and dCasMINI (mCherry) delivery (n=6 transfection replicates shown as dots). VPR and NFZ data from the same experiment are shown in Figure 4.

**Supplementary Figure 18.**
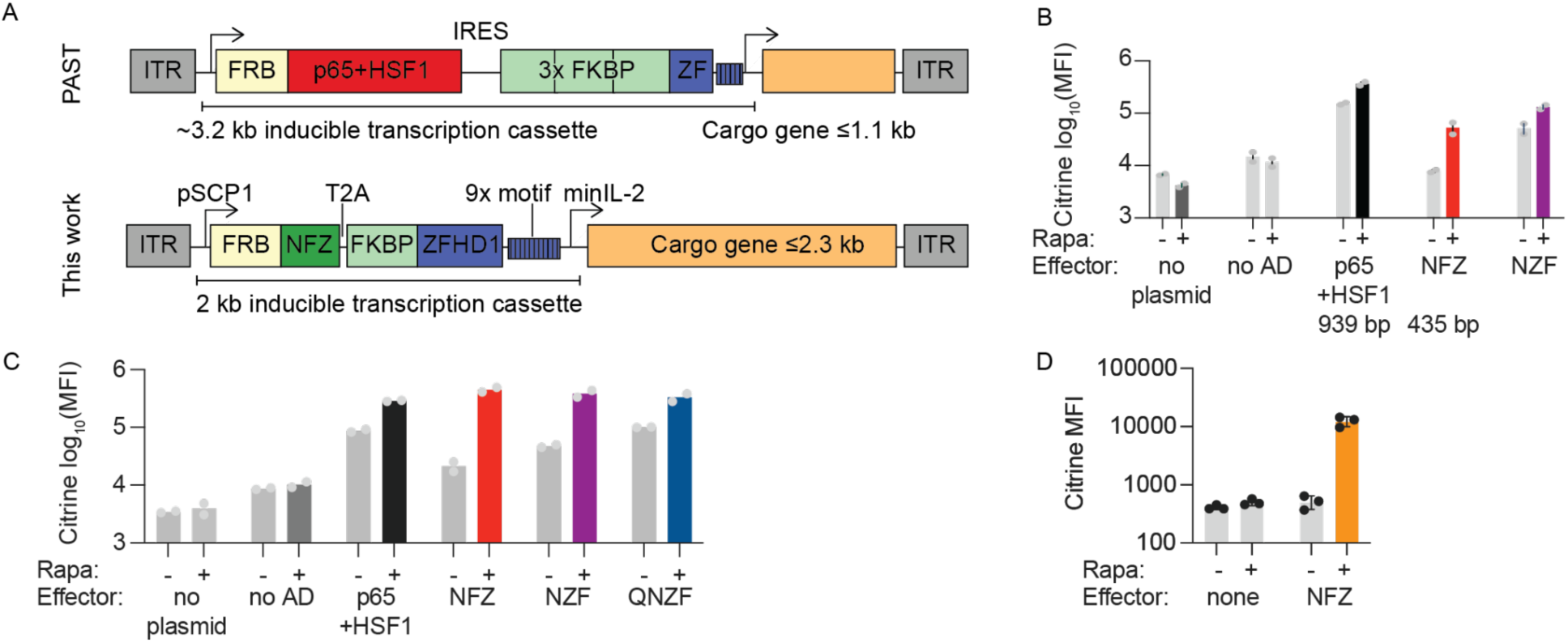
Inducible transgene expression with compact human activator domains. **A**. Schematic comparing the past rapamycin inducible AAV vector design (Rivera et al., 2005) with one using the compact NFZ activator, minimal promoters and terminators (Juven-Gershon et al., 2006; McFarland et al., 2006; Vora et al., 2018), and fewer repeats of FKBP, altogether removing 1.2 kb. If employed in an AAV delivery vector, with a 4.7 kb payload capacity flanked by ITRs, this design would be expected to allow up to 2.3 kb for the cargo gene. **B**. Rapamycin-inducible expression of citrine cargo gene with ZFHD1 recruitment of activators in K562 cells. 2 ug of plasmids were electroporated into cells, 1 day later 100 nM rapamycin was added, and 1 day later citrine mean fluorescence intensity (MFI) was measured by flow cytometry (n=2 electroporation replicates, bar shows mean and error bar shows range). **C**. Rapamycin-inducible expression of citrine with ZFHD1 recruitment activators in HEK293T cells. 1.5 ug of plasmids were transfected into HEK293T cells, one day later 10 nM rapamycin was added, and two days later citrine mean fluorescence intensity (MFI) was measured by flow cytometry (n=2 transfection replicates, bar shows mean). **D**. Rapamycin-inducible expression of reporter citrine gene with ZFHD1 recruitment of activators. Here the plasmid has a bGH polyA for the inducer cassette, whereas a snRP-1 terminator is used in Figure 4J. 3 ug of plasmids were transfected into HEK293T cells, one day later 10 nM rapamycin was added, and two days later citrine mean fluorescence intensity (MFI) was measured by flow cytometry (n=3 transfection replicates, bar shows mean and error bar shows standard deviation).

## Materials and Methods

### Cell lines and cell culture

Experiments presented here were carried out in K562 (ATCC CCL-243, female), HEK293T (Takara Bio #632180), MCF10a (ATCC), and J774 (ATCC) cells. Cells were cultured in a controlled humidifier at 37°C and 5% CO2. K562 cells were cultured in RPMI 1640 (Gibco, 11-875-119) media supplemented with penicillin (10,000 I.U./mL), streptomycin (10,000 μg/ml), 2mM L-glutamine, and 10% Tet Approved FBS (Omega Scientific Lot # 20014T). HEK293T cells were maintained in Dulbecco’s modified Eagle medium (DMEM; Gibco #10569010) supplemented with 25 mM D-glucose (Gibco), 1 mM sodium pyruvate (Gibco), 1x GlutaMAX^TM^ (Gibco), penicillin (10,000 I.U./mL), streptomycin (10,000 μg/ml), and 10% Tet Approved FBS (Omega Scientific Lot # 20014T). When HEK293T cells reached 80% confluence, they were gently washed with 1x DPBS (Life Technologies) and passaged using 0.25% Trypsin (Life Technologies). MCF10a cells were maintained in DMEM/F12 (Invitrogen #11330-032) supplemented with insulin (10 μg/ml), cholera toxin (100 ng/ml), hydrocortisone (0.5 mg/ml), EGF (20 ng/ml), penicillin (10,000 I.U./mL), streptomycin (10,000 μg/ml), and 5% Horse Serum (Invitrogen #16050-122). J774 cells were cultured in 10 cm plates in DMEM media supplemented with 2 mM glutamine, 100 U/ml penicillin, 100 µg/ml streptomycin and 10% heat-inactivated FBS, and were passaged 2-3 times weekly by exchanging media when cells reached ∼90% confluency, incubating for 24 h, and scraping. For long-term storage, cells were resuspended in freezing media (10% dimethyl sulfoxide (Sigma) and cell media) in a cryovial and frozen at −80°C.

### Generating reporter cell lines

The minCMV, pEF1α, RSV, UbC, and PGK promoters were selected to span a range of basal expression levels. To include more minimal promoters with low basal expression, two promoters NTX and NT21 (which both lack a TATA box, in contrast to minCMV) were selected from a published resource (Haberle et al., 2019). These promoters were each cloned into a reporter plasmid with homology arms for integration into the AAVS1 safe harbor locus.

Reporter cell lines were generated by TALEN-mediated homology-directed repair via integration of a donor construct into the AAVS1 locus of cells as follows: 1.2×10^6^ K562 cells were electroporated in Amaxa solution (Lonza Nucleofector 2b, setting T-16) or 8×10^4^ HEK293T cells were transfected using Lipofectamine LTX (Invitrogen), according to manufacturer instructions, with 1000 ng of reporter donor plasmid, and 500 ng of each TALEN-L (Addgene #35431) and TALEN-R (Addgene #35432) plasmid (targeting upstream and downstream the intended DNA cleavage site, respectively). Cells were selected with 500 ng/mL puromycin (InvivoGen) starting 48 h post electroporation/transfection for 5 days or until all of the negative control cells died. The TALEN plasmids were gifts from Feng Zhang (Sanjana et al., 2012). These cells were not authenticated.

### Domain library design and cloning

The nuclear protein Pfam domain library was designed previously (Tycko et al., 2020). Briefly, we queried the UniProt database(UniProt Consortium, 2015) for human genes that can localize to the nucleus. We then retrieved Pfam-annotated domains using the ProDy searchPfam function(Bakan et al., 2011). We filtered for domains that were 80 amino acids or shorter and excluded the C2H2 Zinc finger DNA-binding domains, which are highly abundant, repetitive, and not expected to function as transcriptional effectors. We retrieved the sequence of the annotated domain and extended it equally on either side to reach 80 amino acids total. Duplicate sequences were removed, then codon optimization was performed for human codon usage, removing BsmBI sites and constraining GC content to between 20% and 75% in every 50 nucleotide window using DNA chisel (Zulkower & Rosser, 2020). 499 random controls of 80 amino acids lacking stop codons were computationally generated as controls. 362 elements tiling the DMD protein in 80 amino acid tiles with a 10 amino acid sliding window were also included as controls because DMD was not thought to be a transcriptional regulator. In total, the library consists of 5,954 elements.

The chromatin regulator and transcription factor tiling library was previously designed (DelRosso et al., 2023). Briefly, 735 chromatin regulator and 1294 transcription factor proteins were tiled with 80 amino acid tiles with a 10 amino acid tile sliding window. In addition, 2028 random sequence controls (lacking stop codons) were computationally generated. Duplicate sequences were removed, sequences were codon optimized for human codon usage, 7xC homopolymers were removed, BsmBI restriction sites were removed, rare codons (less than 10% frequency) were avoided, and the GC content was constrained to be between 20% and 75% in every 50 nucleotide window (performed with DNAchisel (Zulkower & Rosser, 2020)). The library includes a total of 128,565 elements (including small sub-libraries that were not analyzed here) which were split across 5 subpools that were separately amplified and cloned into the dCas9 recruitment plasmid pJT216.

Oligonucleotides with lengths of 300 nucleotides to encode these protein sequences and flanking cloning and amplification adapters were synthesized as pooled libraries (Twist Biosciences). They were PCR amplified in 6×50 ul reactions set up in a clean PCR hood to avoid amplifying contaminating DNA. For each reaction, we used 5 ng of template ssDNA oligo pool, 0.1 μl of each 100 μM primer, 1 μl of Herculase II polymerase (Agilent), 1 μl of DMSO, 1 μl of 10 nM dNTPs, and 10 μl of 5x Herculase buffer. The thermocycling protocol was 3 minutes at 98°C, then cycles of 98°C for 20 s, 61°C for 20 s, 72°C for 30 s, and then a final step of 72°C for 3 minutes. The default cycle number was 21x, and this was optimized for each library to find the lowest cycle that resulted in a clean visible product for gel extraction (in practice, 21 cycles was the minimum). For some subpools, the annealing temperature was lowered to 58°C. After PCR, the resulting dsDNA libraries were gel extracted by loading ≥4 lanes of a 2% TBE gel, excising the band at the expected length (∼300 bp), and using a QIAgen gel extraction kit.

The libraries were cloned into a lentiviral dCas9 recruitment vector pJT216 (Addgene #187320) with 4-10x 10 μl GoldenGate reactions (75 ng of pre-digested and gel-extracted backbone plasmid, 5 ng of library (2:1 molar ratio of insert:backbone), 0.13 μl of T4 DNA ligase (NEB, 20000 U/μl), 0.75 μl of Esp3I-HF (NEB), and 1 μl of 10x T4 DNA ligase buffer) with 30-65 cycles of digestion at 37°C and ligation at 16°C for 5 minutes each, followed by a final 5 minute digestion at 37°C and then 20 minutes of heat inactivation at 70°C. The reactions were then pooled and purified with MinElute columns (QIAgen), eluting in 6 ul of ddH2O. 2 μl per tube was transformed into two tubes of 50 μl of Endura electrocompetent cells (Lucigen, Cat#60242-2) following the manufacturer’s instructions. For maximum coverage of the larger subpools, we sought to have 4 ul of ∼200-300 ng/ul library DNA per tube of DUO cells after the minElute step. After recovery, the cells were plated on 3-8 large 10” x 10” LB plates with carbenicillin. After overnight growth at 37°C, the bacterial colonies were scraped into a collection bottle and plasmid pools were extracted with a HiSpeed Plasmid Maxiprep kit (QIAgen). 2-3 small plates were prepared in parallel with diluted transformed cells in order to count colonies and confirm the transformation efficiency was sufficient to maintain at least 30x library coverage.

To determine the quality of the libraries, the domains were amplified from the plasmid pool and from the original oligo pool by PCR with primers with extensions that include Illumina adapters and sequenced. The PCR and sequencing protocol were the same as described below for sequencing from genomic DNA, except these PCRs use 10 ng of input DNA and 17 cycles. These sequencing datasets were analyzed as described below to determine the uniformity of coverage and synthesis quality of the libraries. In addition, 20 colonies from the transformations were Sanger sequenced (Quintara) to estimate the cloning efficiency and the proportion of empty backbone plasmids in the pools. The 5 subpools of the tiling library were pooled at the plasmid level before lentiviral packaging.

We also used previously cloned plasmid libraries wherein these domains were cloned onto the rTetR or rTetR(SE-G72P) mutant version of rTetR with reduced leakyness (DelRosso et al., 2023; Roney et al., 2016; Tycko et al., 2020). In addition, we integrated previous HT-recruit datasets which also used these previously cloned libraries, with rTetR used for the Pfam library at the pEF reporter in K562 cells and rTetR(SE-G72P) used for all other screens.

### Large scale lentiviral delivery

To generate lentivirus for large-scale experiments, HEK293T cells were grown to 70% confluence in 15 cm dishes in 30 mL volume. These were transfected with 10 ug of the transfer plasmid and 10 ug of the packaging plasmid mix, along with 100 uL of PEI in 2 mL of serum-free DMEM. 48 hours after transfection, the viral supernatant was harvested and filtered through a 0.45 um bottle top filter (Thermo Scientific 168-0045), and stored at 4°C. An additional 30 mL of full DMEM was gently added back to the dish. This media was filtered 24 hours later (72 hours after transfection), and combined with the 48 hour viral supernatant.

For screens, K562 cells were transduced in 6-well plates by mixing the cells and virus with 1:500 dilution of 4 mg/mL Polybrene (Sigma H9268-5G). For spinfection, 5-6 mL of the mix was plated in each well of a 6-well plate, then the plates were centrifuged at 1,000x g at 33C for 2 hours in covered buckets, before removing the viral supernatant and resuspending the transduced K562 cells in full RPMI media. For screens in HEK293T cells, cells were either transduced by spinfection or incubated overnight with virus and polybrene in T225 flasks, which was then aspirated and replaced with DMEM media.

### HT-recruit to measure repressor activity at reporters

For the nuclear Pfam domain repressor screens, 4 × 10^7^ K562 PGK or UbC reporter cells per replicate were infected with 72 mL of the lentiviral library by spinfection. Cells were started in T150 flasks. 2 days later, cells were ∼10% mCherry^+^ by flow cytometry (ZE5 Cell Analyzer), and selection began with 10 μg/mL blasticidin (InvivoGen) in T225 flasks. Cells were maintained in log growth conditions each day by diluting cell concentrations back to a 5 × 10^5^ cells/mL, with at least 1 × 10^8^ cells total remaining per replicate such that the lowest maintenance coverage was >10,000× cells per library element. On day 7 post-infection, recruitment was induced by treating the cells with 1000 ng/mL doxycycline (Tocris) for 5 days then cells (>5000x coverage) were harvested for magnetic separation.

HEK293T screens were performed in T225 flasks and split daily via dissociation with 1x TrypLE (Gibco). For HEK293T cells with the pEF reporter, 1.25e7 cells per replicate were infected by spinfection with 25 mL of virus in 6-well plates with 5 mL per well, resulting in 85% mCherry^+^ two days later. 2 days after infection, 1 ug/ml doxycycline and 10 ug/ml blasticidin were added. 4 days later, 75% of cells (∼6,500x coverage) were harvested for magnetic separation, and the remaining cells were washed twice and resuspended in fresh media. 8 days later, those cells (10,000x coverage) were harvested for magnetic separation.

For HEK293T cells with the UbC reporter, 1.65e7 cells per replicate were infected by overnight incubation with varied doses of virus in T225 flasks. The screen continued with the high dose (4.13 mL) resulting in ∼24% mCherry^+^ cells two days later. 2 days after infection, 10 ug/ml blasticidin was added. 5 days later the cells were frozen. Later, cells were thawed, then blasticidin was added and refreshed daily for 6 days. Then, 1 ug/ml doxycycline was added and 4 days later, cells (>6,500x coverage) were harvested for magnetic separation.

### HT-recruit to measure activation activity at reporters

For the nuclear Pfam domain activator screens, 4 × 10^7^ K562 NTX or NT21 reporter cells per replicate were infected with 72 mL of the lentiviral library by spinfection. Cells were started in T150 flasks. 2 days later, cells were ∼10% mCherry^+^ by flow cytometry (ZE5 Cell Analyzer), and selection began with 10 μg/mL blasticidin (InvivoGen) in T225 flasks. Cells were maintained in log growth conditions each day by diluting cell concentrations back to a 5 × 10^5^ cells/mL, with at least 1 × 10^8^ cells total remaining per replicate such that the lowest maintenance coverage was >10,000× cells per library element. On day 7 post-infection, recruitment was induced by treating the cells with 1000 ng/mL doxycycline (Tocris) for 2 days then cells (>6500x coverage) were harvested for magnetic separation.

The protocol was similar for HEK293T cells, but cells were maintained daily in a T225 flask at >1.4 × 10^7^ via dissociation with 1x TrypLE (Gibco). For HEK293T cells with the minCMV and silenced PGK reporters, 1.65e7 cells per replicate were infected by overnight incubation with varied doses of virus in T225 flasks. The screen continued with the high dose (4.13 mL) resulting in ∼24% mCherry^+^ cells two days later. 2 days after infection, 10 ug/ml blasticidin was added. The minCMV screen continued: 5 days later 1 ug/ml doxycycline was added and 2 days later the cells (>2,000x coverage) were harvested for magnetic separation. Meanwhile, 5 days after blasticidin addition, PGK cells were frozen. Later, PGK cells were thawed, then blasticidin was added and refreshed daily for 6 days. Then, 1 ug/ml doxycycline was added and 2 days later, cells (>6,500x coverage) were harvested for magnetic separation.

### CRISPR HT-recruit to measure transcriptional effectors at endogenous genes and reporters

CRISPR HT-recruit screens were performed with dCas9 as the DBD and an sgRNA targeting either an endogenous surface marker (CD2, CD20, CD28, or CD43) or the TetO site upstream a reporter with the synthetic surface marker (TetO-minCMV or TetO-pEF1α). First, the sgRNA was stably delivered to K562 cells by lentivirus and selected for with puromycin for 3-4 days. The cells were confirmed to be >95% mCherry^+^ by flow cytometry (BD Accuri C6).

For the first replicate of the CD2 (sg717, sg718) and CD43 screens, 1.35 × 10^7^ of these cell lines per replicate were infected with 20 mL of the 4x-concentrated (LentiX) lentiviral dCas9-Pfam library by spinfection (split and spun in 2x 10mL volumes in 50 mL Falcon tubes). The cells were ∼10% BFP^+^ by flow cytometry (ZE5) after 5 days (as measured from a set-aside plate that did not receive antibiotic selection). For the CD20, CD28, other CD2 sgRNAs, and the second replicate of the CD2 sg717 screens, 1.35 × 10^7^ of these cell lines per replicate were infected with 20 mL of the 4x-concentrated (LentiX) lentiviral dCas9-Pfam library by spinfection in 4 mL per well in 6-well plates. The cells were ∼68% BFP^+^ by flow cytometry (ZE5) after 5 days. For the TetO sgRNA and the second replicate of the CD43 screens, 1.5e7 cells were spinfected with 9 mL of 4x-concentrated virus split in 3 wells of a 6-well plate, resulting in ∼50% BFP^+^ cells.

For all of these samples, cells were maintained in T175 flasks. 2-3 days after infection, selection began with 10 μg/mL blasticidin (InvivoGen). Cells were maintained in log growth conditions each day by diluting cell concentrations back to a 5 × 10^5^ cells/mL (and replenishing blasticidin), with at least 4.2 × 10^7^ cells total remaining per replicate such that the lowest maintenance coverage was >5,000× cells per library element. On day 10 post-infection, cells (>20,000x coverage) were harvested for magnetic separation.

For the dCas9-CRTF screens, lentivirus for the library was generated using 16x 15 cm dishes of HEK293T cells and then concentrated 4x using LentiX. Then 1.15 × 10^8^ K562-sgRNA cells per replicate were infected with 72 mL of the lentiviral library by spinfection for 2 hours, with two separate biological replicates of the infection, resulting in 18-23% BFP+ cells in unselected cells after 4 days. 2 days after infection, the cells were selected with 10 μg/mL blasticidin (InvivoGen). Cells were >95% BFP^+^ by the final timepoint. On day 11 post-infection, 5 × 10^8^ cells (>3,000x coverage) were harvested for magnetic separation.

An additional growth-based CRISPR HT-recruit screen was performed with the Pfam domain library and an sgRNA targeting the eGATA1 enhancer (which has a fitness effect when repressed) or a safe-targeting negative control with no fitness effect. For the first replicate, 1.35 × 10^7^ of these cell lines per replicate were infected with 20 mL of the 4x-concentrated (LentiX) lentiviral dCas9-Pfam library by spinfection (split and spun in 2x 10mL volumes in 50 mL Falcon tubes). Cells were maintained in T175 flasks. The cells were ∼10% BFP^+^ by flow cytometry (ZE5) after 5 days (as measured from a set-aside plate that did not receive antibiotic selection). 3 days after infection, cells were selected with 10 μg/mL blasticidin (InvivoGen) for 7 days. The second replicate was performed later, with 1.5e7 cells spinfected with 9 mL of virus split in 3 wells of a 6-well plate, resulting in ∼60% BFP^+^ cells. Cells were maintained in log growth conditions each day by diluting cell concentrations back to a 5 × 10^5^ cells/mL (and replenishing blasticidin), with at least 4.2 × 10^7^ cells total remaining per replicate such that the lowest maintenance coverage was >5,000× cells per library element. On day 13 post-infection, cells (>20,000x coverage) were harvested for genomic DNA extraction and sequencing.

### Magnetic separation of HT-recruit screens

Magnetic separation of HT-recruit screen cells were performed for all screens targeting a reporter or surface marker gene. Cells were spun down at 300 x g for 5 minutes and media was aspirated. Cells were then resuspended in the same volume of PBS (Gibco) and the spin down and aspiration was repeated, to wash the cells and remove any IgG from serum. Dynabeads M-280 Protein G (ThermoFisher, 10003D) were resuspended by vortexing for 30 s. 50 mL of blocking buffer was prepared per 2 x 10^8^ cells by adding 1 g of biotin-free BSA (Sigma Aldrich) and 200 mL of 0.5 M pH 8.0 EDTA into DPBS (Gibco), vacuum filtering with a 0.22-um filter (Millipore), and then kept on ice. For activation reporter screens, 30 uL of beads was prepared for every 1 x 10^7^ cells. For repression reporter screens, 90-120 uL of beads was prepared for every 1 x 10^7^ cells. This volume varies to optimize cost and separation purity: if the cell population is mostly ON, more beads provides greater purity. Beads were prepared by resuspension in the vial by vortexing for 30 seconds, then adding 1 mL of buffer per 200 ul of beads, vortexing for 5 seconds, and putting the tube on the magnetic stand. After 1 minute, supernatant was removed, the tube was removed from the stand and beads were resuspending in the desired amount of blocking buffer (default was 12 mL buffer for a screen with >50M cells to be separated in a 15 mL tube on a large magnetic stand). We added beads to cells at no more than 10 million cells per 100 uL of buffer-bead mixture and incubated the mixture at room temperature while rocking for at least 30 minutes. We then put the mixture on a magnetic stand for >2 minutes. We saved the Unbound supernatant in a new tube and placed that tube on the magnet again for >2 min to remove any remaining beads. We moved this supernatant to a new tube and kept it as the Unbound fraction and recorded its volume. Meanwhile, we resuspended all the beads in the same volume of blocking buffer, and magnetically separated them again to help get a clean ON population, then set aside the “Bound wash off” supernatant (for flow analysis before disposal), while keeping the tube with beads as the Bound fraction. We resuspended the Bound fraction in blocking buffer (or PBS) and recorded its volume. We then used flow cytometry to measure the cell concentration in the Bound and Unbound fractions and confirm the separation of ON and OFF cells. Finally, we spun down the cells and froze the two pellets (Bound and Unbound) at –20C for subsequent genomic DNA extraction.

For dCas9 HT-recruit screens targeting reporters with the TetO sgRNA, the cells were similarly directly separated with 90 ul or 60 ul of magnetic beads per 10M cells for the pEF and minCMV reporters, respectively. For endogenous gene-targeting dCas9 HT-recruit screens, cells were stained with antibodies against the target surface marker before magnetic separation. Cells were first washed with 1% BSA (Sigma) in 1× DPBS (Life Technologies) and spun down and supernatant was aspirated without disturbing the pellet. 2-5 mL of cells were then incubated on ice for 1 h with fluorophore conjugated primary antibody. The following primary antibodies were used: allophycocyanin (APC)-labeled anti-CD2 antibody (1:50 dilution, 130-116-253, Miltenyi-Biotec), APC-labeled anti-CD43 antibody (1:500, clone 4-29-5-10-21, eBioscience, Catalog # 17-0438-42), APC-labeled anti-CD20 antibody (1:100, clone L26, eBioscience, Catalog # 14-0202-82), or APC-labeled anti-CD28 antibody (1:20, clone CD28.2, eBioscience, Catalog # 17-0289-42). Afterwards, cells were washed with 45 mL of 1% BSA/DPBS. They were then magnetically separated with 30-60 ul of Protein G Dynabeads per 10M cells.

### Pooled measurements of FLAG-tagged DBD-domain fusion expression levels

The expression level measurements were made in cells infected with the 3XFLAG-tagged nuclear Pfam domain library. For dCas9 in K562 cells, screen cells with safe N4293 and CD43 sg15 were frozen 9 days post-infection, then later thawed and 1.6-3e7 cells were harvested for FLAG staining. For rTetR in HEK293T cells, the minCMV reporter cell screen was used, and 1.6-3e7 cells (>600x coverage) were harvested 6 days after blasticidin addition (with no doxycycline), which was 8 days post-infection.

Fix Buffer I (BD Biosciences, BDB557870) was preheated to 37°C for 15 minutes and Permeabilization Buffer III (BD Biosciences, BDB558050) and PBS (GIBCO) with 10% FBS (Hyclone) were chilled on ice. The library of cells expressing domains was collected and cell density was counted by flow cytometry (ZE5 Cell Analyzer). To fix, cells were resuspended in a volume of Fix Buffer I (BD Biosciences, BDB557870) corresponding to pellet volume, with 20 μL per 1 million cells, at 37°C for 10 – 15 minutes. Cells were washed with 1 mL cold PBS containing 10% FBS, spun down at 500 x g for 5 minutes and then supernatant was aspirated. Cells were permeabilized for 30 minutes on ice using cold BD Permeabilization Buffer III (BD Biosciences, BDB558050), with 20 μL per 1 million cells, which was added slowly and mixed by vortexing. Cells were then washed twice in 1 mL PBS+10% FBS, as before, and then supernatant was aspirated. Antibody staining was performed for 1 hour at room temperature, protected from light, using 5 μL / 1 x 10^6^ cells of α-FLAG-Alexa647 (RNDsystems, IC8529R). We then washed the cells and resuspended them at a concentration of 3 x 10^7^ cells / mL in PBS+10%FBS. Cells were sorted into two bins based on the level of APC-A fluorescence (Sony SH800S) after gating for mCherry^+^ viable cells. A small number of unstained control cells was also analyzed on the sorter to confirm staining was above background. After sorting, the cellular coverage ranged from 355 – 1,364 cells and 55 – 350 cells per library element across FLAG High and Low samples for the dCas9 and HEK293T screens, respectively. The sorted cells were spun down at 500 x g for 5 minutes and then resuspended in PBS. Genomic DNA extraction was performed following the manufacturer’s instructions (QIAgen Blood Maxi kit was used for samples with >1 x 10^7^ cells, and QIAamp DNA Mini kit with one column per up to 5 x 10^6^ cells was used for samples with ≤1 x 10^7^ cells) with one modification: the Proteinase K + AL buffer incubation was performed overnight at 56°C.

### Library preparation to sequence domains

For domain screens (HT-recruit with magnetic separation or growth readouts, and FLAG-based expression measurements), sequencing libraries were prepared as follows. Genomic DNA was extracted with the QIAgen Blood Maxi Kit following the manufacturer’s instructions with up to 1 x 10^8^ cells per column. DNA was eluted in EB and not AE to avoid subsequent PCR inhibition. The DNA was measured on a NanoDrop. We checked if the 260/230 ratio was >2 as low ratios (e.g <1.5) are indicative of salt contamination from the extraction process that can inhibit the PCR. For samples with low ratios, we diluted the sample with NF-H2O by 4x before the PCR. Genomic DNA was diluted to 200 ng/uL for samples with concentrations above that.

The domain-encoding DNA sequences were then amplified by PCR with primers containing Illumina adapters and sample indices as extensions. A test PCR was performed using 5 ug of genomic DNA in a 50 uL (half-size) reaction to verify if the PCR conditions would result in a visible band at the expected size for each sample. Then, 12-48x 100 uL reactions were set up on ice (in a clean PCR hood to avoid amplifying contaminating DNA), with the number of reactions depending on the amount of genomic DNA available in each experiment. 10 ug of genomic DNA, 0.5 mL of each 100 mM primer, and 50 mL of NEBnext Ultra 2x Master Mix (NEB) was used in each reaction. The thermocycling protocol was to preheat the thermocycler to 98C, then add samples for 3 minutes at 98C, then an optimized number of cycles of 98C for 10 s, 63C for 30 s, 72C for 30 s, and then a final step of 72C for 2 minutes. The default number of cycles was 33x. All subsequent steps were performed outside the PCR hood. The PCR reactions were pooled and 145 uL were run on a 2% TAE gel, the library band ∼422 bp was cut out, and DNA was purified using the QIAquick Gel Extraction kit (QIAgen) with a 30 ul elution into non-stick tubes (Ambion). These libraries were then quantified with a Qubit HS kit (Thermo Fisher). Libraries were pooled with 10-20% PhiX. Single end sequencing with 115-250 cycles was performed on an Illumina NextSeq or by Admera Health on a NovaSeq or HiSeq platform. Each sequencing run contained samples amplified with ≥4 distinct forward primers as they add staggers of various length that avoid sequencing issues in the constant region of the amplicon.

### HT-recruit analysis

The HT-recruit-Analyze pipeline (https://github.com/bintulab/HT-recruit-Analyze) was used to run Bowtie to align Illumina sequencing reads to an index of domain-encoding sequences and compute effector enrichment scores. Sequencing reads were trimmed to remove the 19-24 bp stagger plus constant primer handle region from the beginning of the read. For most samples, reads were further trimmed to retain only the first 131 bp of the domain– encoding sequence. Alignment parameters were optimized for high alignment rate (>80%) and low ambiguous alignment rate (<1%). With these metrics, for the Pfam library, mismatch tolerance was set to Bowtie –m 3 (as each domain is codon optimized so they can differ at more DNA basepairs than amino acid residues), while for the tiling library no mismatches were allowed. This mismatch tolerance approach is tailored towards ignoring sequencing errors while removing reads with synthesis or PCR errors because it allows mismatches if the sum of their Q scores is <30 and the number of mismatches in the initial 28 bp is less than-m.

Then, the log2(OFF:ON) score was computed using sample depth-normalized counts in the OFF (unbound) and ON (bound) samples. To avoid inaccurate measurements from low coverage, if an element had fewer than 5 counts in one sample (bound or unbound fraction), its count was set to 5. If an element falls below that threshold in both samples, it was removed from the data tables. For most downstream analyses, the Pfam and tiling library were further filtered for elements with >5 or >20 counts, respectively, in both samples, as described in the figure legends.

Analysis was performed similarly for the FLAG-based expression level measurements, using the FLAG High and Low samples. An element was defined as well-expressed if its log2(FLAG High:Low) score was greater than 1 standard deviation above the median of the random controls.

Then, an element from the Pfam library was defined as an activator or repressor hit if its log2(OFF:ON) score was 2 standard deviations below (for activators) or above (for repressors) the median of the poorly-expressed domains (as defined by FLAG measurements with that DBD and cell-type). This approach provided a conservative hit threshold that was chosen to identify robust effectors but sub-threshold domains may also have some effector activity. For some analyses of the pEF1α rTetR screen in K562 cells, we also report a lower threshold (0.9), equivalent to the strength of the weakest repressor (DUF1087) that was individually validated (Tycko et al., 2020). The tally of domains that are hits in both replicates of a screen excludes growth-based screens, screen memory timepoints after doxycycline washout, and screens with one replicate (sgCD2_718, sgCD20_135, sgCD20_148, sgCD20_275, sgCD28_07, sgCD28_56, sgCD28_94, sgCD2_39, sgCD2_42, sgCD2_89).

For the tiling library recruited to CD2 or CD43, hits were defined as tiles with scores >2 standard deviations below (activators) or above (repressors) the median of the random controls. Most downstream analyses use tiles that are hits in both replicates. Then, hit domains were called from overlapping hit tiles (that are past the hit threshold in both replicates). A hit domain started anywhere the previous tile was not a hit and ended anywhere the next successive tile was not a hit. Domains started at the first residue of the first tile and extended until the last residue of the last tile within the domain. In addition, the minimized hit domain sequence was defined as the minimal region wherein all overlapping tiles are hits. (For the integrated data from the rTetR recruitment screens with the tiling library, the previously computed hit thresholds, which take into account FLAG-staining data, were used here (DelRosso et al., 2023)). These are 2 standard deviations below the mean of the poorly expressed random controls for minCMV and, more conservatively due to the large amount of signal, 3 standard deviations above the mean of the poorly expressed random controls for pEF and PGK. We did not FLAG-stain the dCas9 tiling library, but would expect the great majority of random controls to be poorly expressed.)

Protein annotations were downloaded from UniProt’s Family and Domain section. Interaction domains were defined as any annotation that contained either “Leucine-zipper”, “dimer”, “bHLH”, or “Interaction with” (and not “Interaction with DNA”). We defined domains that overlapped annotated effector domains as those that overlapped annotations containing any of the following terms: “Repress”; “Negative regulat”; “Inhibition”; “Activ”; “9aaTAD”.

### CRISPRi effector benchmarking growth screens

Two CRISPRi benchmarking screens using growth readouts were performed with different libraries. One is the endogenous repression benchmarking library of 7600 sgRNAs targeting growth genes, enhancers, and safe-targeting and non-targeting controls. This library was used with ZNF10 KRAB and its enhanced mutant as well as combinations with MGA and RYBP. The top 50 essential genes in K562 cells as defined by casTLE analysis (Morgens et al., 2016) of a CRISPRi screen(Gilbert et al., 2014) plus the essential gene MYC, and the bottom 10 least essential genes were selected. For each of these genes, 50 TSS-targeting sgRNAs were retrieved from Dolcetto (Sanson et al., 2018). The library also included sgRNAs tiling the locus of the essential gene GATA1 and targeting the DNase-hypersensitive sites near MYC, and sgRNAs associated with off-target toxicities in K562 cells with CRISPRi (Tycko et al., 2019), that were not included in analysis here.

The second library is called the endogenous repression batch tiling sgRNA library (Yao *et al*., *in review*). It includes 13592 sgRNAs targeting growth gene promoters and enhancers, and safe-targeting negative control sgRNAs. Here, growth genes are defined as genes within the top 500 growth genes in a previously published screen which used CRISPRi with ZNF10 KRAB (Gilbert et al., 2014). New sgRNAs were designed to target these genes, using GuideScan (Perez et al., 2017), so this library mostly does not overlap with the library used to define these growth genes. sgRNAs targeting promoters of growth genes were included for analysis here.

Oligonucleotide libraries encoding the sgRNAs were synthesized by Agilent and then cloned into an sgRNA expression vector pMCB320 that had been cut with BstXI and BlpI restriction enzymes, by ligation with T4 ligase (NEB M0202M).

Then, we used lentivirus from a 15 cm dish of HEK293T packaging cells to infect two replicates each of 10 million wildtype K562 cells to generate stable cell lines expressing various dCas9-repressor fusions with a BFP marker and blasticidin resistance (MOI=0.3). Screens were maintained in 60 ml of media in T175 flasks. 2 days after infections, the cells were selected with 10 μg/mL of blasticidin for 6 days. All lines were confirmed to be >90% BFP+ cells by flow cytometry (Attune). Then, 2 days later, we delivered the library of sgRNAs by lentivirus spinfection at a MOI=0.3-0.5 in 2 infection replicates to achieve >600x infection coverage. 3 days later, we selected for the sgRNAs with 1 μg/mL puromycin for 3 days, then continued passaging cells at ≥1000x maintenance coverage (or ≥5000x for the batch tiling screen) for a total of 14 days since sgRNA infection. Finally, we extracted genomic DNA using the QIAamp DNA Blood Maxi kit (Qiagen 51194).

The genomic integrated sgRNA libraries were amplified and indexed in two rounds of PCR using the Herculase II Fusion DNA Polymerase kit (Agilent Technologies 600679). The libraries were sequenced on an Illumina NextSeq with a custom primer oMCB1672 for read 1 and ≥21 cycles in read 1. The custom primer was added by pipetting 4.5 uL of 100 uM primer into the ∼1.5 mL of liquid in well 20 of the sequencing kit (Read 1 default primer well) and not selecting the custom primer option in the Illumina software. We used the makeCounts and makeRhos functions of the CasTLE package to align the reads to the sgRNA library and compute a foldchange between the initial plasmid pool and final genomic DNA. If an sgRNA had <10 counts in one sample then that count was set to 10, but if it fell below that threshold in both samples then the sgRNA was removed.

### Staining surface markers in individual recruitment assays

For individual experiments analyzing the expression of an endogenous surface marker gene, cells were first washed with 1% BSA (Sigma) in 1× DPBS (Life Technologies) and spun down and supernatant was aspirated without disturbing the pellet. If adherent, cells were dissociated with 1x TrypLE (Gibco) before washing. 100 ul of cells were then incubated on ice for 1 h with fluorophore conjugated primary antibody. The following primary antibodies were used: 10 ul of allophycocyanin (APC)-labeled anti-CD2 antibody (clone LT2; Invitrogen, Catalog # MA1-10132), 5 uL of APC-labeled anti-CD81 antibody (clone 1D6, eBioscience, Catalog # 17-0819-42), 5 ul of APC-labeled anti-CD43 antibody (clone 4-29-5-10-21, eBioscience, Catalog # 17-0438-42), 5 uL of APC-labeled anti-CD32 Monoclonal Antibody (clone 6C4, eBioscience, Catalog # 17-0329-42), 5 uL of APC-labeled anti-CD20 antibody (clone L26, eBioscience, Catalog # 14-0202-82), or 5 uL of APC-labeled anti-CD28 antibody (clone CD28.2, eBioscience, Catalog # 17-0289-42). Afterwards, cells were washed 1-3 times with 1x PBS or 3 times with 1% BSA/DPBS and then analyzed by flow cytometry. For J774 cells, APC anti-mouse Cd2 Antibody (BioLegend 100112, 1:100 dilution) was used.

### Individual lentiviral infections

For some individual experiments, recruitment fusion proteins and guide RNAs were delivered by lentivirus. To generate lentivirus for small-scale experiments, HEK293T cells (ATCC) were grown to 70% confluence in 6-well plates in 2 mL. These cells were transfected by combining 1000 ng of the transfer plasmid and 1000 ng of an equal weight mix of three packaging plasmids (pCMV-VSV-G, pRSV-REV, and pCMV-MDL), along with 10 uL of 1 mg/mL PEI (Polysciences 24765-1) in 200 uL of serum-free DMEM. 48 hours after transfection, the viral supernatant was harvested and filtered through a 0.45 um syringe filter (Millipore SLHV033RB). In some cases lentivirus was concentrated using the LentiX Concentrator (Takara) according to manufacturer’s instructions, with an overnight incubation at 4°C.

To infect K562 cells, 250k cells were mixed with 1 ml of virus and 8 ug/mL polybrene in a 24-well plate or 15 mL conical tube, then spun for 2 hours at 1000 x g at 33C. The cells were resuspended and spun down in a 15 mL tube, then the virus was removed from the pellet. The cell pellet was resuspended in 2 mL media and the cells were grown in a 24-well plate. To infect MCF10a cells, 300k cells were seeded per well of a 6-well plate, and were then infected the next day with 1 mL of concentrated virus supernatant, with 2 mL of fresh media and polybrene (8 μg/ml). The cells were incubated overnight and the media was refreshed the day after infection. To infect J774 cells with dCas9-NFZ, 1e5 cells were incubated in 1 mL virus (either unconcentrated or concentrated with LentiX following the manufacturer’s instructions) for 24 h. A BFP-expressing lentivirus (pSLQ1371) was used as a positive control for transduction. To infect HEK293T cells, 100k cells were plated in a 24-well plate and infected with 100 uL of virus in 1 mL media by overnight incubation.

### Individual activator or repressor recruitment assays with rTetR

Individual effector domains were cloned as fusions with rTetR(SE-G72P), upstream of a T2A-mCherry-BSD marker using GoldenGate cloning into backbones pJT126 (Addgene #161926). For validations, each of the individual rTetR-effector plasmids was stably integrated into the respective reporter line by lentivirus at a MOI of approximately 0.3. Cells were then selected with 10 µg/mL blasticidin (InvivoGen) starting 48 h post transduction until >80% of the cells were mCherry^+^ (6-7 days). When we attempted to use lentivirus to deliver rTetR-VPR (4.6 kb between cPPT and WPRE) to either K562 or HEK293T cells, the cell lines did not survive selection, (while parallel efforts with rTetR-VP64 and others were successful).

These stable cell lines were split into separate wells of a 24-well plate and either treated with doxycycline (Tocris) or left untreated until analyzed by flow cytometry. Cells were assayed by flow cytometry after 5 days of 1 µg/mL doxycycline (Tocris) treatment for repressors and 2 days of doxycycline for activators. In experiments with a post-doxycycline memory measurement, after 5 days of treatment, doxycycline was removed by spinning down the cells, replacing media with DPBS (Gibco) to dilute any remaining doxycycline, and then spinning down the cells again and transferring them to fresh media. Time points were measured every 2-3 days by flow cytometry analysis of >10,000 cells (BioRad ZE5). For experiment with BRM014 (MedChemExpress #HY-119374), cells were co-treated with 1 uM BRM014 during 2 days of doxycycline treatment.

To calculate the fraction of cells silenced (OFF) or activated (ON) during doxycycline treatment, a manual gate (vertical dashed line on flow plots) was imposed on the Citrine reporter fluorescence to determine the percentage of OFF or ON cells for each sample. The gate was selected to contain 1–5% of the Citrine signal in untreated cells. Using the time-matched untreated control, the following equations were used to calculate the background normalized fraction of cells silenced (OFF) or activated (ON):

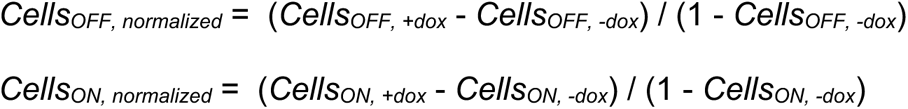

For calculating the fraction of cells super-activated, the same equation and gating was used as in *CellsON, normalized* but in the context of the pEF1α-Citrine reporter.

### Individual recruitment assays with transient transfections of dCas9 in HEK293T cells

Individual effector domains were cloned as fusions with dCas9, upstream of a T2A-mTagBFP2-P2A-BSD marker, using BsmBI GoldenGate cloning into backbone pJT216. Approximately 80,000 HEK293T cells were seeded per well in a 24-well plate and the next day cells were transfected using Lipofectamine LTX (Invitrogen) according to manufacturer instructions. A maximum of 1000 ng of DNA vector was used per transfection. 600 ng of dCas9-effector and 400 ng of sgRNA were co-delivered. For experiments involving dCas9-repressors in HEK293T cells, cells were analyzed 5 days post transfection, and in some cases after being sorted (Sony SH800S sorter) for silencing (TagRFP-negative cells). On the day of flow cytometry analysis, HEK293T cells were collected using 0.25% Trypsin (Life Technologies). A fraction of the HEK293T cells (varying between one half to one twentieth, depending on cell density) were re-plated for the next time point. The remaining cells were resuspended in flow buffer (1x Hank’s balanced salt solution (Life Technologies) and 2.5 mg/mL bovine serum albumin (BSA) (Sigma)) and filtered through a 40 μm strainer (Corning) to remove cell clumps. Cellular fluorescence distributions were measured with the ZE5 Cell Analyzer (Bio-Rad Laboratories) and the Everest Software (Bio-Rad Laboratories).

### Individual recruitment assays with lentiviral delivery of dCas9-repressors

For dCas9 validations of repressors at endogenous genes in K562 cells, individual effector domains were cloned into the pJT216 lenti pEF-dCas9-3XFLAG-Effector-T2A-BFP-P2A-Blast vector, then transfected into HEK293T cells to package lentivirus. After transduction, the K562 cells were selected with 10 μg/ml Blasticidin for 7 days to attain >80% BFP^+^ cells. Lentiviral sgRNAs with mCherry and puromycin selection marker were delivered to these cell lines by spinfection. 2 days after transduction, these cells were selected with 1 μg/ml puromycin for 6 days. The cells were then recovered in fresh media for 1 day. After recovery, the cells were stained with APC-conjugated surface marker antibodies and gated on dCas9-effector-BFP and guide-mCherry expressing cells to quantify activation or repression using flow cytometry (Attune).

Experiments were performed targeting KRAB and mutant KRAB repressors, or non-KRAB repressors and combinations of repressors, to endogenous CD81 in K562 cells. First, the dCas9 fusions (pJT216, 241, 242 in one experiment, pJT216, 242, 244, 250, 341, 338, 340 in another) were stably delivered by lentivirus to K562 cells and then the cells were selected with blasticidin until they were >70% BFP^+^. Then, the CD81 sg3 sgRNA was delivered by lentivirus, three days later puromycin was added to select for the sgRNA, and 9 days post-infection of the sgRNA the cells were stained for CD81 expression and fixed for flow cytometry analysis (BioRad ZE5).

Experiments were performed targeting KRAB and mutant KRAB repressors, or non-KRAB repressors and combinations of repressors, to endogenous CD43 in K562 cells. Cell lines stably expressing the sg10 or sg15 sgRNA (cloned into pMCB320) were first generated using lentiviral delivery, followed by puromycin selection 2 days later for 4 days. These cell lines were then infected with dCas9-repressors lentiviruses, three days later blasticidin selection was initiated, then cells stained for analysis by flow cytometry 9 or 10 days after dCas9 infection (BioRad ZE5). In some cases, cells were fixed after staining for subsequent flow analysis.

### Individual recruitment assays with lentiviral delivery of dCas9 fusions to target reporters

Experiments were performed with dCas9 and sgTetO targeting the minCMV (pDY32) or pEF1α (pJT039) reporter in K562 cells. First, the TetO sgRNA was delivered by lentivirus and the cells were selected with puromycin. Then, dCas9 fusions (pJT216, 241, 244, 252, 258-260, 282, 300, 331, 336 and 342) were delivered by spinfection into 100k cells using 1 mL of unconcentrated lentivirus or virus that was concentrated 5x with LentiX Concentrator (Takara). Blasticidin was added at day 3 or 5 to select for dCas9 delivery. 2-9 days after delivery, the cells were analyzed by flow cytometry (BioRad ZE5 or Attune).

### Individual recruitment assays with lentiviral delivery of dCas9-activators in K562 cells

An experiment was performed targeting combinations of N, F, and Z activators and VP64 and VPR controls to CD2 with dCas9 in K562 cells. First, the CD2 sgRNAs 717 or 718 were delivered by lentivirus and the cells were selected with puromycin. Then dCas9 fusions (pJT252-260) were delivered by spinfection into 100k cells using 1 mL of lentivirus that was concentrated 5x with LentiX Concentrator (Takara). 4 days later, the cells were stained for CD2 expression (using the staining method described above) and analyzed by flow cytometry (BioRad ZE5).

Experiments were performed comparing dCas9-NFZ and VPR fusions targeting CD2, CD20, CD28 surface marker genes in K562 cells. sgRNAs, cloned into pMCB306, were first installed by lentiviral delivery and puromycin selection. Then, in one experiment, dCas9 plasmids were electroporated. Two days later, cells were immunostained for CD2 (APC), CD20 (APC), or CD28 (PE) expression and analyzed by flow cytometry after gating for dCas9 (TagBFP) and sgRNA (GFP). In another experiment, those stable sgRNA-expressing cell lines were thawed and dCas9 fusions (pJT216 dCas9 only, 259 VP64, 260 VPR, 258 NFZ, and 282 NZF) were delivered by unconcentrated lentivirus, and then delivery was measured by flow cytometry for BFP 3 days later (Attune). In another experiment, those sgRNA-expressing cell lines were infected with 5x concentrated lentivirus (LentiX Concentrator, Takara) for the same dCas9 fusions. Blasticidin selection was started 5 days later, and at day 9 post-infection the cells were stained and analyzed by flow cytometry (Attune).

dCas9-QNZF were delivered to K562 cells by lentivirus and selected for with blasticidin, then sgRNAs were delivered by lentivirus and selected for with puromycin, then 8 days after sgRNA delivery the cells were stained for the targeted surface marker genes and measured by flow cytometry (BioRad ZE5).

### Individual recruitment assay with transient electroporation of dCas9 in K562 cells

K562 cell lines stably expressing sgRNAs targeting CD2 (sg39, sg42, sg46, sg89), CD20 (sg135, sg148, sg275), or CD28 (sg07, sg56, sg94) were generated using lentivirus and puromycin selection until >90% of cells were GFP^+^. 1e6 cells were electroporated using 500 ng of dCas9-effector. After 2 days, cells were stained for surface marker expression with APC-conjugated antibody. Cellular fluorescence distributions were measured with the ZE5 flow cytometer (BioRad).

### Individual recruitment assay with transient electroporation of dCas12a in K562 cells

1e6 K562 cells were electroporated using 1000 ng of dCas12a-effector and 1000 ng of gRNA plasmid. Safe control gRNA was pKL03 and CD2_g1 gRNA was used to target CD2. After 3 days, cells were stained for CD2 expression with APC-conjugated antibody. Cellular fluorescence distributions were measured with the Attune NxT flow cytometer (Thermo Fisher). mCherry-APC bleedthrough compensation was applied and data was gated for high gRNA delivery (mCherry).

### Individual recruitment assays with lentiviral delivery of dCas12a fusions

Individual repressor domains were cloned into the pAR010 pEF-dEnAsCas12a-1xNLS-3xHA-Effector-T2A-BlastR-WPRE lentiviral vector, and delivered to K562 cells by spinfection. After transduction, the K562 cells were selected with 50 μg/ml blasticidin for 20 days. Then, guides with mCherry-puromycin selection marker targeting the CD32 or CD43 genes or a safe-targeting negative control were delivered by lentivirus to the cell lines. 2 days after transduction, these cells were selected with 1 μg/ml puromycin for 6 days. The cells were then recovered in fresh media for 1 day. After recovery, the cells were stained with APC conjugated surface marker antibodies as described above, along with HA antibody for gating on dCas12a-repressor expressing cells. The repression of the surface marker protein was then quantified by flow cytometry (Attune).

Individual activator domains were cloned into the pAR010 vector and delivered to K562 cells by spinfection. After transduction, the K562 cells were selected with 10 μg/ml blasticidin for 8 days. Then, guides with mCherry-puromycin selection marker targeting the CD2 TSS or a safe-targeting negative control were delivered by lentivirus to the cell lines. 3 days after transduction, these cells were selected with 1 μg/ml puromycin and 10 μg/ml blasticidin for 5 days. Then, the cells were permeabilized and stained with APC-conjugated CD2 antibody, along with HA antibody for gating on dCas12a-activator expressing cells. CD2 was then quantified by flow cytometry with gates for guide and dCas12a delivery (Attune).

### Individual recruitment assays with dCasMINI

The dCasMINI constructs were cloned using a previously published dCasMINI cassette (Xu et al., 2021). The sgRNA constructs used in GFP activation assays were generated previously (Xu et al., 2021), as were the HEK293T cells with an integrated GFP reporter (Gao et al., 2016). This HEK293T reporter line was generated by transducing cells with lentivirus containing EGFP expressed from a pTRE3G (Clontech) promoter. Pure populations of reporter cells were bulk sorted by fluorescence activated cell sorting (FACS) using a BD FACS Aria2 for stable fluorescent marker expression. For sorting GFP+ cells, 1 μg/mL doxycycline was added, and cells were transfected with a transactivator rtTA plasmid (Clontech). After sorting, the cells were cultured in the absence of doxycycline.

For transient transfections, cells were seeded in 24-well plates with 50,000 cells per well the day before transfection. 500 ng of dCasMINI constructs and 250 ng of sgRNA constructs were transfected to the reporter cell line using 2.5 uL of TransIT-LT1 (Mirus) in 100 ul of Opti-MEM reduced serum media (Thermo Fisher).

GFP expression was analyzed using CytoFLEX S flow cytometer (Beckman Coulter) 2 days posttransfection. Transfected cells were dissociated with 0.05% trypsin EDTA (Life Technologies) and resuspended in DPBS with 5% FBS for analysis. 15,000 cells were collected and analyzed from the population containing dCasMINI and its sgRNA constructs (mCherry-and BFP-positive after applying gates based on the non-transfected control).

### Individual recruitment assay with dCas9-activators in J774 cells

First, J774 cells were infected with the lentiviral dCas9-NFZ with blasticidin resistance and a BFP marker. 1e6 cells were incubated in 3 mL of 1x virus for 3 days then allowed to recover for 1 day, and then selected with 1 μg/ml blasticidin for 7 days or until >95% of cells were BFP^+^. sgRNA sequences were cloned into the pRAK127 backbone (which contains puromycin resistance and EGFP) at the BbsI site. 1e6 J774 cells with stably expressing dCas9-NFZ construct were then infected with 3 mL of 1x sgRNA lentivirus for 3 days before they were allowed to recover for 1 day, and were then selected with 1 μg/ml puromycin for 3 days or until >95% of cells were GFP^+^.

For qPCR, cells were harvested by exchanging media when cells reached ∼90% confluency, incubating for 24 h, and scraping. Cells were counted and 1e6 cells for each sample were collected and washed with PBS before RNA were extracted by QuickExtract RNA Extraction Solution (Lucigen, QER090150). SuperScript IV VILO Master Mix (Thermo Scientific, 11756050) was used to generate cDNA. qPCR were set up with three technical replicates for each sample with corresponding mouse Tacman gene expression probe (Taqman Probe ID: Mm01333821_m1 FAM dye for Actc1, Mm02620530_s1 FAM dye for Gpr84, Mm99999915_g1 VIC dye for Gapdh). All qPCR reactions were run on a BioRad CFX96™ Real-Time PCR Detection System.

For flow cytometry, 1e6 cells were similarly harvested and washed twice with PBS + 10% heat-inactivated FBS. Cells were then incubated in staining buffer with APC anti-mouse Cd2 Antibody (BioLegend 100112, 1:100 from the stock concentration) for 1 hour at 4°C. Cells were washed three times with PBS + 10% heat-inactivated FBS then analyzed with an Attune NxT Flow Cytometer (ThermoFisher).

### Western blot for effector fusion expression

K562 cells were transduced with a lentiviral vector containing a dCas9-3XFLAG-effector-T2A-BFP-BSD and 3 days later selected with blasticidin (10 μg/mL) for 15 days until complete. >80% of the cells appeared BFP^+^ by microscopy. 5-10 million cells were lysed in lysis buffer (1% Triton X-100, 150mM NaCl, 50mM Tris pH 7.5, Protease inhibitor cocktail). Protein amounts were quantified using the Pierce BCA Protein Assay kit (Bio-Rad). Equal amounts were loaded onto a gel and transferred to a PVDF membrane. Membrane was probed using FLAG M2 monoclonal antibody (1:1000, mouse, Sigma-Aldrich, F1804) and beta-actin antibody (1:1000, Rabbit, Abcam, ab8227) as primary antibodies. Anti-rabbit 680 (IRDye® 680LT Donkey anti-Rabbit IgG Secondary Antibody, 926-68023, Li-Cor) and anti-mouse 800 (IRDye® 800CW Goat anti-Mouse IgG Secondary Antibody, 926-32210, Li-Cor) at 1:10000 dilution were used as secondary antibodies. Blots were imaged on a Li-Cor Odyssey CLx. Band intensities were quantified using ImageJ (Rueden et al., 2017).

### Chromatin modification mapping

Stable K562 cell lines expressing both the dCas9-repressor and CD43 sg15 were generated with lentiviral delivery and selected with antibiotics as described above for the recruitment assay. After being frozen, these cells were thawed and sorted to generate a pure population based on fluorescence of the BFP marker for dCas9 and mCherry for sgRNA (Sony SH800S sorter), although both markers were already high due to selection. Profiling of chromatin modifications at the CD43 endogenous gene in K562 cells was performed using the EpiCypher CUTANA ChIC/CUT&RUN kit (EpiCypher, 14-1048), according to manufacturer instructions (Kit v3). The following conditions were used: an input of ∼400,000 cells per antibody condition and an overnight antibody incubation at 4°C with H3K4me3 (0.5 μg/ml, EpiCypher), H3K9me3 (0.9 μg/ml, Diagenode C15410193), H3K27me3 (1:50, Cell Signaling Technology 9733S), H2AK119Ub (1:100, Cell Signaling Technology 8240S), and IgG (0.5 μg/ml, EpiCypher) antibodies. Dual-indexed sequencing libraries were prepared using NEBNext Ultra II DNA Library Prep Kit for Illumina (New England Biolabs) with 10 ng or 10 uL of input DNA (whichever was lower volume). DNA size selection was performed with SPRIselect (Beckman Coulter). Library concentrations were quantified with the Qubit 1x dsDNA HS Assay Kit (ThermoFisher), and library fragment sizes were assessed with an Agilent TapeStation System, then sequenced with paired end 2×150 cycle reads on an Illumina HiSeq X (Admera Health). Each sample had ∼16e6 paired end reads. Sequencing reads were trimmed with Trimmomatic, aligned to the human GRCh38 reference genome with Bowtie2, and filtered to remove duplicates with Picard and Samtools. RPKM normalized coverage data was visualized with Integrative Genomics Viewer (J. T. Robinson et al., 2011).

### Rapamycin-induced expression of reporter

An experiment was performed where plasmids with the rapamycin inducible circuit and the citrine reporter gene were delivered to K562 cells. 2 ug of Rapamycin inducible constructs containing the activators NZF, NFZ, S3H and no effector control were electroporated using Lonza 2b nucleofector per 300k K562 cells, resuspend in 1 mL RPMI in 24 Well plate. On day 1 post electroporation, cells were split in 2 wells, in one well 100 nM Rapamycin (LC Laboratories) was added and the other well contained regular RPMI complete media. On day 2 post electroporation,

Experiments were performed where plasmids with the rapamycin inducible circuit and the citrine reporter gene were delivered to HEK293T cells. The day prior to transfection, 500k HEK293T cells were plated in a 6– well plate. 3 ug of plasmids were transfected into HEK293T cells using PEI transfection reagent. One day later, cells were split and left untreated or 10 nM rapamycin was added, and two days later citrine mean fluorescence intensity (MFI) was measured by flow cytometry on the Attune cytometer.

### ELISA for HGF concentrations

HEK293T cells were transfected using Lipofectamine LTX (Invitrogen) with 1000 ng plasmid containing inducible HGF (Addgene #188749) or a constitutive pEF1α-HGF (Addgene #188748) plasmid as a control. One day later the cells were treated with varying doses of rapamycin for two days. Then, the tissue culture medium was collected in a microcentrifuge tube and centrifuged at 1400 RPM for 5 mins. Supernatant was transferred to a clean microcentrifuge tube and stored at –80°C until ready for use in enzyme-linked immunosorbent assay (ELISA). Protein concentration of human HGF was determined by ELISA using the Quantikine Human HGF Immunoassay (R&D Systems #DHG00B) according to manufacturer instructions and analyzed by a SpectraMax ID3 microplate reader. The dose response curve was fitted using GraphPad Prism’s built-in nonlinear regression equation called “log(agonist) vs. response – Variable slope”, which is defined as: Y = Bottom + (Top – Bottom) / (1 + 10^((LogEC50 – X) * HillSlope)). EC50 is the concentration of rapamycin agonist that gives a response halfway between Bottom and Top. HillSlope describes the steepness of the curve. Top and Bottom are plateaus in the units of the Y axis (i.e. HGF concentration).

### Analysis software for flow cytometry data

The flow cytometry data were analyzed with the MATLAB program EasyFlow (https://antebilab.github.io/easyflow/) or the Python program Cytoflow (https://cytoflow.github.io/).

### Statistical analyses

Non-linear regression, Kruskal-Wallis test and two-way ANOVA were performed in Prism 9.3.1. Other statistical analyses were performed in Python using SciPy (Virtanen et al., 2020). The statistical tests used were two-sided (where applicable) and are indicated in the text and/or figure legends. The “n” for each analysis is indicated in the main text or in figure legends of relevant analyses. Significance was set at p<0.05. No methods were used to determine whether the data met assumptions of the statistical approach.

### Data availability

All Illumina sequencing data generated in this study will be deposited in the NCBI GEO database.

### Code availability

Code to analyze HT-recruit domain screens (https://github.com/bintulab/HT-recruit-Analyze) and sgRNA screens (https://github.com/elifesciences-publications/dmorgens-castle) is available online.

### Materials availability

Plasmids generated in this study will be made available from Addgene.

## Acknowledgements

We thank Raeline Valbuena and other members of the Bassik and Bintu labs for helpful conversations and assistance. J.T. is supported by the F99/K00 Fellowship of the National Institutes of Health (NIH-1F99DK126120-01; NIH-4K00DK126120-03). M.C.B. is supported by a grant from Stanford ChEM-H and an NIH Director’s New Innovator Award (1DP2HD08406901). This work was supported by grants from NIH/ENCODE 5UM1HG009436-02 (M.C.B.), NIH/NIGMS R35M128947 (L.B.), and NIH/NHGRI 5R01HG011866 (M.C.B. and L.B.). M.V.V. was supported by NIH T32 Training Grant (T32GM007276). N.D. was supported by grants from NSF GRFP DGE-1656518 and ARCS Foundation. X.X. was supported by Chan Zuckerberg Initiative Neurodegeneration Collaborative Pairs Phase 2. L.S.Q. is a Chan Zuckerberg Biohub – San Francisco Investigator.

## Author contributions

J.T., M.C.B., and L.B. designed the overall study. J.T., M.V.V., A.R., and C.L. generated reporter cell lines. J.T. and K.S. cloned pooled libraries. J.T., M.V.V., and A.R. performed HT-recruit and other high-throughput experiments, and individual validations. G.H. designed and tested sgRNAs. A.R. and K.L. designed and performed dCas12 experiments. J.T., D.Y., N.D., A.X.M., and P.H.S. designed oligonucleotide libraries. P.H.S. characterized homeodomain mutants. J.T., A.R., and D.Y. performed CRISPRi benchmarking screens. M.G. and R.A.K. performed experiments with J774 cells and K.L. with MCF10a cells. X.X. designed and performed dCasMINI experiments with support from L.S.Q.. J.T. and M.V.V. analyzed the data and prepared the figures, with additional analysis of the tiling library by N.D.. J.T., M.V.V., L.B., and M.C.B. wrote the manuscript with input from all authors.

## Supplementary Text

### Combinations of Repressors

Non-KRAB repressors have been used for gene silencing and epigenome editing (Liu et al., 2016; O’Geen et al., 2017), but these have generally not demonstrated a similar efficiency as KRAB for CRISPRi applications. In our HT-recruit screens, we found that non-KRAB repressors were less robust across contexts (**Figure 1D and Figure 2F**). Accordingly, in individual assays, when fused to dCas9, many repressor domains that are strong when recruited with rTetR in K562, including RYBP and MGA (Tycko et al., 2020), were much weaker than KRAB when targeted to CD43 in K562 cells (**Supplementary Figure 13A**), or to a reporter in HEK293T cells (**Supplementary Figure 13B**). These results agreed with data on RYBP_RYBP from the HT-recruit screens (**Figure 2F and Supplementary Figure 9A**) and MGA was not included in the Pfam library. One mixed case was SUMO3, which silenced CD43 with KRAB-like efficiency using the guide sg15, but it did not silence with sg10, which was also the trend for SUMO3 in the dCas9 HT-recruit screens with these two guides (**Supplementary Figure 7I and Supplementary Figure 13A**). Despite these initial difficulties, we knew these repressors could silence in some contexts (e.g. on rTetR) and that they depended on different co-repressors than KRAB (e.g. not KAP1), which motivated further efforts to develop them as tools.

Others have reported enhancement of CRISPRi by combining multiple repressors with ZNF10 KRAB (Nuñez et al., 2021; Yeo et al., 2018) so we also tested combinations of these non-KRAB repressors. First, when we combined MGA[381-390] and MGA[2431-2460] together we observed super-additive repression (23% versus 0% and 10% individually) at the HEK293T reporter (**Supplementary Figure 13B**). However, adding this MGA combination and/or RYBP to KRAB did not increase silencing at endogenous CD43 and CD81 in K562 cells or at the HEK293T reporter (**Supplementary Figure 13C–E**). We used CUT&RUN to measure histone modifications at the target CD43 locus, and found similar modifications mediated by the KRAB[WSR7EEE] repressor alone and the KRAB[WSR7EEE]+RYBP+MGA1+2 combination (**Supplementary Figure 13F**). Specifically, there was a local deposition of heterochromatic mark H3K9me3 in the 5 kb region around the target, while the activation-associated H3K4me3 mark was locally ablated and partially reduced at a distal site 16 kb away. While previous literature associates MGA with polycomb chromatin modifications (Stielow et al., 2018), we observed no changes for those H3K27me3 or H2AK119Ub marks. Moreover, these repressor combinations have the same epigenetic memory as ZNF10 KRAB when targeting the HEK293T reporter (**Supplementary Figure 13G**), consistent with RYBP-MGA mediating similar chromatin modifications as KRAB.

We further benchmarked non-KRAB repressor combinations in a CRISPRi screen targeting promoters that were previously shown to be essential with a ZNF10 KRAB screen (Gilbert et al., 2014). Here, KRAB+RYBP+MGA showed somewhat lower efficiency than KRAB alone (**Supplementary Figure 13H**). Across the experiments here, only MGA[381-390] plus MGA[2431-2460] showed synergy. When added to KRAB, all second repressors were neutral or somewhat reduced silencing strength, consistent with previous observations that repressor fusions are susceptible to steric blocking of activity (O’Geen et al., 2017). These results are also concordant with our recent HT-recruit screen of effector combinations which showed very few repressors in the Pfam library can enhance silencing when concatenated with KRAB; GSX2 homeodomain was one exception (Mukund et al., 2022).

### Combinations of Activators

We identified NZF as a potent tripartite activator. To test further domain concatenation, we fused another short activator domain, SMARCA2 QLQ, to NZF and found the resulting QNZF was a stronger activator at the CD2 gene and minCMV reporter, but this improvement did not hold at CD20 or CD28, or in HEK293T cells. This was consistent with QLQ activating CD2 and minCMV but not CD20 or CD28, and being stronger in K562 than HEK293T cells in the HT-recruit screens (**Figure 4A and Supplementary Figure 15L–N**). In addition, we found the Q, N, and F (but not Z) domains can synergistically activate CD2 when fused as homotypic bipartite combinations (**Supplementary Figure 15O**). However, heterotypic combinations lack repetitive sequences and should be simpler to synthesize and package in viral vectors, so we selected NFZ for further study.

